# Intra-host genomic variation of *Haemophilus influenzae* isolates from asymptomatic nasopharyngeal carriers involves genes encoding proteins with diverse inferred functions

**DOI:** 10.1101/2025.07.03.663008

**Authors:** Randall J. Olsen, S. Wesley Long, Yuvanesh Vedaraju, Sandra Tomasdottir, Helga Erlendsdottir, Ásgeir Haraldsson, Karl G. Kristinsson, James M. Musser, Gunnsteinn Haraldsson

## Abstract

*Haemophilus influenzae* is a human-specific pathogen that causes infections ranging in severity from otitis media to potentially fatal meningitis. It also asymptomatically colonizes the upper respiratory tract. Although intra-host genomic variation of *H. influenzae* has been investigated in some anatomic sites, the genes most frequently acquiring nonsynonymous (amino acid changing) or nonsense (protein truncating) single nucleotide polymorphisms (SNPs) during human carriage or infection remain largely unidentified. To study intra-host genomic variation of *H. influenzae* during human asymptomatic carriage in the nasopharynx, the genomes of 805 isolates recovered from 24 healthy Icelandic children were sequenced. Most children were colonized with isolates with a single multilocus sequence type (MLST), although some were concurrently colonized with isolates with multiple MLSTs. Intra-host genomic variation was discovered with 120 genes acquiring SNPs in at least one isolate. Among them, 69 genes were recurrently polymorphic in isolates recovered from multiple children, and 72 SNPs occurred in multiple isolates recovered from the same child. The polymorphic genes encode proteins with diverse inferred functions, including transcription regulators and putative virulence factors. Many of the proteins likely play roles in bacterial fitness, virulence, and host-pathogen molecular interactions. This intra-host variation study provides a model for understanding the genomic diversity acquired by *H. influenzae* during human asymptomatic carriage in the nasopharynx.

## INTRODUCTION

*Haemophilus influenzae*, a human-specific pathogen, is a Gram-negative coccobacillus historically recognized as a cause of serious infections ^1, 2^. Strains are serotyped based on expression of capsular polysaccharides (types a–f) ^2^. Before the introduction and broad distribution of a polysaccharide conjugate vaccine targeting *H. influenzae* type b (Hib), Hib was among the primary causes of life-threatening pediatric meningitis, bacteremia, pneumonia, and epiglottitis ^3, 4^. While infections caused by Hib and other encapsulated serotypes have markedly declined in populations with high vaccine uptake, unencapsulated (serologically nontypeable) strains have become more common ^5–7^. *H. influenzae* (particularly unencapsulated strains) also commonly colonizes the upper respiratory tract of young children and elderly adults ^2, 8–11^. Although asymptomatic colonization rates vary by age, race, sex, season, country, and environmental exposures, they are particularly high among individuals with chronic respiratory tract conditions and close person-to-person contacts such as daycare center attendees ^2, 12–18^.

Sequencing of bacterial genomes began in 1995 with *H. influenzae* strain Rd ^19^. Since then, researchers have sequenced the genomes of more than seventeen thousand *H. influenzae* strains to study population genomic structure ^20–24^. These studies have revealed new insights to genomic diversity, gene content, and gene polymorphisms, particularly among serologically nontypeable strains. However, intra-host genomic variation has been far-less studied, leaving many key questions unanswered ^21, 25–27^. Specifically, the genes that most frequently acquire nonsynonymous (amino acid-changing) or nonsense (protein-truncating) single nucleotide polymorphisms (SNPs) during human colonization or infection remain largely unidentified.

Studies of intra-host genomic variation with influenza A virus ^28^, SARS-CoV-2 ^28, 29^, *Klebsiella pneumoniae* ^30^, *Streptococcus pyogenes* ^31^, and other human pathogens ^32–37^ have provided key insights to microbial pathogenesis, including host-pathogen molecular interactions, strain fitness and virulence. To address this gap for *H. influenzae*, we recently investigated^38^ intra-host genomic variation in serologically nontypeable isolates obtained from ear drainage fluid collected from children with otitis media. The analysis demonstrated that intra-host genomic variation in the middle ear affects numerous genes encoding proteins with diverse inferred functions ^38^. To expand on this line of investigation, we sequenced isolates recovered from nasopharyngeal swabs of healthy children with asymptomatic carriage.

## Materials and Methods

### Specimen collection and *H. influenzae* culture

Since 2009, investigators at Landspítali - the National University Hospital of Iceland have conducted an annual surveillance study examining asymptomatic carriage of *H. influenzae*, *Streptococcus pneumoniae*, and *S. pyogenes* in the Reykjavik capital region ^39–41^. In March of each year, researchers visit 15 daycare centers, which are randomly assigned alphabetic designations to preserve anonymity (Table 1), located across the Reykjavík Capital Region, comprising Reykjavík and 6 neighboring municipalities. The Reykjavík Capital Region accounts for approximately 65% of the Icelandic population (01 May 2025, www.statice.is) ^42^. Parents provided informed consent for nasopharyngeal swab collection and completed a questionnaire on recent antibiotic use. Each year, one nasopharyngeal swab sample (eSwab, Copan Italia s.p.a., Brescia, Italy) is collected from approximately 500 healthy children aged 1 to 6 years for asymptomatic carriage analysis (Figure 1). In 2017 and 2018, 507 and 467 children were sampled, respectively. Swabs were immersed in Amies transport medium, and within 18 hours, swabs were plated on one blood agar plate and one chocolate agar plate with a bacitracin disc (Becton, Dickinson, and Company, Vaud, Switzerland) to isolate *H. influenzae*. Swabs were also plated on MacConkey, tryptic soy blood, nutrient, and SS agars to recover *S. pneumoniae* and *S. pyogenes* isolates not used in this study. Plates were incubated aerobically in the presence of 5% CO_2_ for 18–20 hours at 36°C. Within 6 hours of plating, swabs were frozen at −80°C without cryopreservative additives. *H. influenzae* was identified using its characteristic colony morphology, resistance to bacitracin, and proteome from MALDI-TOF mass spectrometry (Bruker Daltonics, Billerica, MA). Bacterial growth was semi-quantified on a scale of 0 to 3+ following standard clinical microbiology laboratory procedures ^43^. Children with nasopharyngeal swabs producing 3+ *H. influenzae* colonies were randomly assigned a numeric designation for anonymity (Table 1).

**Fig. 1.**
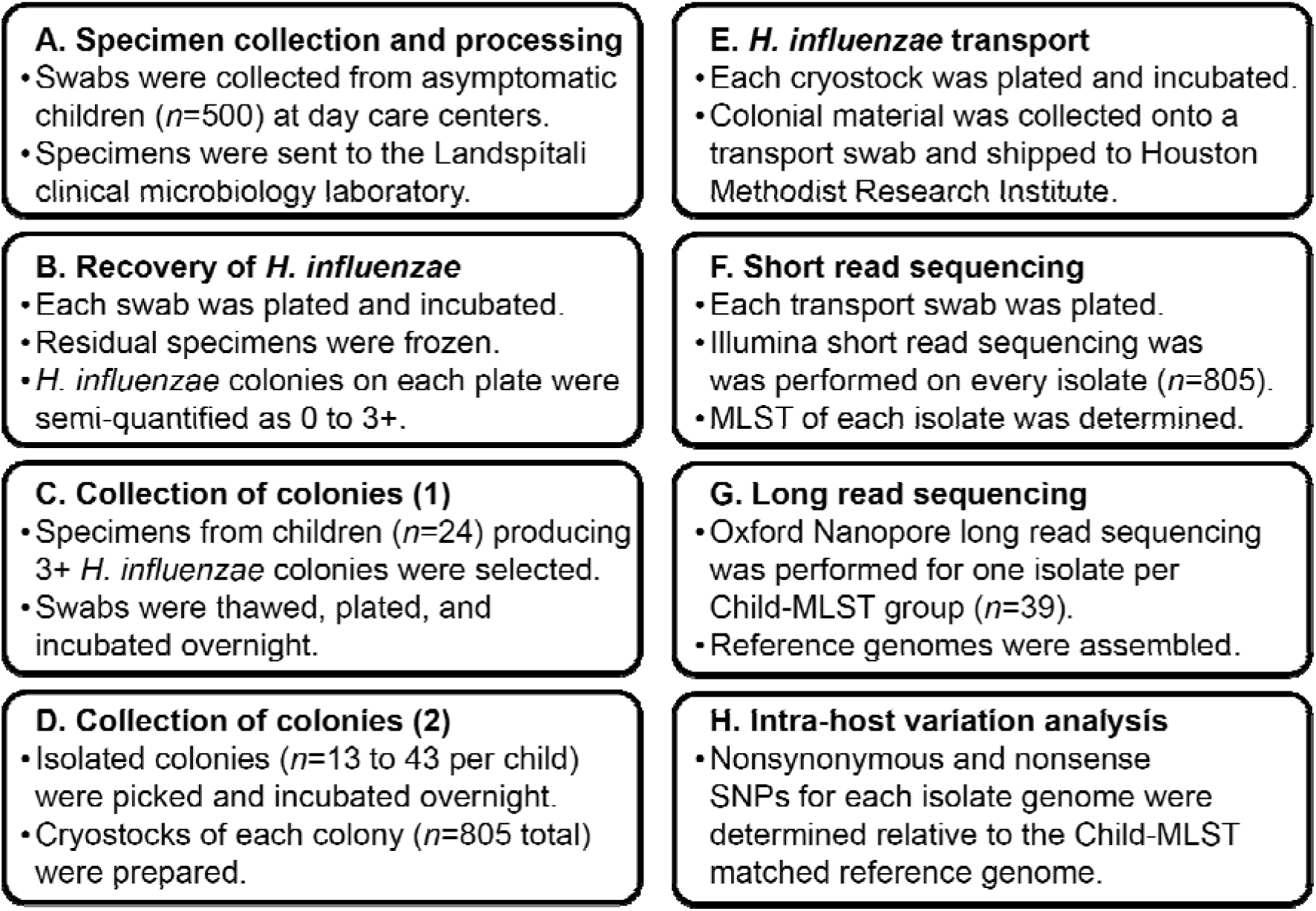
Specimen acquisition, processing and intra-host genomic variation analysis methods.

**Table 1.** Demographic information for the 24 children.

| **Child** | **Age (years)** | **Sex** | **Day Care Center** | **Year** | **Isolates (number)** |
| --- | --- | --- | --- | --- | --- |
| 1 | 4 | M | A | 2018 | 39 |
| 2 | 5 | F | B | 2018 | 25 |
| 7 | 6 | M | A | 2018 | 26 |
| 8 | 3 | M | B | 2018 | 34 |
| 10 | 5 | M | B | 2018 | 39 |
| 13 | 5 | M | A | 2018 | 34 |
| 15 | 3 | M | A | 2018 | 38 |
| 16 | 3 | F | C | 2018 | 32 |
| 20 | 6 | F | A | 2018 | 35 |
| 27 | 3 | F | D | 2017 | 26 |
| 27 | 4 | F | D | 2018 | 33 |
| 29 | 2 | F | E | 2017 | 23 |
| 29 | 3 | F | E | 2018 | 43 |
| 36 | 2 | F | A | 2018 | 17 |
| 39 | 2 | F | A | 2018 | 13 |
| 43 | 3 | F | F | 2018 | 35 |
| 46 | 1 | F | G | 2018 | 35 |
| 52 | 3 | F | H | 2018 | 33 |
| 56 | 2 | M | B | 2018 | 20 |
| 57 | 3 | M | B | 2018 | 35 |
| 59 | 3 | M | C | 2018 | 32 |
| 61 | 3 | M | I | 2018 | 31 |
| 64 | 3 | F | I | 2018 | 35 |
| 68 | 3 | M | D | 2018 | 29 |
| 73 | 2 | M | J | 2018 | 31 |
| 75 | 3 | M | K | 2018 | 32 |

### *H. influenzae* isolate recovery for intra-host variation analysis

For this study, nasopharyngeal swabs from 24 children collected in 2018 that produced 3+ *H. influenzae* colonies were selected for intra-host genomic variation analysis (Table 1). Two children (Child 27 and Child 29) had nasopharyngeal swabs available from 2017 that were also included. Swabs were removed from the freezer and plated on two chocolate agar plates that were incubated overnight at 36°C with 5% CO_2_ (Figure 1). All isolated colonies (*n*=13 to 43 colonies per swab; *n*=805 colonies total) were picked (Table S1), systematically labeled, subcultured, and confirmed as *H. influenzae* by MALDI-TOF mass spectrometry. Subsequently, a loopful of colonial material from each subculture, representing one colony recovered from the nasopharyngeal swab, was cryopreserved to create a stock. The stocks were later plated, incubated, collected onto a transport swab submerged in charcoal media (Medical Wire and Equipment Co Ltd, U.K.), and transferred to the Houston Methodist Research Institute for whole-genome sequencing (Figure 1).

### Whole-genome sequencing

The genomes of the 805 *H. influenzae* isolates were sequenced with well-established protocols ^38, 44–47^. Briefly, Illumina short-read sequencing was performed on all isolates. Bioinformatic analysis determined the multilocus sequence type (MLST) of each isolate (described below). To generate one closed reference genome for isolates of each MLST from each child, Oxford Nanopore Technology long-read sequencing was also performed (*n*=39 genomes, Table 2). For both sequencing technologies, isolates were incubated overnight at 37°C in the presence of 5% CO_2_ on chocolate agar, with DNeasy Blood and Tissue kits (Qiagen, Germantown, MD) used for genomic DNA extraction. Nextera XT kits and a NextSeq 550 instrument (Illumina, San Diego, CA) were used for short-read sequencing. V14 Native Barcoding Kits, R10.4 flow cells, and a GridION instrument were used for long-read sequencing (Oxford Nanopore Technologies, OX4 4DQ, UK).

**Table 2.** ***H. influenzae* asymptomatic carriage reference genomes.**

| **Isolate** | **Child** | **MLST** | **Genome (bp)** | **Genes (number)** |
| --- | --- | --- | --- | --- |
| MHI3301 | 1 | 266 | 1,942,768 | 1,964 |
| MHI3331 | 2 | 1040 | 1,954,682 | 1,938 |
| MHI3332 | 2 | 544 | 1,896,225 | 1,907 |
| MHI3338 | 2 | 1191 | 1,807,729 | 1,774 |
| MHI3341 | 2 | 652 | 1,801,476 | 1,770 |
| MHI3361 | 16 | 395 | 1,858,256 | 1,843 |
| MHI3391 | 27 | 2954 | 1,874,458 | 1,880 |
| MHI3401 | 27 | 494 | 1,873,966 | 1,880 |
| MHI3421 | 29 | 266 | 1,915,076 | 1,935 |
| MHI3428 | 29 | 1030 | 1,867,825 | 1,851 |
| MHI3451 | 43 | 836 | 1,837,652 | 1,918 |
| MHI3481 | 46 | 266 | 1,943,360 | 1,966 |
| MHI3511 | 27 | 1034 | 1,804,984 | 1,798 |
| MHI3512 | 27 | 683 | 1,927,999 | 1,961 |
| MHI3542 | 29 | 199 | 1,842,476 | 1,828 |
| MHI3543 | 29 | 598 | 1,834,549 | 1,814 |
| MHI3546 | 29 | 536 | 1,890,480 | 1,875 |
| MHI3578 | 7 | 159 | 1,815,639 | 1,780 |
| MHI3620 | 52 | 105 | 1,982,422 | 2,023 |
| MHI3652 | 61 | 658 | 1,958,690 | 1,980 |
| MHI3655 | 61 | 1012 | 1,889,727 | 1,889 |
| MHI3685 | 73 | 1030 | 1,866,203 | 1,847 |
| MHI3716 | 75 | 544 | 1,933,379 | 1,960 |
| MHI3941 | 8 | 431 | 1,842,302 | 1,898 |
| MHI3963 | 8 | 2955 | 1,901,096 | 1,909 |
| MHI3976 | 10 | 57 | 1,960,738 | 2,002 |
| MHI4015 | 13 | 2156 | 1,971,532 | 1,994 |
| MHI4050 | 15 | 2956 | 1,960,255 | 1,924 |
| MHI4051 | 15 | 2957 | 1,862,158 | 1,863 |
| MHI4060 | 15 | 1238 | 1,957,679 | 1,983 |
| MHI4090 | 20 | 946 | 1,934,415 | 1,967 |
| MHI4125 | 36 | 36 | 1,848,113 | 1,850 |
| MHI4160 | 39 | 159 | 1,815,725 | 1,781 |
| MHI4194 | 56 | 334 | 1,860,596 | 1,846 |
| MHI4227 | 57 | 161 | 1,981,786 | 2,002 |
| MHI4262 | 59 | 1409 | 1,775,912 | 1,768 |
| MHI4264 | 59 | 280 | 2,022,617 | 2,079 |
| MHI4297 | 64 | 2958 | 1,815,031 | 1,785 |
| MHI4332 | 68 | 264 | 1,927,212 | 1,943 |

### Whole-genome sequence analysis

Whole-genome sequence analysis was performed using established protocols ^38^. Briefly, Trimmomatic, Musket, and Fastq-pair were used for quality trimming, adaptor decontamination, and read pairing ^48–50^. SRST2 and custom database files were used to assess gene content, including alleles for multilocus sequence typing, capsule genotyping, and fucose operon genotyping ^51^. SRST2 and PlasmidFinder.fasta and ARGannot.r1.fasta database files were used to identify plasmid replicons and antimicrobial resistance genes ^51^. Bifrost and Corer were used to determine the *H. influenzae* core genome from 99 genome sequences (Table S2) deposited in the NCBI Microbial Genome Database (accessed 05/01/2025) ^52, 53^. MUMmer dna-diff was used to determine phylogenetic relationships among the 805 *H. influenzae* asymptomatic carriage isolates relative to the core genome ^54^. Rapidnj, SplitsTree, and Dendroscope were used to generate and visualize phylogenetic trees ^55^.

### Reference genome assembly, closure, and annotation

Oxford nanopore long reads were generated for one isolate representing each Child-MLST group (Table 2). FastQC was used to assess read length and sequence quality, and MinKNOW (v.24.11.8) with high accuracy parameters was used to call nucleotides (Oxford Nanopore Technologies). Reference genomes were hybrid-assembled from the long- and short-read sequences. Filtlong (v.0.2.1), applying a minimum read length of 1 kb length and a retention threshold of 90%, was used to filter the long-read FASTQ files ^56^. Short-read FASTQ files were trimmed and filtered using Trimmomatic (v.0.39) ^48^. Hybrid genome assembly was initially conducted with Unicycler using the “normal” setting (v.0.5.0), SPAdes (v.3.15.4), and Racon (v.1.5.0) ^57–59^. Genomes that did not assemble to closure were reattempted using the Unicycler “bold” setting. Genomes that did not assemble with Unicycler were reassembled using Trycycler in combination with Minimap2 (v.2.17-r941), Miniasm (v.0.3-r179), Raven (v.1.8.1), Minipolish, Polypolish, and POLCA ^46^. PROKKA with default settings was used to annotate the closed circularized reference genomes ^60^. Artemis (v.18.2.0) was used for genome visualization ^61^.

### Intra-host genomic variation determination

Contigs were assembled from the trimmed, error-corrected, paired short-reads using SPAdes ^57^. MUMmer dna-diff was used to determine polymorphisms relative to the Child-MLST-matched reference genome ^54^. An exclusion file for each closed circularized reference genome was created by identifying mismapped short reads to itself. Genes encoding transfer RNAs, ribosomal proteins, and pilis proteins were excluded due to their highly repetitive coding regions. Hypothetical genes were also excluded since they have uncertain inferred functions. Prephix, Phrecon, pre2snpfx, and snpfx in-house scripts were used for polymorphism data processing as previously described ^30^. MEGA (v.11) was used to determine pairwise distances between strains ^62^. Rapidnj, SplitsTree, and Dendroscope were used to generate and visualize phylogenetic trees ^55^. Annotated genes with nonsynonymous (amino acid-changing) or nonsense (protein-truncating) SNPs compared to the reference genome underwent manual curation. UniProt and literature review were used to assign an inferred function to the protein encoded by each polymorphic gene ^63^. Prism (v.10.4.2) was used for graphing and statistical analysis (GraphPad Software, Boston, MA).

## RESULTS

### Gene content analysis

To assess intra-host genomic variation during human asymptomatic carriage, whole-genome sequencing of 805 *H. influenzae* isolates recovered from nasopharyngeal swabs collected in 2018 from 24 healthy Icelandic children (*n*=13 to 43 isolates per child) was performed (Figure 1 and Table 1). Samples collected in 2017 were also available for two children (child 27 and child 29). First, to assess genetic relationships among the asymptomatic carriage isolates, MLSTs were determined. In total, 39 distinct child-MLST groups were identified (Table 2 and S1). While 17 children were colonized with isolates with a single MLST, 7 children were concurrently colonized with isolates with multiple MLSTs. Child 27 and child 29 were colonized with isolates from multiple different MLSTs during 2017 and 2018 (Table S1). In most instances, isolates with different MLSTs differed at multiple non-contiguous loci, suggesting concurrent asymptomatic carriage of distantly related lineages (Table S2). Additionally, isolates from four MLSTs were carried by multiple children (Table 2, S1, and S2). The clonal complexes containing the 39 MLSTs are known causes of human infection ^64^.

Gene content across the 805 *H. influenzae* asymptomatic carriage isolates was analyzed (Table S1). First, the presence of the fucose (*fuc*) operon was assessed, because previous reports indicate that *fucK* deletion occurs in some *H. influenzae* isolates and may lead to an undefined MLST ^65, 66^. All isolates had an intact *fuc* operon, except MLST2956 isolates from child 15 (Table S1). Second, genes associated with capsule production, plasmid replication, and antimicrobial resistance were identified. All isolates were unencapsulated (nontypeable) and did not contain plasmid replicons (Table S1). Although most isolates lacked antimicrobial resistance genes, a TEM-1D beta-lactamase was present in isolates recovered from child 2 (1 of 1 MLST652 isolates and 12 of 13 MLST1040 isolates), child 27 (all 14 MLST683 isolates), and child 59 (3 of 11 MLST280 isolates and 1 of 21 MLST1409 isolates). Notably, parental surveys indicated no recent antibiotic treatment for the three children with isolates with a TEM-1D beta-lactamase.

### Phylogenetic analysis

To place the 805 *H. influenzae* asymptomatic carrier isolates into a broader phylogenetic context, a phylogenetic tree using SNPs determined relative to the *H. influenzae* core genome was constructed (Table S3). This analysis revealed two key findings. First, the 39 child-MLST groups were genomically diverse, differing at 192,419 unique SNP loci, representing 10.3% of the nucleotides in the core genome (Figure 2). Second, isolates with the same MLST from different children formed closely related yet distinct subpopulations. For example, isolates classified as MLST159 from child 7 and child 39 were closely aligned relative to isolates from other MLSTs (Figure 2) but formed distinct subpopulations (Figure 3). Subsequent intra-host variation analyses for isolates from each child-MLST group were performed independently due to this high genomic diversity and evidence of distinct molecular evolutionary pathways.

**Fig. 2.**
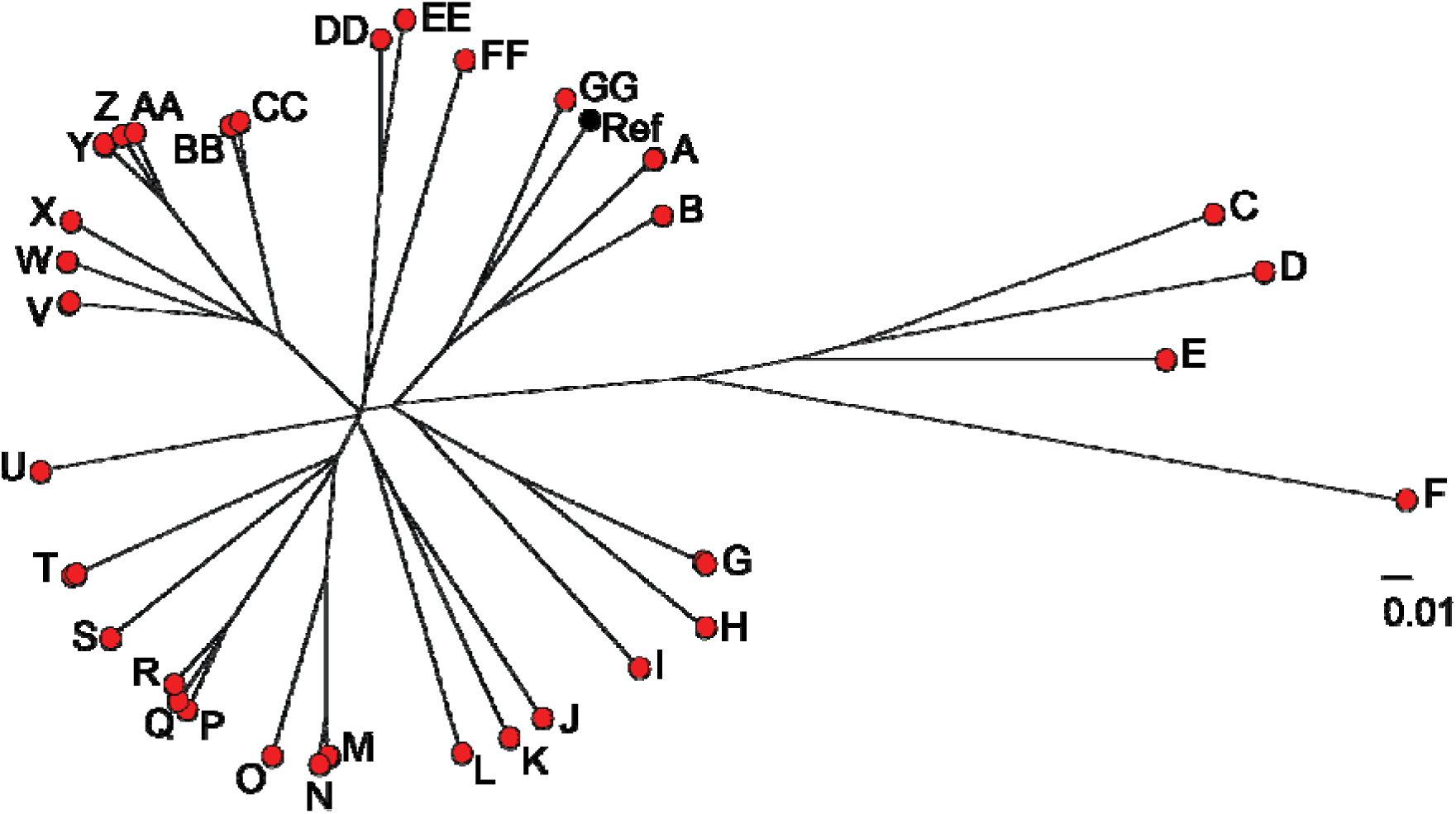
Phylogeny of *H. influenzae* isolates. Phylogeny was inferred by neighbor-joining based on SNPs identified by Illumina short-read whole-genome sequencing of the 39 closed circularized reference genomes with read mapping relative to the *H. influenzae* core genome. Isolates (A) MHI3391 and MHI3401, (B) MHI3511, (C) MHI3512, (D) MHI3543, (E) MHI3338, (F) MHI4050, (G) MHI3578 and MHI4160, (H) MHI3361, (I) MHI3976, (J) MHI4015, (K) MHI4227, (L) MHI4194, (M) MHI3301 and MHI3481, (N) MHI3421, (O) MHI4090, (P) MHI3652, (Q) MHI4264, (R) MHI3620, (S) MHI3655, (T) MHI3428 and MHI3685, (U) MHI4051, (V) MHI3941 and MHI3963, (W) MHI3546, (X) MHI4060, (Y) MHI3332 and MHI3716, (Z) MHI4262, (AA) MHI4297, (BB) MHI3331, (CC) MHI4332, (DD) MHI3341, (EE) MHI3542, (FF) MHI4125, and (GG) MHI3451 are shown. Genomically closely related isolates, such as the same MLST or a single locus variant, may appear as overlapping circles.

**Fig. 3.**
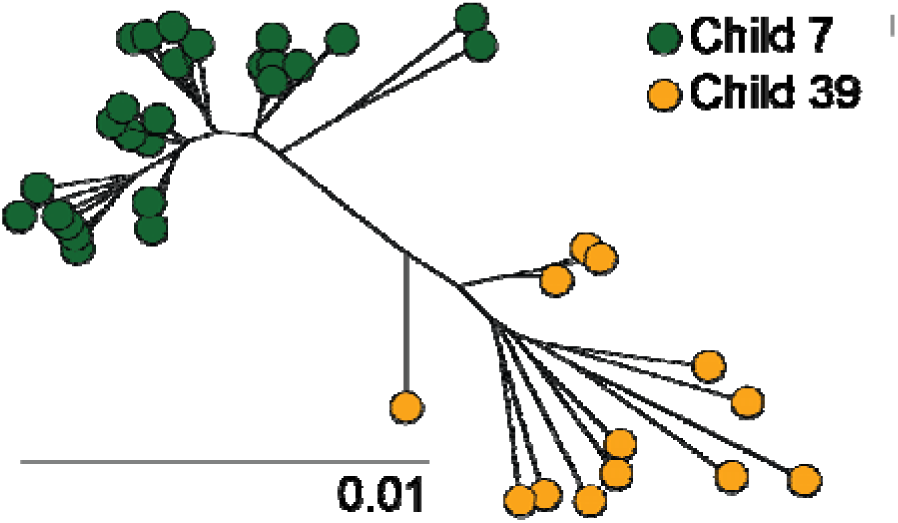
Phylogeny of MLST159 *H. influenzae* isolates recovered from child 7 and child 39. Phylogeny was inferred by neighbor-joining based on SNPs identified by Illumina short-read whole-genome sequencing with read mapping relative to the *H. influenzae* core genome.

### Intra-host genetic variation analysis

To identify *H. influenzae* genes undergoing intra-host genomic variation within each child’s nasopharynx, a closed circularized reference genome for each child-MLST group was assembled (Table 2). A comparable strategy has been effectively used in prior studies of closely related organisms ^30, 38, 45, 46^, including intra-host genomic variation analysis of *H. influenzae* isolates recovered from otitis media patients ^38^. Using clonally related reference genomes allows for highly accurate SNP determination ^67^. The resulting 39 closed circularized reference genomes ranged from 1,775,912 to 2,022,617 base pairs and included 1,768 to 2,079 genes (Table 2), which is consistent with the genome sizes and gene counts of other publicly available *H. influenzae* genomes (Table S2).

For each *H. influenzae* asymptomatic carrier isolate, SNPs were determined relative to the child-MLST-matched reference genome. A total of 120 genes (representing 6.3% of the mean 1,897 genes in the reference genomes) acquired nonsynonymous (amino acid-changing) or nonsense (protein-truncating) SNPs in at least one isolate (Table S4). No gene deletions or frameshift mutations were identified. Among the 120 polymorphic genes, 69 had recurrent polymorphisms across isolates from multiple children (Table 3), and 72 SNPs appeared in multiple isolates from the same child (Table S4). The inferred functions of the proteins encoded by the polymorphic genes included amino acid metabolism (*n*=3 genes), antibiotic synthesis (*n*=1 gene), carbohydrate metabolism (*n*=9 genes), cell wall biosynthesis (*n*=19 genes), environmental stress response (*n*=9 genes), glycolipid metabolism (*n*=5 genes), iron metabolism (*n*=5 genes), recombination and repair (*n*=5 genes), transcriptional regulator (*n*=8 genes), transcription and translation (*n*=28 genes), small molecule transport (*n*=26 genes), and virulence (*n*=2 genes) (Figure 4A). Genes encoding proteins implicated in transcription and translation, small molecule transport, and cell wall biosynthesis were most frequently polymorphic across isolates from multiple children (Figure 4B). Additionally, these polymorphic genes were present in most of the 39 reference genomes (Table S5).

**Fig. 4.**
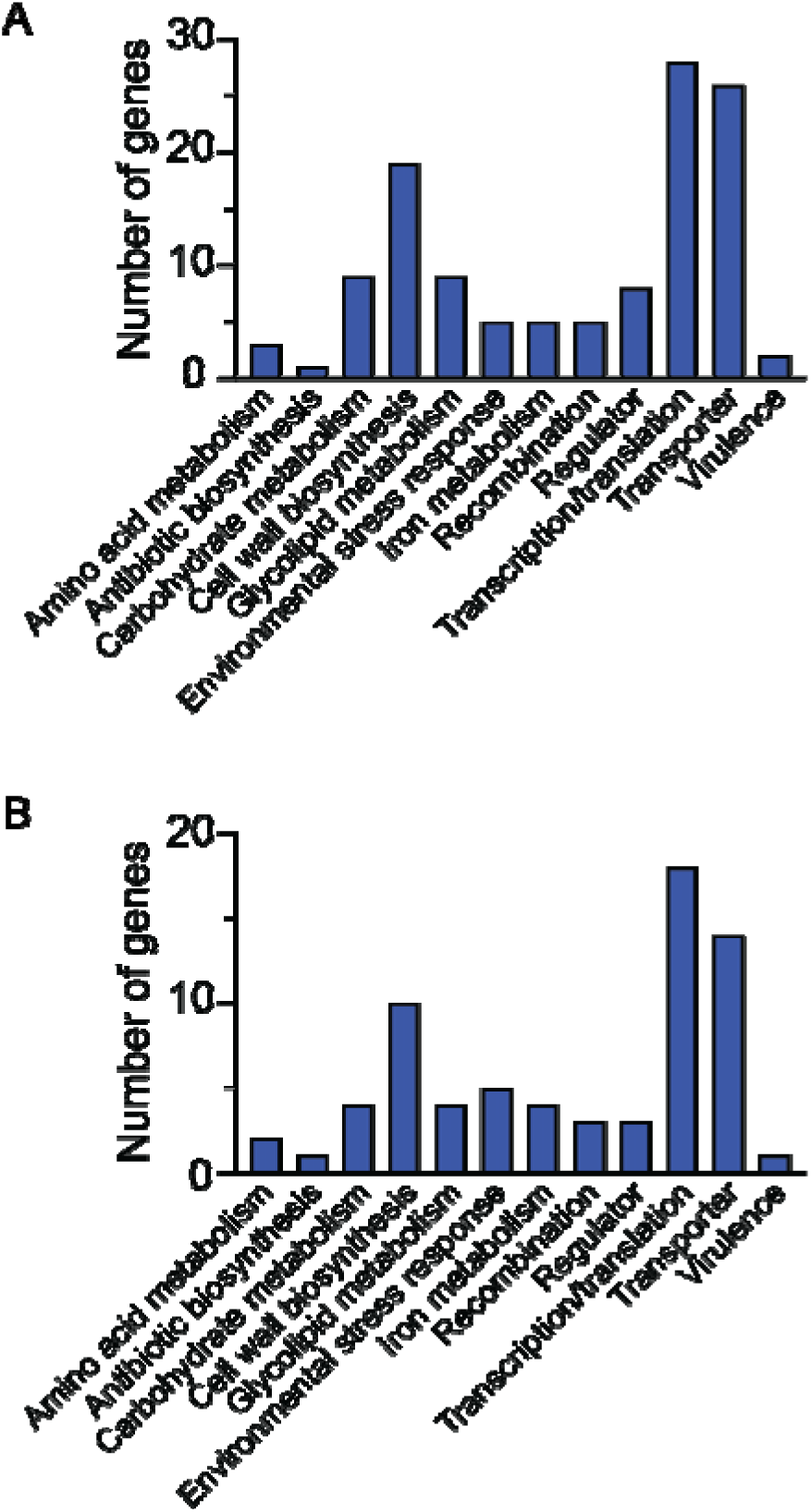
Function of genes undergoing intra-host genomic variation during human asymptomatic carriage. (**A**) Inferred function of the proteins encoded by the 120 genes that acquired a nonsynonymous or nonsense SNP in at least one asymptomatic carrier isolate. (**B**) Inferred function of the proteins encoded by the 69 genes identified as recurrently polymorphic in isolates recovered from multiple child-MLST groups.

**Table 3.**
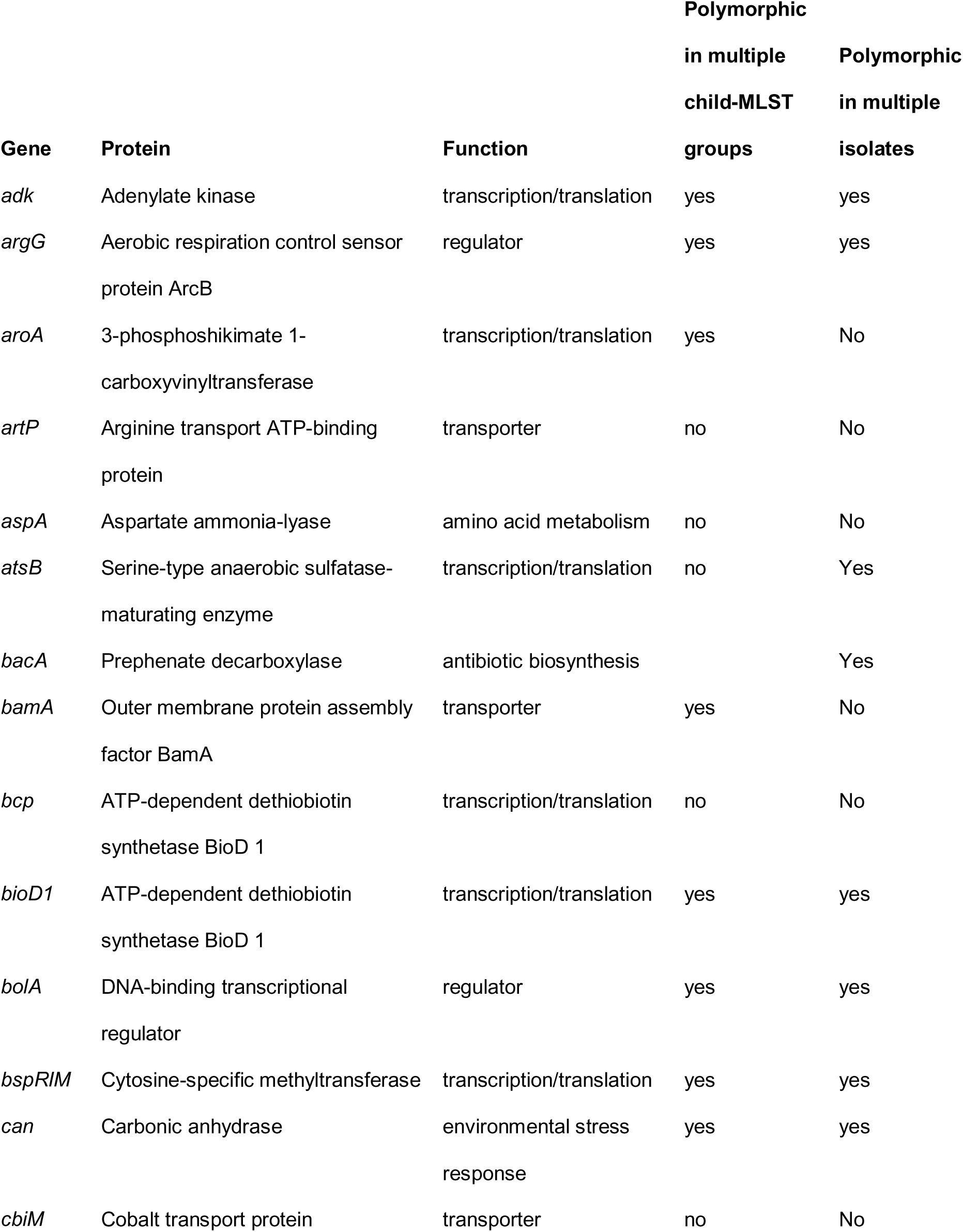
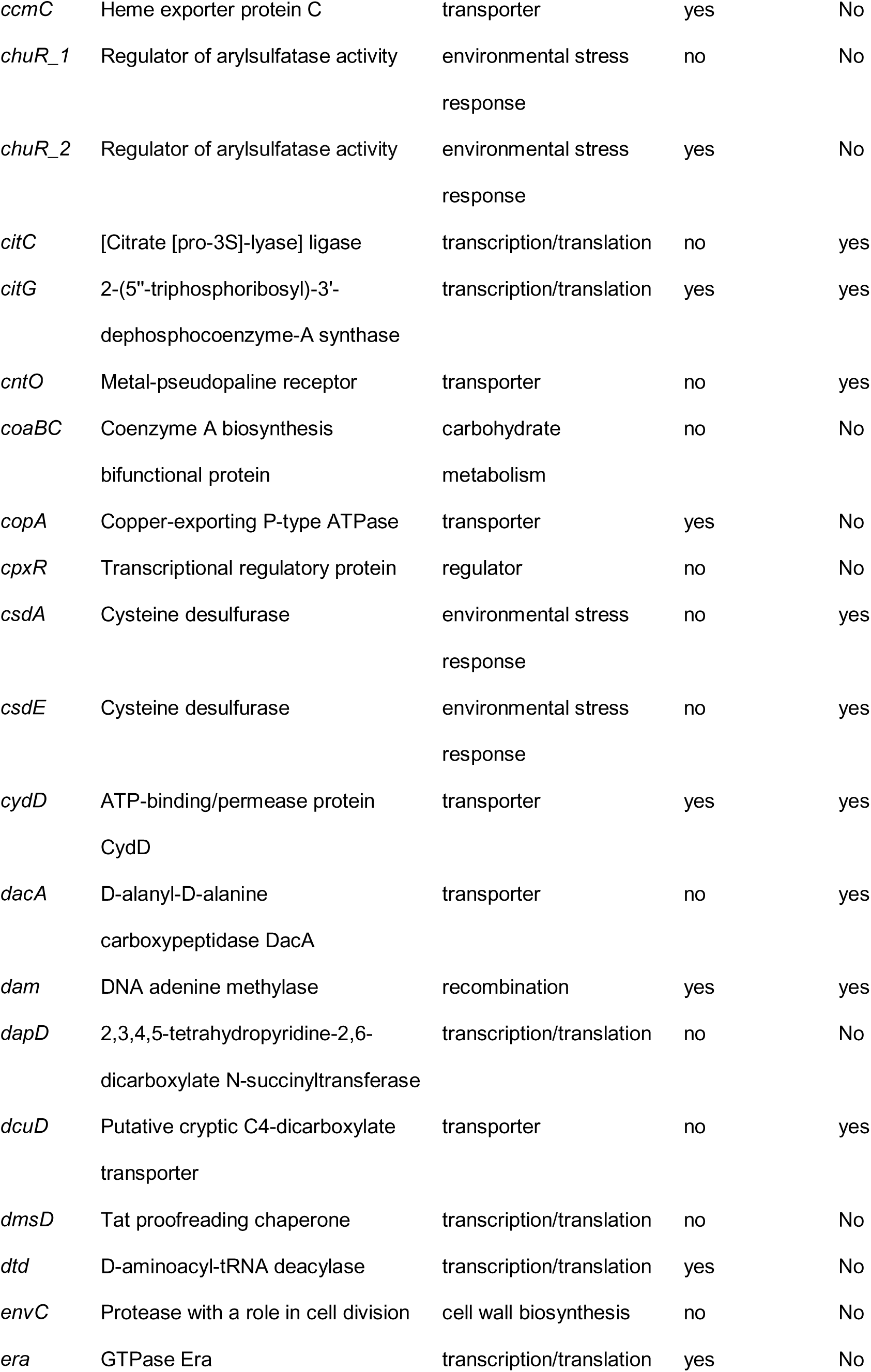
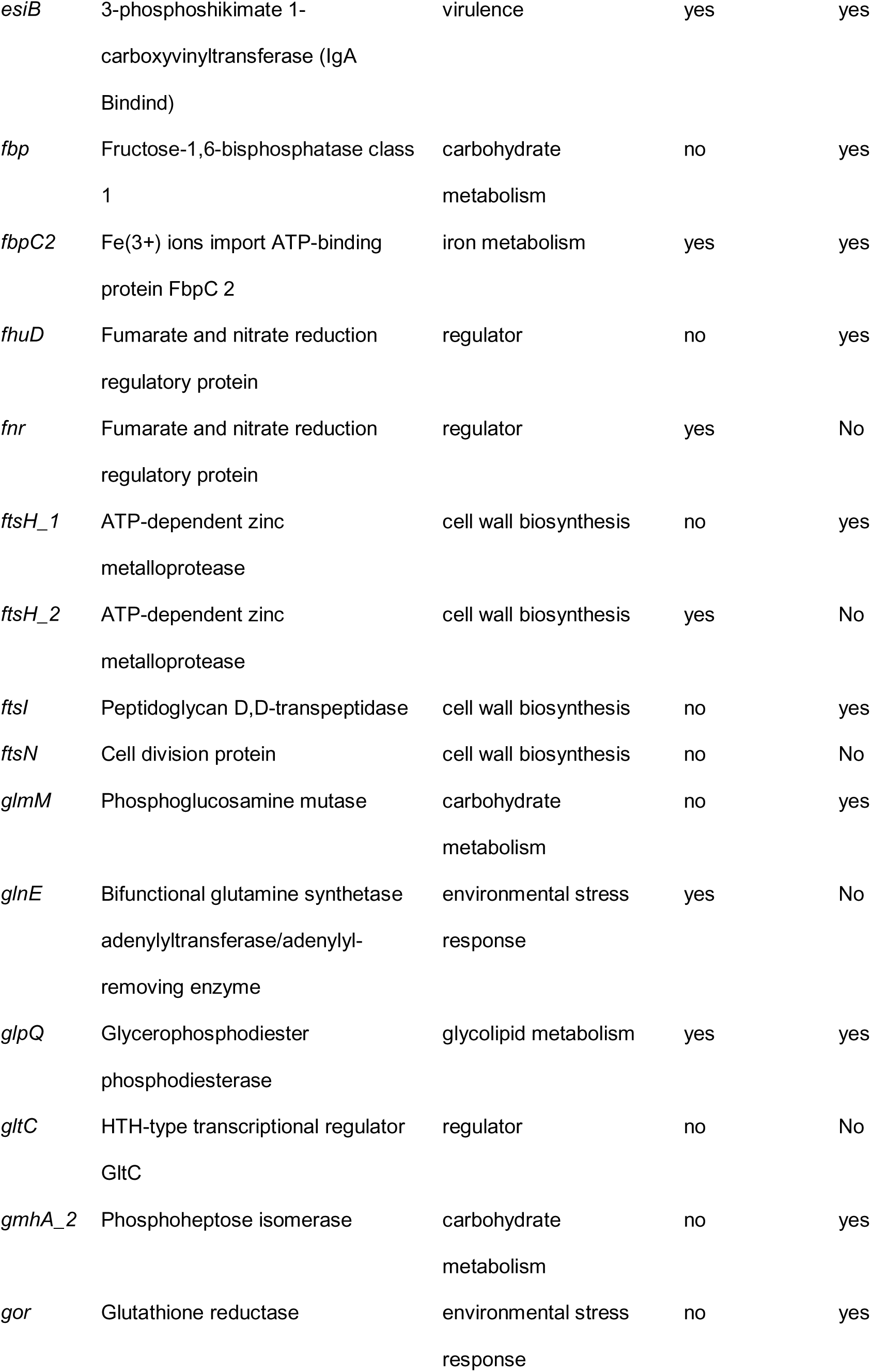
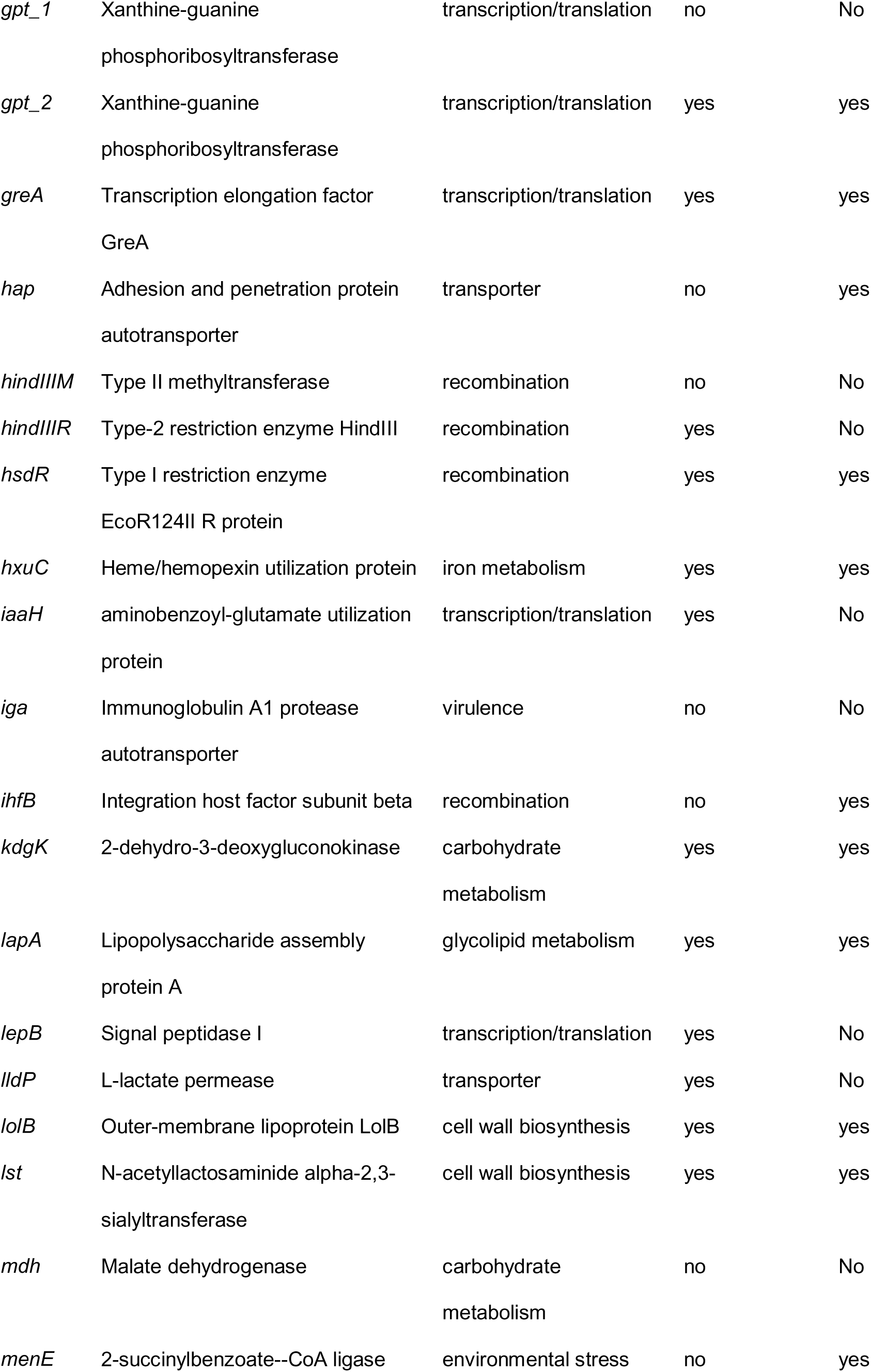
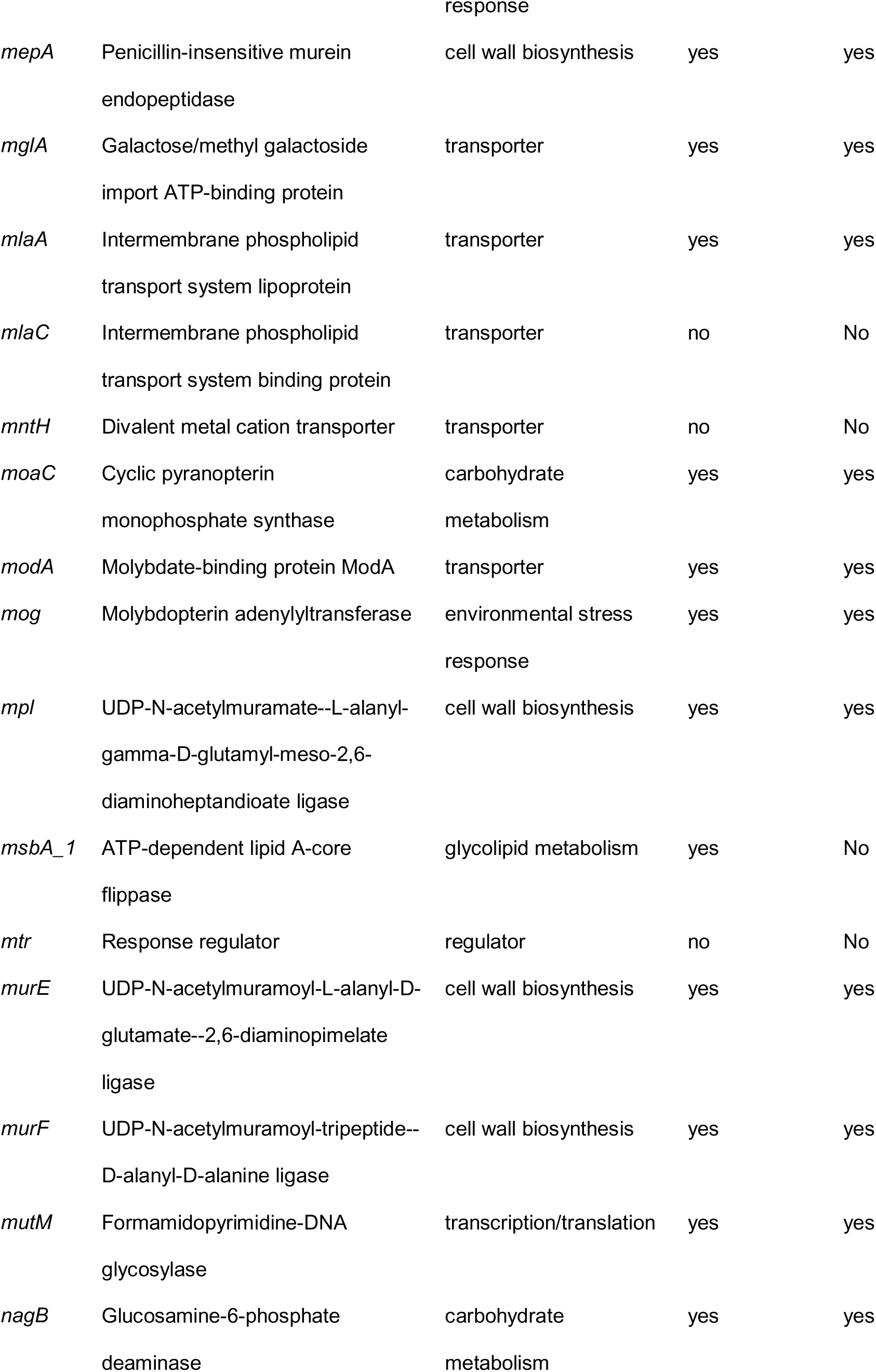
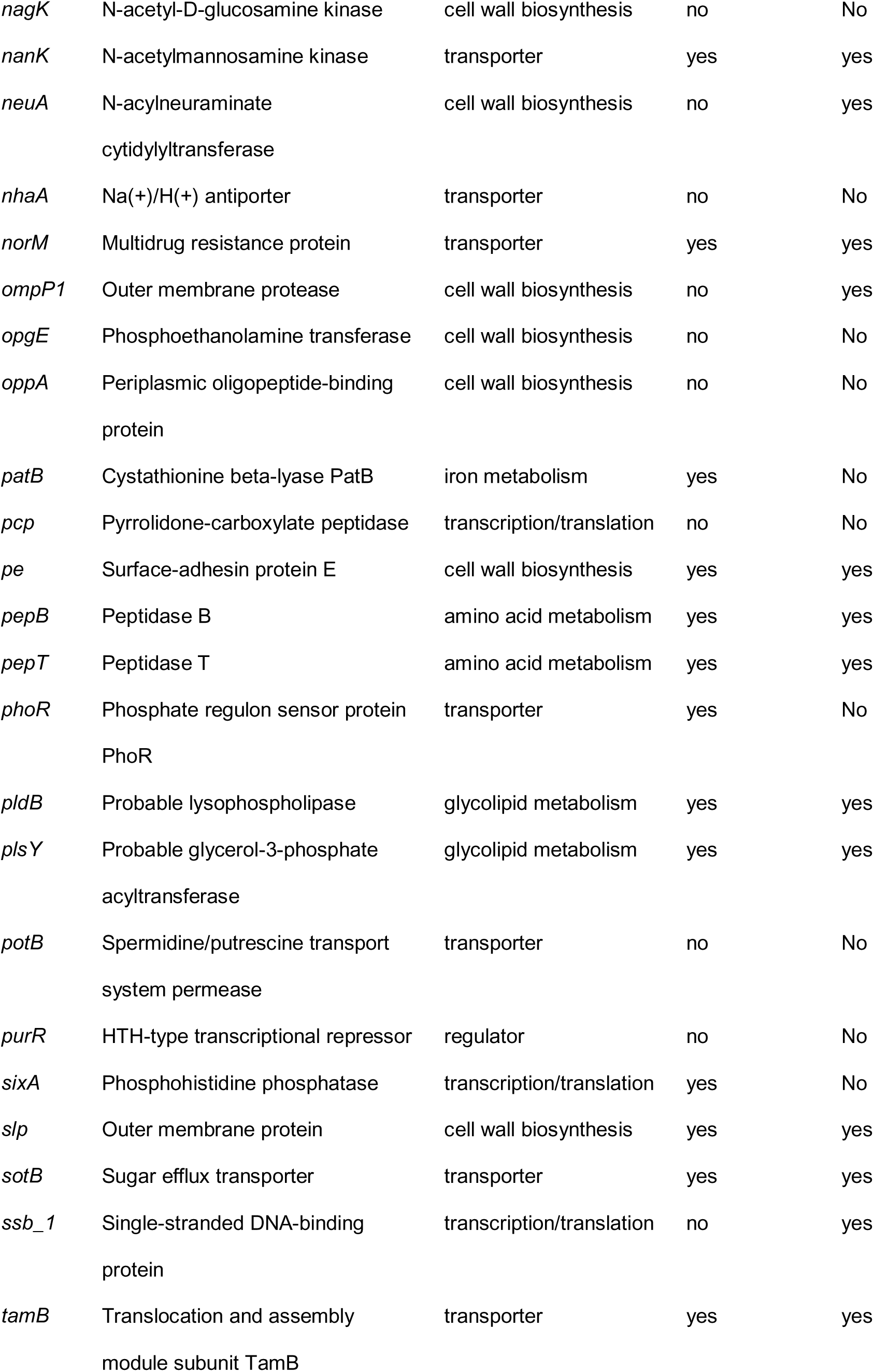
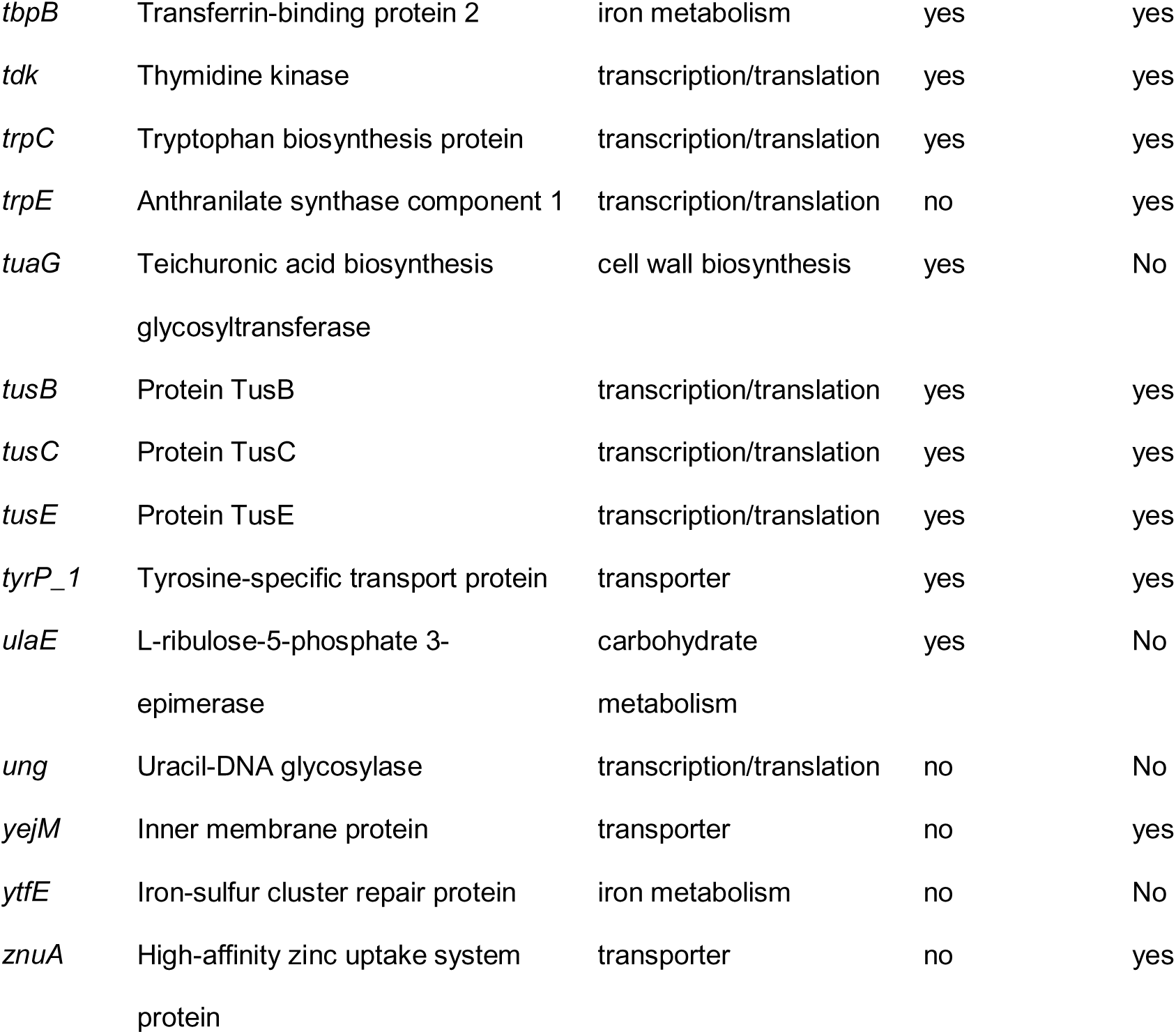
Polymorphic genes in isolates recovered from children with asymptomatic carriage.

### Diversifying selection of recurrently polymorphic genes

To test the hypothesis that amino acid substitutions and protein truncations identified in the 69 recurrently polymorphic genes occurred through diversifying selection, the nucleotide changes defining the 461 variant alleles were evaluated (Table S6). The nucleotide changes were significantly concentrated at the first two positions of variant codons, allowing rejection of the hypothesis of selective neutrality (378 observed vs. 307 expected if evenly distributed across all three positions; χ test, *P* < .0001, χ = 27.95, df = 1, z = 5.286). Further, the dN/dS ratio for the 69 recurrently polymorphic genes was 2.25, indicating positive selection for genomic diversification (Table S6). Taken together, these data support a model in which intra-host variation drives the emergence of new alleles through diversifying selection.

## DISCUSSION

Asymptomatic carriage is a common biological phenomenon that may drive microbial genomic adaption and facilitate person-to-person transmission of many pathogens ^68^. To investigate intra-host genomic variation of *H. influenzae* during asymptomatic carriage in the human nasopharynx, the genomes of 805 isolates recovered from 24 healthy Icelandic children were sequenced. Substantial intra-host genomic variation was discovered among many genes encoding proteins that likely contribute to strain fitness, virulence, and host-pathogen molecular interactions.

Two key discoveries emerged from this analysis. First, the level of genomic diversification was high compared to other studies of intra-host variation. Specifically, 120 genes, representing 6.3% of the genome, acquired nonsynonymous or nonsense SNPs in at least one isolate. In total, 335 isolates (41.6%) acquired at least one SNP. For comparison, in our recent study of intra-host genomic variation in *H. influenzae* isolates from children with otitis media, only 88 genes (4.6%) in 56 isolates (11.2%) acquired SNPs ^38^. Similarly, a previous intra-host genomic variation study of *S. pyogenes* in a nonhuman primate model of necrotizing myositis revealed SNPs in only 20% of isolates, with approximately 50% occurring in one gene ^31^. Second, frequent recurrence of polymorphic genes in the nasopharyngeal isolates was observed. Of the genes with SNPs, 69 (57.5%) were polymorphic in isolates from multiple child-MLST groups, while 72 (60.0%) were polymorphic in multiple isolates within the same child-MLST group. That is, compared to other investigations of intra-host genomic variation, diversification of *H. influenzae* in human nasopharyngeal asymptomatic carriage was common.

We recently investigated intra-host genomic variation in *H. influenzae* isolates recovered from Icelandic children with symptomatic otitis media ^38^. Since the nasopharynx and middle ear have distinct anatomic and physiologic features that likely exert unique selective pressures, we hypothesized that different genes would undergo intra-host genomic variation at each site. Consistent with this hypothesis, only three genes were recurrently polymorphic in isolates from both sites: *nanK* (N-acetylmannosamine kinase NanK), *patB* (cystathionine beta-lyase PatB), and *phoR* (phosphate regulon sensor protein PhoR). Notably, *phoR* was the most frequently polymorphic gene (4/13, 30.8% of child-MLST groups) in isolates associated with otitis media and the third most polymorphic gene (13/39, 33.3% of child-MLST groups) in isolates from asymptomatic carriage ^38^. Although the function of PhoR has not been studied in *H. influenzae*, in other bacterial species, PhoR serves as the signaling histidine kinase in a two-component regulatory system responsive to fluctuating phosphate levels ^69^. Since phosphate is naturally present in body fluids, including those in the middle ear and nasopharynx, its levels may change in various disease and health states ^70, 71^. Phosphate acquisition from the host environment is essential for many human pathogens, including *H. influenzae* ^72^. We hypothesize that phenotypic diversity in PhoR, arising through intra-host genomic variation, enhances fitness by enabling adaptation to the nutrient-limited environments of the middle ear and nasopharynx. PhoR also regulates the expression of many genes encoding proteins implicated in environmental stress response, serum bactericidal activity resistance, antimicrobial peptide resistance, and other functions that may further contribute to fitness ^72^.

As an additional example illustrating differing host-pathogen molecular interactions and selective pressures driving genomic diversification across different anatomic sites, our study of *H. influenzae* otitis media isolates revealed no acquired SNPs in genes encoding transcriptional regulators or putative virulence factors ^38^. In comparison, in the current asymptomatic carrier study, eight transcriptional regulators and two putative virulence factors exhibited polymorphisms. Notably, both of the identified virulence factors target immunoglobin A (IgA), which is present in high concentrations in saliva and nasal secretions and acts as a key host immune defense molecule in the human upper respiratory tract ^73^. In particular, *esiB*, which encodes a putative IgA-binding protein, acquired 31 distinct nonsynonymous or nonsense SNPs in isolates from eight child-MLST groups. Although *esiB* has not been previously studied in *H. influenzae*, other IgA-binding proteins have been recognized as critical virulence factors in many upper respiratory tract pathogens, such as *S. aureus* and *S. pyogenes* ^74–77^. Similarly, *iga*, encoding the immunoglobin A1 protease autotransporter that cleaves IgA1 into Fc and Fab fragments, acquired three nonsynonymous SNPs within isolates from one child ^78^. The proximity of these amino acid substitutions may represent a recombination event in this naturally competent pathogen. In general, IgA proteases are proven virulence factors ^79, 80^. In *H. influenzae*, IgA1 protease activity is significantly lower in asymptomatic nasopharyngeal carriage relative to sterile site infections ^78^. Also, amino acid substitutions are known to be structurally destabilizing and reduce IgA1 protease activity ^81^. The three amino acid substitutions occurred in the autotransporter domain, which could interfere with normal transport of the protease to the cell surface and reduce IgA1 protease activity ^63^. We hypothesize that intra-host genomic diversification of *esiB* and *iga* may favor asymptomatic carriage by altering the ability of IgA to interact with bacterial antigens or host cell receptors to mute the immune response ^82^.

This asymptomatic carrier study is the largest *H. influenzae* intra-host genomic variation investigation to date, including the largest number of subjects and isolates per subject. However, there are some limitations to the study design. First, because the children naturally acquired *H. influenzae*, the exact duration of colonization remains unknown, which could introduce an unknown bias related to gene selection or SNP accumulation. Second, only a single sampling timepoint was obtained, so day-to-day or longitudinal fluctuations in bacterial burden are unknown. Third, the samples underwent multiple passages from swab collection to genome sequencing, which could have an unknown effect on isolate recovery. Fourth, selecting samples from children with the highest nasopharyngeal colonization density could also introduce biases related to SNP diversity.

To summarize, whole-genome sequence analysis of 805 *H. influenzae* isolates from 24 healthy Icelandic children with asymptomatic nasopharyngeal carriage identified 120 polymorphic genes, including 69 genes that were recurrently polymorphic in multiple children. The genes encode proteins with diverse inferred functions that likely contribute to strain fitness, virulence, and host-pathogen molecular interactions. These data will generate new hypotheses bearing on novel targets for diagnostics, therapies, and vaccines. Future studies examining *H. influenzae* intra-host genomic variation with isolates from different anatomic sites or patient populations are warranted to expand on these findings.

## ETHICAL STATEMENT

The study was approved by the Icelandic National Bioethics Committee (license no: 12.010.S1).

## AUTHORS CONTRIBUTIONS

Conceptualization: RJO, KGK, HE, AH, JMM, and GH. Specimen collection: ST. Strain culturing, banking, cataloging, and shipping: HE, and ST. Sequencing, data collection, sequence data analysis: RJO, SWL, and YV. Analysis: all authors. Original draft preparation RJO and GH. Writing, reviewing, and editing: all authors. All authors read, provided input, and approved the final manuscript.

## ACKNOWLEDGMENTS

We thank the staff at the clinical diagnostic laboratory at Landspítali. Alma Amaya, Wendy Barragan, Merissa Dias, Ryan Gadd, Regan Mangham, Eleanor Nichols, Matthew Ojeda Saavedra, Jordan Pachuca, Sindy Pena, and Kristina Reppond at Houston Methodist Research Institute contributed to bacterial culture and genome sequencing. Zoltan Palmai, Sabeen Raza, and Stephen Beres assisted with bioinformatic analysis. Dilzi Mody provided logistical support. Sasha Pejerrey and Heather McConnell provided editorial support.

## DATA AVAILABILITY

Whole genome sequencing data for this investigation were submitted to the National Center for Biotechnology Information Sequence Read Archive and closed genomes were submitted to GenBank under BioProject accession PRJNA1248003.

## REFERENCES

[1] Khattak ZE, Anjum F: Haemophilus influenzae Infection. StatPearls. Treasure Island (FL) ineligible companies, 2025.

[2] Marrs CF, Krasan GP, McCrea KW, Clemans DL, Gilsdorf JR: Haemophilus influenzae - human specific bacteria. Front Biosci 2001, 6:E41–60.

[3] Guerra AM, Waseem M: Epiglottitis. StatPearls. Treasure Island (FL) ineligible companies, 2024.

[4] Khattak ZE, Anjum F: Haemophilus influenzae Infection. StatPearls. Treasure Island (FL) ineligible companies, 2024.

[5] Soeters HM, Blain A, Pondo T, Doman B, Farley MM, Harrison LH, Lynfield R, Miller L, Petit S, Reingold A, Schaffner W, Thomas A, Zansky SM, Wang X, Briere EC: Current Epidemiology and Trends in Invasive Haemophilus influenzae Disease-United States, 2009-2015. Clin Infect Dis 2018, 67:881–9.

[6] Slack MPE, Cripps AW, Grimwood K, Mackenzie GA, Ulanova M: Invasive Haemophilus influenzae Infections after 3 Decades of Hib Protein Conjugate Vaccine Use. Clin Microbiol Rev 2021, 34:e0002821.

[7] Oliver SE, Rubis AB, Soeters HM, Reingold A, Barnes M, Petit S, Farley MM, Harrison LH, Como-Sabetti K, Khanlian SA, Wester R, Thomas A, Schaffner W, Marjuki H, Wang X, Hariri S: Epidemiology of Invasive Nontypeable Haemophilus influenzae Disease-United States, 2008-2019. Clin Infect Dis 2023, 76:1889–95.

[8] Mukundan D, Ecevit Z, Patel M, Marrs CF, Gilsdorf JR: Pharyngeal colonization dynamics of Haemophilus influenzae and Haemophilus haemolyticus in healthy adult carriers. J Clin Microbiol 2007, 45:3207–17.

[9] St Sauver J, Marrs CF, Foxman B, Somsel P, Madera R, Gilsdorf JR: Risk factors for otitis media and carriage of multiple strains of Haemophilus influenzae and Streptococcus pneumoniae. Emerg Infect Dis 2000, 6:622–30.

[10] Ma C, Zhang Y, Wang H: Characteristics of Haemophilus influenzae carriage among healthy children in China: A meta-analysis. Medicine (Baltimore) 2023, 102:e35313.

[11] Saha S, Fozzard N, Grimwood K, Lambert SB, Ware RS: The Ages When Healthy Children Are First Colonized by Three Common Potentially Pathogenic Bacteria: A Birth Cohort Study. Pediatr Infect Dis J 2025.

[12] Giufre M, Dorrucci M, Lo Presti A, Farchi F, Cardines R, Camilli R, Pimentel de Araujo F, Mancini F, Ciervo A, Corongiu M, Pantosti A, Cerquetti M, Valdarchi C, Group F: Nasopharyngeal carriage of Haemophilus influenzae among adults with co-morbidities. Vaccine 2022, 40:826–32.

[13] Murphy TF, Brauer AL, Schiffmacher AT, Sethi S: Persistent colonization by Haemophilus influenzae in chronic obstructive pulmonary disease. Am J Respir Crit Care Med 2004, 170:266–72.

[14] Fluit AC, Bayjanov JR, Benaissa-Trouw BJ, Rogers MRC, Diez-Aguilar M, Canton R, Tunney MM, Elborn JS, Ekkelenkamp MB: Whole-genome analysis of Haemophilus influenzae strains isolated from persons with cystic fibrosis. J Med Microbiol 2022, 71.

[15] Pham LL, Varon E, Bonacorsi S, Boubaya M, Benhaim P, Amor-Chelihi L, Houlier M, Koehl B, Missud F, Brousse V, Gajdos V, Bizot E, Briand C, Malka A, Odievre MH, Romain AS, Hau I, Pondarre C, See H, Guitton C, Zenkhri F, Holvoet L, Benkerrou M, Da Silveira C, Belaid N, Laurent O, Vassal M, Basmaci R, Aupiais C, Bloch-Queyrat C, Levy C, Cohen R, Ouldali N, De Pontual L, Carbonnelle E, Gaschignard J: Nasopharyngeal Carriage and Antibiotic Resistance in Children With Sickle Cell Disease: The DREPANOBACT French Multicenter Prospective Study. Pediatr Infect Dis J 2025, 44:387–93.

[16] Barbosa-Cesnik C, Farjo RS, Patel M, Gilsdorf J, McCoy SI, Pettigrew MM, Marrs C, Foxman B: Predictors for Haemophilus influenzae colonization, antibiotic resistance and for sharing an identical isolate among children attending 16 licensed day-care centers in Michigan. Pediatr Infect Dis J 2006, 25:219–23.

[17] Farjo RS, Foxman B, Patel MJ, Zhang L, Pettigrew MM, McCoy SI, Marrs CF, Gilsdorf JR: Diversity and sharing of Haemophilus influenzae strains colonizing healthy children attending day-care centers. Pediatr Infect Dis J 2004, 23:41–6.

[18] D’Mello A, Murphy TF, Wade M, Kirkham C, Kong Y, Tettelin H, Pettigrew MM: Host-pathogen interaction profiling of nontypeable Haemophilus influenzae and Moraxella catarrhalis coinfection of bronchial epithelial cells. mSphere 2025:e0024225.

[19] Fleischmann RD, Adams MD, White O, Clayton RA, Kirkness EF, Kerlavage AR, Bult CJ, Tomb JF, Dougherty BA, Merrick JM, et al.: Whole-genome random sequencing and assembly of Haemophilus influenzae Rd. Science 1995, 269:496–512.

[20] De Chiara M, Hood D, Muzzi A, Pickard DJ, Perkins T, Pizza M, Dougan G, Rappuoli R, Moxon ER, Soriani M, Donati C: Genome sequencing of disease and carriage isolates of nontypeable Haemophilus influenzae identifies discrete population structure. Proc Natl Acad Sci U S A 2014, 111:5439–44.

[21] Pettigrew MM, Ahearn CP, Gent JF, Kong Y, Gallo MC, Munro JB, D’Mello A, Sethi S, Tettelin H, Murphy TF: Haemophilus influenzae genome evolution during persistence in the human airways in chronic obstructive pulmonary disease. Proc Natl Acad Sci U S A 2018, 115:E3256–E65.

[22] Krisna MA, Jolley KA, Monteith W, Boubour A, Hamers RL, Brueggemann AB, Harrison OB, Maiden MCJ: Development and implementation of a core genome multilocus sequence typing scheme for Haemophilus influenzae. Microb Genom 2024, 10.

[23] Tonnessen R, Garcia I, Debech N, Lindstrom JC, Wester AL, Skaare D: Molecular epidemiology and antibiotic resistance profiles of invasive Haemophilus influenzae from Norway 2017-2021. Front Microbiol 2022, 13:973257.

[24] Watts SC, Judd LM, Carzino R, Ranganathan S, Holt KE: Genomic Diversity and Antimicrobial Resistance of Haemophilus Colonizing the Airways of Young Children with Cystic Fibrosis. mSystems 2021:e0017821.

[25] Roman F, Canton R, Perez-Vazquez M, Baquero F, Campos J: Dynamics of long-term colonization of respiratory tract by Haemophilus influenzae in cystic fibrosis patients shows a marked increase in hypermutable strains. J Clin Microbiol 2004, 42:1450–9.

[26] Aziz A, Sarovich DS, Nosworthy E, Beissbarth J, Chang AB, Smith-Vaughan H, Price EP, Harris TM: Molecular Signatures of Non-typeable Haemophilus influenzae Lung Adaptation in Pediatric Chronic Lung Disease. Front Microbiol 2019, 10:1622.

[27] Murphy TF, Kirkham C, D’Mello A, Sethi S, Pettigrew MM, Tettelin H: Adaptation of Nontypeable Haemophilus influenzae in Human Airways in COPD: Genome Rearrangements and Modulation of Expression of HMW1 and HMW2. mBio 2023, 14:e0014023.

[28] Roder AE, Johnson KEE, Knoll M, Khalfan M, Wang B, Schultz-Cherry S, Banakis S, Kreitman A, Mederos C, Youn JH, Mercado R, Wang W, Chung M, Ruchnewitz D, Samanovic MI, Mulligan MJ, Lassig M, Luksza M, Das S, Gresham D, Ghedin E: Optimized quantification of intra-host viral diversity in SARS-CoV-2 and influenza virus sequence data. mBio 2023, 14:e0104623.

[29] Goldswain H, Penrice-Randal R, Donovan-Banfield I, Duffy CW, Dong X, Randle N, Ryan Y, Rzeszutek AM, Pilgrim J, Keyser E, Weller SA, Hutley EJ, Hartley C, Prince T, Darby AC, Aye Maung N, Nwume H, Hiscox JA, Emmett SR: SARS-CoV-2 population dynamics in immunocompetent individuals in a closed transmission chain shows genomic diversity over the course of infection. Genome Med 2024, 16:89.

[30] Long SW, Olsen RJ, Eagar TN, Beres SB, Zhao P, Davis JJ, Brettin T, Xia F, Musser JM: Population Genomic Analysis of 1,777 Extended-Spectrum Beta-Lactamase-Producing Klebsiella pneumoniae Isolates, Houston, Texas: Unexpected Abundance of Clonal Group 307. mBio 2017, 8.

[31] Eraso JM, Olsen RJ, Beres SB, Kachroo P, Porter AR, Nasser W, Bernard PE, DeLeo FR, Musser JM: Genomic Landscape of Intrahost Variation in Group A Streptococcus: Repeated and Abundant Mutational Inactivation of the fabT Gene Encoding a Regulator of Fatty Acid Synthesis. Infection and immunity 2016, 84:3268–81.

[32] Long SW, Beres SB, Olsen RJ, Musser JM: Absence of patient-to-patient intrahospital transmission of Staphylococcus aureus as determined by whole-genome sequencing. MBio 2014, 5:e01692–14.

[33] Houtak G, Bouras G, Nepal R, Shaghayegh G, Cooksley C, Psaltis AJ, Wormald PJ, Vreugde S: The intra-host evolutionary landscape and pathoadaptation of persistent Staphylococcus aureus in chronic rhinosinusitis. Microb Genom 2023, 9.

[34] Liu WJ, Shi W, Zhu W, Jin C, Zou S, Wang J, Ke Y, Li X, Liu M, Hu T, Fan H, Tong Y, Zhao X, Chen W, Zhao Y, Liu D, Wong G, Chen C, Geng C, Xie W, Jiang H, Kamara IL, Kamara A, Lebby M, Kargbo B, Qiu X, Wang Y, Liang X, Liang M, Dong X, Wu G, Gao GF, Shu Y: Intra-host Ebola viral adaption during human infection. Biosaf Health 2019, 1:14–24.

[35] Persyn E, Sassi M, Aubry M, Broly M, Delanou S, Asehnoune K, Caroff N, Cremet L: Rapid genetic and phenotypic changes in Pseudomonas aeruginosa clinical strains during ventilator-associated pneumonia. Sci Rep 2019, 9:4720.

[36] Cassidy H, Schuele L, Lizarazo-Forero E, Couto N, Rossen JWA, Friedrich AW, van Leer-Buter C, Niesters HGM: Exploring a prolonged enterovirus C104 infection in a severely ill patient using nanopore sequencing. Virus Evol 2022, 8:veab109.

[37] Casali N, Broda A, Harris SR, Parkhill J, Brown T, Drobniewski F: Whole Genome Sequence Analysis of a Large Isoniazid-Resistant Tuberculosis Outbreak in London: A Retrospective Observational Study. PLoS Med 2016, 13:e1002137.

[38] Olsen RJ, Long SW, Vedaraju Y, Tomasdottir S, Erlendsdottir H, Kristinsson KG, Musser JM, Haraldsson G: Intra-host genomic variation of serologically nontypeable Haemophilus influenzae isolates from otitis media. Microbiol Spectr 2025, 13:e0308924.

[39] Quirk SJ, Haraldsson G, Erlendsdottir H, Hjalmarsdottir MA, van Tonder AJ, Hrafnkelsson B, Sigurdsson S, Bentley SD, Haraldsson A, Brueggemann AB, Kristinsson KG: Effect of Vaccination on Pneumococci Isolated from the Nasopharynx of Healthy Children and the Middle Ear of Children with Otitis Media in Iceland. J Clin Microbiol 2018, 56.

[40] Sveinsdottir H, Bjornsdottir JB, Erlendsdottir H, Hjalmarsdottir MA, Hrafnkelsson B, Haraldsson A, Kristinsson KG, Haraldsson G: The Effect of the 10-Valent Pneumococcal Nontypeable Haemophilus influenzae Protein D Conjugate Vaccine on H. influenzae in Healthy Carriers and Middle Ear Infections in Iceland. J Clin Microbiol 2019, 57.

[41] Tomasdottir IA, Erlendsdottir H, Kristinsdottir I, Kristinsson KG, Haraldsson A, Beres SB, Olsen RJ, Musser JM, Thors V: A Striking Increase in Carriage Among Young Children in Iceland Paralleled the Unprecedented Increase of Invasive Group A Streptococcal Infection From 2022 to 2023. Pediatr Infect Dis J 2025.

[42] Dyvik E: Iceland - Statistics & Facts. Statistica, 2024.

[43] Leber AL, Burnham CAD: Clinical Microbiology: Procedures Handbook: John Wiley & Sons, Limited, 2025.

[44] Long SW, Olsen RJ, Nguyen HAT, Ojeda Saavedra M, Musser JM: Draft Genome Sequence of Candida auris Strain LOM, a Human Clinical Isolate from Greater Metropolitan Houston, Texas. Microbiol Resour Announc 2019, 8.

[45] Beres SB, Olsen RJ, Long SW, Langley R, Williams T, Erlendsdottir H, Smith A, Kristinsson KG, Musser JM: Increase in invasive Streptococcus pyogenes M1 infections with close evolutionary genetic relationship, Iceland and Scotland, 2022 to 2023. Euro Surveill 2024, 29.

[46] Beres SB, Olsen RJ, Long SW, Eraso JM, Boukthir S, Faili A, Kayal S, Musser JM: Analysis of the Genomics and Mouse Virulence of an Emergent Clone of Streptococcus dysgalactiae Subspecies equisimilis. Microbiol Spectr 2023, 11:e0455022.

[47] Kachroo P, Eraso JM, Beres SB, Olsen RJ, Zhu L, Nasser W, Bernard PE, Cantu CC, Saavedra MO, Arredondo MJ, Strope B, Do H, Kumaraswami M, Vuopio J, Grondahl-Yli-Hannuksela K, Kristinsson KG, Gottfredsson M, Pesonen M, Pensar J, Davenport ER, Clark AG, Corander J, Caugant DA, Gaini S, Magnussen MD, Kubiak SL, Nguyen HAT, Long SW, Porter AR, DeLeo FR, Musser JM: Integrated analysis of population genomics, transcriptomics and virulence provides novel insights into Streptococcus pyogenes pathogenesis. Nat Genet 2019, 51:548–59.

[48] Bolger AM, Lohse M, Usadel B: Trimmomatic: a flexible trimmer for Illumina sequence data. Bioinformatics 2014, 30:2114–20.

[49] Edwards JA, Edwards RA: Fastq-pair: efficient synchronization of paired-end fastq files. bioRxiv 2019:552885.

[50] Liu Y, Schroder J, Schmidt B: Musket: a multistage k-mer spectrum-based error corrector for Illumina sequence data. Bioinformatics 2013, 29:308–15.

[51] Inouye M, Dashnow H, Raven LA, Schultz MB, Pope BJ, Tomita T, Zobel J, Holt KE: SRST2: Rapid genomic surveillance for public health and hospital microbiology labs. Genome Med 2014, 6:90.

[52] Holley G, Melsted P: Bifrost: highly parallel construction and indexing of colored and compacted de Bruijn graphs. Genome Biol 2020, 21:249.

[53] Schulz T, Wittler R, Stoye J: Sequence-based pangenomic core detection. iScience 2022, 25:104413.

[54] Kurtz S, Phillippy A, Delcher AL, Smoot M, Shumway M, Antonescu C, Salzberg SL: Versatile and open software for comparing large genomes. Genome Biol 2004, 5:R12.

[55] Huson DH: SplitsTree: analyzing and visualizing evolutionary data. Bioinformatics 1998, 14:68–73.

[56] Wick RR, Judd LM, Holt KE: Assembling the perfect bacterial genome using Oxford Nanopore and Illumina sequencing. PLoS Comput Biol 2023, 19:e1010905.

[57] Bankevich A, Nurk S, Antipov D, Gurevich AA, Dvorkin M, Kulikov AS, Lesin VM, Nikolenko SI, Pham S, Prjibelski AD, Pyshkin AV, Sirotkin AV, Vyahhi N, Tesler G, Alekseyev MA, Pevzner PA: SPAdes: a new genome assembly algorithm and its applications to single-cell sequencing. J Comput Biol 2012, 19:455–77.

[58] Vaser R, Sovic I, Nagarajan N, Sikic M: Fast and accurate de novo genome assembly from long uncorrected reads. Genome Res 2017, 27:737–46.

[59] Wick RR, Judd LM, Gorrie CL, Holt KE: Unicycler: Resolving bacterial genome assemblies from short and long sequencing reads. PLoS Comput Biol 2017, 13:e1005595.

[60] Seemann T: Prokka: rapid prokaryotic genome annotation. Bioinformatics 2014, 30:2068–9.

[61] Carver T, Harris SR, Berriman M, Parkhill J, McQuillan JA: Artemis: an integrated platform for visualization and analysis of high-throughput sequence-based experimental data. Bioinformatics 2012, 28:464–9.

[62] Tamura K, Stecher G, Kumar S: MEGA11: Molecular Evolutionary Genetics Analysis Version 11. Mol Biol Evol 2021, 38:3022–7.

[63] UniProt C: UniProt: the Universal Protein Knowledgebase in 2023. Nucleic Acids Res 2023, 51:D523–D31.

[64] Jolley KA, Bray JE, Maiden MCJ: Open-access bacterial population genomics: BIGSdb software, the PubMLST.org website and their applications. Wellcome Open Res 2018, 3:124.

[65] de Gier C, Kirkham LA, Norskov-Lauritsen N: Complete Deletion of the Fucose Operon in Haemophilus influenzae Is Associated with a Cluster in Multilocus Sequence Analysis-Based Phylogenetic Group II Related to Haemophilus haemolyticus: Implications for Identification and Typing. J Clin Microbiol 2015, 53:3773–8.

[66] Shuel ML, Karlowsky KE, Law DK, Tsang RS: Nonencapsulated or nontypeable Haemophilus influenzae are more likely than their encapsulated or serotypeable counterparts to have mutations in their fucose operon. Can J Microbiol 2011, 57:982–6.

[67] Bush SJ, Foster D, Eyre DW, Clark EL, De Maio N, Shaw LP, Stoesser N, Peto TEA, Crook DW, Walker AS: Genomic diversity affects the accuracy of bacterial single-nucleotide polymorphism-calling pipelines. Gigascience 2020, 9.

[68] Chisholm RH, Campbell PT, Wu Y, Tong SYC, McVernon J, Geard N: Implications of asymptomatic carriers for infectious disease transmission and control. R Soc Open Sci 2018, 5:172341.

[69] Wanner BL: Is cross regulation by phosphorylation of two-component response regulator proteins important in bacteria? J Bacteriol 1992, 174:2053–8.

[70] Makiishi-Shimobayashi C, Tsujimura T, Iwasaki T, Kakihana M, Shimano K, Terada N, Sakagami M: Localization of osteopontin at calcification sites of cholesteatoma: possible role as a regulator of deposition of calcium phosphate in the middle ear. Auris Nasus Larynx 2004, 31:3–9.

[71] Noben-Trauth K, Latoche JR: Ectopic mineralization in the middle ear and chronic otitis media with effusion caused by RPL38 deficiency in the Tail-short (Ts) mouse. J Biol Chem 2011, 286:3079–93.

[72] Lamarche MG, Wanner BL, Crepin S, Harel J: The phosphate regulon and bacterial virulence: a regulatory network connecting phosphate homeostasis and pathogenesis. FEMS Microbiol Rev 2008, 32:461–73.

[73] Macpherson AJ, McCoy KD, Johansen FE, Brandtzaeg P: The immune geography of IgA induction and function. Mucosal Immunol 2008, 1:11–22.

[74] Bolton A, Song XM, Willson P, Fontaine MC, Potter AA, Perez-Casal J: Use of the surface proteins GapC and Mig of Streptococcus dysgalactiae as potential protective antigens against bovine mastitis. Can J Microbiol 2004, 50:423–32.

[75] Kong F, Gidding HF, Berner R, Gilbert GL: Streptococcus agalactiae Cbeta protein gene (bac) sequence types, based on the repeated region of the cell-wall-spanning domain: relationship to virulence and a proposed standardized nomenclature. J Med Microbiol 2006, 55:829–37.

[76] Wines BD, Ramsland PA, Trist HM, Gardam S, Brink R, Fraser JD, Hogarth PM: Interaction of human, rat, and mouse immunoglobulin A (IgA) with Staphylococcal superantigen-like 7 (SSL7) decoy protein and leukocyte IgA receptor. J Biol Chem 2011, 286:33118–24.

[77] Woof JM: The human IgA-Fc alpha receptor interaction and its blockade by streptococcal IgA-binding proteins. Biochem Soc Trans 2002, 30:491–4.

[78] Vitovski S, Dunkin KT, Howard AJ, Sayers JR: Nontypeable Haemophilus influenzae in carriage and disease: a difference in IgA1 protease activity levels. JAMA 2002, 287:1699–705.

[79] Murphy TF, Kirkham C, Jones MM, Sethi S, Kong Y, Pettigrew MM: Expression of IgA Proteases by Haemophilus influenzae in the Respiratory Tract of Adults With Chronic Obstructive Pulmonary Disease. J Infect Dis 2015, 212:1798–805.

[80] Murphy TF, Kirkham C, Gallo MC, Yang Y, Wilding GE, Pettigrew MM: Immunoglobulin A Protease Variants Facilitate Intracellular Survival in Epithelial Cells By Nontypeable Haemophilus influenzae That Persist in the Human Respiratory Tract in Chronic Obstructive Pulmonary Disease. J Infect Dis 2017, 216:1295–302.

[81] Chen CW, Ho CH: Substitutions in the nonactive site of the passenger domain on the activity of Haemophilus influenzae immunoglobulin A1 protease. Infection and immunity 2024, 92:e0019324.

[82] Woof JM, Russell MW: Structure and function relationships in IgA. Mucosal Immunology 2011, 4:590–7.

