## Supplemental Files for "Intra-host genomic variation of *Haemophilus influenzae* isolates from asymptomatic nasopharyngeal carriers involves genes encoding proteins with diverse inferred functions"

Table S1. Gene content of 805 *H. influenzae* isolates.

| Strain | Child | MLST | *fuc Operon* | Capsule Genotype | Plasmid Replicons | AMR Genes |
|---|---|---|---|---|---|---|
| MHI3301 | 1 | 266 | present | unencapsulated | none detected | none detected |
| MHI3302 | 1 | 266 | present | unencapsulated | none detected | none detected |
| MHI3303 | 1 | 266 | present | unencapsulated | none detected | none detected |
| MHI3304 | 1 | 266 | present | unencapsulated | none detected | none detected |
| MHI3305 | 1 | 266 | present | unencapsulated | none detected | none detected |
| MHI3306 | 1 | 266 | present | unencapsulated | none detected | none detected |
| MHI3307 | 1 | 266 | present | unencapsulated | none detected | none detected |
| MHI3308 | 1 | 266 | present | unencapsulated | none detected | none detected |
| MHI3309 | 1 | 266 | present | unencapsulated | none detected | none detected |
| MHI3310 | 1 | 266 | present | unencapsulated | none detected | none detected |
| MHI3311 | 1 | 266 | present | unencapsulated | none detected | none detected |
| MHI3312 | 1 | 266 | present | unencapsulated | none detected | none detected |
| MHI3313 | 1 | 266 | present | unencapsulated | none detected | none detected |
| MHI3315 | 1 | 266 | present | unencapsulated | none detected | none detected |
| MHI3316 | 1 | 266 | present | unencapsulated | none detected | none detected |
| MHI3317 | 1 | 266 | present | unencapsulated | none detected | none detected |
| MHI3318 | 1 | 266 | present | unencapsulated | none detected | none detected |
| MHI3319 | 1 | 266 | present | unencapsulated | none detected | none detected |
| MHI3320 | 1 | 266 | present | unencapsulated | none detected | none detected |
| MHI3321 | 1 | 266 | present | unencapsulated | none detected | none detected |
| MHI3322 | 1 | 266 | present | unencapsulated | none detected | none detected |
| MHI3323 | 1 | 266 | present | unencapsulated | none detected | none detected |
| MHI3324 | 1 | 266 | present | unencapsulated | none detected | none detected |
| MHI3325 | 1 | 266 | present | unencapsulated | none detected | none detected |
| MHI3326 | 1 | 266 | present | unencapsulated | none detected | none detected |
| MHI3327 | 1 | 266 | present | unencapsulated | none detected | none detected |
| MHI3328 | 1 | 266 | present | unencapsulated | none detected | none detected |
| MHI3329 | 1 | 266 | present | unencapsulated | none detected | none detected |
| MHI3330 | 1 | 266 | present | unencapsulated | none detected | none detected |
| MHI3574 | 1 | 266 | present | unencapsulated | none detected | none detected |
| MHI3575 | 1 | 266 | present | unencapsulated | none detected | none detected |
| MHI4367 | 1 | 266 | present | unencapsulated | none detected | none detected |
| MHI4368 | 1 | 266 | present | unencapsulated | none detected | none detected |
| MHI4369 | 1 | 266 | present | unencapsulated | none detected | none detected |
| MHI4370 | 1 | 266 | present | unencapsulated | none detected | none detected |
| MHI4371 | 1 | 266 | present | unencapsulated | none detected | none detected |
| MHI4372 | 1 | 266 | present | unencapsulated | none detected | none detected |
| MHI4373 | 1 | 266 | present | unencapsulated | none detected | none detected |
| MHI4374 | 1 | 266 | present | unencapsulated | none detected | none detected |
| MHI3332 | 2 | 544 | present | unencapsulated | none detected | none detected |
| MHI3334 | 2 | 544 | present | unencapsulated | none detected | none detected |
| MHI3336 | 2 | 544 | present | unencapsulated | none detected | none detected |
| MHI3337 | 2 | 544 | present | unencapsulated | none detected | none detected |
| MHI3348 | 2 | 544 | present | unencapsulated | none detected | none detected |
| MHI3351 | 2 | 544 | present | unencapsulated | none detected | none detected |
| MHI3354 | 2 | 544 | present | unencapsulated | none detected | none detected |
| MHI3341 | 2 | 652 | present | unencapsulated | none detected | TEM |
| MHI3331 | 2 | 1040 | present | unencapsulated | none detected | TEM |
| MHI3333 | 2 | 1040 | present | unencapsulated | none detected | TEM |
| MHI3335 | 2 | 1040 | present | unencapsulated | none detected | TEM |
| MHI3343 | 2 | 1040 | present | unencapsulated | none detected | TEM |
| MHI3344 | 2 | 1040 | present | unencapsulated | none detected | TEM |
| MHI3346 | 2 | 1040 | present | unencapsulated | none detected | TEM |
| MHI3349 | 2 | 1040 | present | unencapsulated | none detected | TEM |

| | | | | | | |
|---|---|---|---|---|---|---|
| MHI3352 | 2 | 1040 | present | unencapsulated | none detected | TEM |
| MHI3353 | 2 | 1040 | present | unencapsulated | none detected | TEM |
| MHI3357 | 2 | 1040 | present | unencapsulated | none detected | TEM |
| MHI3360 | 2 | 1040 | present | unencapsulated | none detected | TEM |
| MHI3576 | 2 | 1040 | present | unencapsulated | none detected | TEM |
| MHI4375 | 2 | 1040 | present | unencapsulated | none detected | none detected |
| MHI3338 | 2 | 1191 | present | unencapsulated | none detected | none detected |
| MHI3358 | 2 | 1191 | present | unencapsulated | none detected | none detected |
| MHI3577 | 2 | 1191 | present | unencapsulated | none detected | none detected |
| MHI4378 | 2 | 1191 | present | unencapsulated | none detected | none detected |
| MHI3578 | 7 | 159 | present | unencapsulated | none detected | none detected |
| MHI3579 | 7 | 159 | present | unencapsulated | none detected | none detected |
| MHI3580 | 7 | 159 | present | unencapsulated | none detected | none detected |
| MHI3581 | 7 | 159 | present | unencapsulated | none detected | none detected |
| MHI3582 | 7 | 159 | present | unencapsulated | none detected | none detected |
| MHI3583 | 7 | 159 | present | unencapsulated | none detected | none detected |
| MHI3584 | 7 | 159 | present | unencapsulated | none detected | none detected |
| MHI3585 | 7 | 159 | present | unencapsulated | none detected | none detected |
| MHI3586 | 7 | 159 | present | unencapsulated | none detected | none detected |
| MHI3587 | 7 | 159 | present | unencapsulated | none detected | none detected |
| MHI3588 | 7 | 159 | present | unencapsulated | none detected | none detected |
| MHI3589 | 7 | 159 | present | unencapsulated | none detected | none detected |
| MHI3601 | 7 | 159 | present | unencapsulated | none detected | none detected |
| MHI3602 | 7 | 159 | present | unencapsulated | none detected | none detected |
| MHI3603 | 7 | 159 | present | unencapsulated | none detected | none detected |
| MHI3605 | 7 | 159 | present | unencapsulated | none detected | none detected |
| MHI3606 | 7 | 159 | present | unencapsulated | none detected | none detected |
| MHI3607 | 7 | 159 | present | unencapsulated | none detected | none detected |
| MHI3608 | 7 | 159 | present | unencapsulated | none detected | none detected |
| MHI3609 | 7 | 159 | present | unencapsulated | none detected | none detected |
| MHI4381 | 7 | 159 | present | unencapsulated | none detected | none detected |
| MHI4382 | 7 | 159 | present | unencapsulated | none detected | none detected |
| MHI4383 | 7 | 159 | present | unencapsulated | none detected | none detected |
| MHI4384 | 7 | 159 | present | unencapsulated | none detected | none detected |
| MHI4385 | 7 | 159 | present | unencapsulated | none detected | none detected |
| MHI4386 | 7 | 159 | present | unencapsulated | none detected | none detected |
| MHI3941 | 8 | 431 | present | unencapsulated | none detected | none detected |
| MHI3942 | 8 | 431 | present | unencapsulated | none detected | none detected |
| MHI3943 | 8 | 431 | present | unencapsulated | none detected | none detected |
| MHI3944 | 8 | 431 | present | unencapsulated | none detected | none detected |
| MHI3945 | 8 | 431 | present | unencapsulated | none detected | none detected |
| MHI3946 | 8 | 431 | present | unencapsulated | none detected | none detected |
| MHI3947 | 8 | 431 | present | unencapsulated | none detected | none detected |
| MHI3948 | 8 | 431 | present | unencapsulated | none detected | none detected |
| MHI3949 | 8 | 431 | present | unencapsulated | none detected | none detected |
| MHI3950 | 8 | 431 | present | unencapsulated | none detected | none detected |
| MHI3951 | 8 | 431 | present | unencapsulated | none detected | none detected |
| MHI3952 | 8 | 431 | present | unencapsulated | none detected | none detected |
| MHI3953 | 8 | 431 | present | unencapsulated | none detected | none detected |
| MHI3954 | 8 | 431 | present | unencapsulated | none detected | none detected |
| MHI3955 | 8 | 431 | present | unencapsulated | none detected | none detected |
| MHI3956 | 8 | 431 | present | unencapsulated | none detected | none detected |
| MHI3957 | 8 | 431 | present | unencapsulated | none detected | none detected |
| MHI3958 | 8 | 431 | present | unencapsulated | none detected | none detected |
| MHI3959 | 8 | 431 | present | unencapsulated | none detected | none detected |
| MHI3960 | 8 | 431 | present | unencapsulated | none detected | none detected |
| MHI3961 | 8 | 431 | present | unencapsulated | none detected | none detected |

| | | | | | | |
|---|---|---|---|---|---|---|
| MHI3962 | 8 | 431 | present | unencapsulated | none detected | none detected |
| MHI3967 | 8 | 431 | present | unencapsulated | none detected | none detected |
| MHI3969 | 8 | 431 | present | unencapsulated | none detected | none detected |
| MHI3970 | 8 | 431 | present | unencapsulated | none detected | none detected |
| MHI3971 | 8 | 431 | present | unencapsulated | none detected | none detected |
| MHI3972 | 8 | 431 | present | unencapsulated | none detected | none detected |
| MHI3973 | 8 | 431 | present | unencapsulated | none detected | none detected |
| MHI3974 | 8 | 431 | present | unencapsulated | none detected | none detected |
| MHI3963 | 8 | 2955 | present | unencapsulated | none detected | none detected |
| MHI3964 | 8 | 2955 | present | unencapsulated | none detected | none detected |
| MHI3965 | 8 | 2955 | present | unencapsulated | none detected | none detected |
| MHI3966 | 8 | 2955 | present | unencapsulated | none detected | none detected |
| MHI3975 | 8 | 2955 | present | unencapsulated | none detected | none detected |
| MHI3976 | 10 | 57 | present | unencapsulated | none detected | none detected |
| MHI3977 | 10 | 57 | present | unencapsulated | none detected | none detected |
| MHI3978 | 10 | 57 | present | unencapsulated | none detected | none detected |
| MHI3979 | 10 | 57 | present | unencapsulated | none detected | none detected |
| MHI3980 | 10 | 57 | present | unencapsulated | none detected | none detected |
| MHI3981 | 10 | 57 | present | unencapsulated | none detected | none detected |
| MHI3982 | 10 | 57 | present | unencapsulated | none detected | none detected |
| MHI3983 | 10 | 57 | present | unencapsulated | none detected | none detected |
| MHI3984 | 10 | 57 | present | unencapsulated | none detected | none detected |
| MHI3985 | 10 | 57 | present | unencapsulated | none detected | none detected |
| MHI3986 | 10 | 57 | present | unencapsulated | none detected | none detected |
| MHI3987 | 10 | 57 | present | unencapsulated | none detected | none detected |
| MHI3988 | 10 | 57 | present | unencapsulated | none detected | none detected |
| MHI3989 | 10 | 57 | present | unencapsulated | none detected | none detected |
| MHI3990 | 10 | 57 | present | unencapsulated | none detected | none detected |
| MHI3991 | 10 | 57 | present | unencapsulated | none detected | none detected |
| MHI3992 | 10 | 57 | present | unencapsulated | none detected | none detected |
| MHI3993 | 10 | 57 | present | unencapsulated | none detected | none detected |
| MHI3994 | 10 | 57 | present | unencapsulated | none detected | none detected |
| MHI3995 | 10 | 57 | present | unencapsulated | none detected | none detected |
| MHI3996 | 10 | 57 | present | unencapsulated | none detected | none detected |
| MHI3997 | 10 | 57 | present | unencapsulated | none detected | none detected |
| MHI3998 | 10 | 57 | present | unencapsulated | none detected | none detected |
| MHI3999 | 10 | 57 | present | unencapsulated | none detected | none detected |
| MHI4000 | 10 | 57 | present | unencapsulated | none detected | none detected |
| MHI4001 | 10 | 57 | present | unencapsulated | none detected | none detected |
| MHI4002 | 10 | 57 | present | unencapsulated | none detected | none detected |
| MHI4003 | 10 | 57 | present | unencapsulated | none detected | none detected |
| MHI4004 | 10 | 57 | present | unencapsulated | none detected | none detected |
| MHI4005 | 10 | 57 | present | unencapsulated | none detected | none detected |
| MHI4006 | 10 | 57 | present | unencapsulated | none detected | none detected |
| MHI4007 | 10 | 57 | present | unencapsulated | none detected | none detected |
| MHI4008 | 10 | 57 | present | unencapsulated | none detected | none detected |
| MHI4009 | 10 | 57 | present | unencapsulated | none detected | none detected |
| MHI4010 | 10 | 57 | present | unencapsulated | none detected | none detected |
| MHI4011 | 10 | 57 | present | unencapsulated | none detected | none detected |
| MHI4012 | 10 | 57 | present | unencapsulated | none detected | none detected |
| MHI4013 | 10 | 57 | present | unencapsulated | none detected | none detected |
| MHI4014 | 10 | 57 | present | unencapsulated | none detected | none detected |
| MHI4015 | 13 | 2156 | present | unencapsulated | none detected | none detected |
| MHI4016 | 13 | 2156 | present | unencapsulated | none detected | none detected |

| | | | | | | |
|---|---|---|---|---|---|---|
| MHI4017 | 13 | 2156 | present | unencapsulated | none detected | none detected |
| MHI4018 | 13 | 2156 | present | unencapsulated | none detected | none detected |
| MHI4019 | 13 | 2156 | present | unencapsulated | none detected | none detected |
| MHI4020 | 13 | 2156 | present | unencapsulated | none detected | none detected |
| MHI4022 | 13 | 2156 | present | unencapsulated | none detected | none detected |
| MHI4023 | 13 | 2156 | present | unencapsulated | none detected | none detected |
| MHI4024 | 13 | 2156 | present | unencapsulated | none detected | none detected |
| MHI4025 | 13 | 2156 | present | unencapsulated | none detected | none detected |
| MHI4026 | 13 | 2156 | present | unencapsulated | none detected | none detected |
| MHI4027 | 13 | 2156 | present | unencapsulated | none detected | none detected |
| MHI4028 | 13 | 2156 | present | unencapsulated | none detected | none detected |
| MHI4029 | 13 | 2156 | present | unencapsulated | none detected | none detected |
| MHI4030 | 13 | 2156 | present | unencapsulated | none detected | none detected |
| MHI4031 | 13 | 2156 | present | unencapsulated | none detected | none detected |
| MHI4032 | 13 | 2156 | present | unencapsulated | none detected | none detected |
| MHI4033 | 13 | 2156 | present | unencapsulated | none detected | none detected |
| MHI4034 | 13 | 2156 | present | unencapsulated | none detected | none detected |
| MHI4035 | 13 | 2156 | present | unencapsulated | none detected | none detected |
| MHI4036 | 13 | 2156 | present | unencapsulated | none detected | none detected |
| MHI4037 | 13 | 2156 | present | unencapsulated | none detected | none detected |
| MHI4038 | 13 | 2156 | present | unencapsulated | none detected | none detected |
| MHI4039 | 13 | 2156 | present | unencapsulated | none detected | none detected |
| MHI4040 | 13 | 2156 | present | unencapsulated | none detected | none detected |
| MHI4041 | 13 | 2156 | present | unencapsulated | none detected | none detected |
| MHI4042 | 13 | 2156 | present | unencapsulated | none detected | none detected |
| MHI4043 | 13 | 2156 | present | unencapsulated | none detected | none detected |
| MHI4044 | 13 | 2156 | present | unencapsulated | none detected | none detected |
| MHI4045 | 13 | 2156 | present | unencapsulated | none detected | none detected |
| MHI4046 | 13 | 2156 | present | unencapsulated | none detected | none detected |
| MHI4047 | 13 | 2156 | present | unencapsulated | none detected | none detected |
| MHI4048 | 13 | 2156 | present | unencapsulated | none detected | none detected |
| MHI4049 | 13 | 2156 | present | unencapsulated | none detected | none detected |
| MHI4060 | 15 | 1238 | present | unencapsulated | none detected | none detected |
| MHI4068 | 15 | 1238 | present | unencapsulated | none detected | none detected |
| MHI4074 | 15 | 1238 | present | unencapsulated | none detected | none detected |
| MHI4050 | 15 | 2956 | *fucK* deleted | unencapsulated | none detected | none detected |
| MHI4052 | 15 | 2956 | *fucK* deleted | unencapsulated | none detected | none detected |
| MHI4056 | 15 | 2956 | *fucK* deleted | unencapsulated | none detected | none detected |
| MHI4058 | 15 | 2956 | *fucK* deleted | unencapsulated | none detected | none detected |
| MHI4059 | 15 | 2956 | *fucK* deleted | unencapsulated | none detected | none detected |
| MHI4079 | 15 | 2956 | *fucK* deleted | unencapsulated | none detected | none detected |
| MHI4085 | 15 | 2956 | *fucK* deleted | unencapsulated | none detected | none detected |
| MHI4051 | 15 | 2957 | present | unencapsulated | none detected | none detected |
| MHI4053 | 15 | 2957 | present | unencapsulated | none detected | none detected |
| MHI4054 | 15 | 2957 | present | unencapsulated | none detected | none detected |
| MHI4055 | 15 | 2957 | present | unencapsulated | none detected | none detected |
| MHI4057 | 15 | 2957 | present | unencapsulated | none detected | none detected |
| MHI4061 | 15 | 2957 | present | unencapsulated | none detected | none detected |
| MHI4062 | 15 | 2957 | present | unencapsulated | none detected | none detected |
| MHI4063 | 15 | 2957 | present | unencapsulated | none detected | none detected |
| MHI4064 | 15 | 2957 | present | unencapsulated | none detected | none detected |
| MHI4065 | 15 | 2957 | present | unencapsulated | none detected | none detected |
| MHI4066 | 15 | 2957 | present | unencapsulated | none detected | none detected |
| MHI4067 | 15 | 2957 | present | unencapsulated | none detected | none detected |
| MHI4069 | 15 | 2957 | present | unencapsulated | none detected | none detected |

| | | | | | | |
|---|---|---|---|---|---|---|
| MHI4070 | 15 | 2957 | present | unencapsulated | none detected | none detected |
| MHI4071 | 15 | 2957 | present | unencapsulated | none detected | none detected |
| MHI4072 | 15 | 2957 | present | unencapsulated | none detected | none detected |
| MHI4073 | 15 | 2957 | present | unencapsulated | none detected | none detected |
| MHI4075 | 15 | 2957 | present | unencapsulated | none detected | none detected |
| MHI4076 | 15 | 2957 | present | unencapsulated | none detected | none detected |
| MHI4077 | 15 | 2957 | present | unencapsulated | none detected | none detected |
| MHI4078 | 15 | 2957 | present | unencapsulated | none detected | none detected |
| MHI4081 | 15 | 2957 | present | unencapsulated | none detected | none detected |
| MHI4082 | 15 | 2957 | present | unencapsulated | none detected | none detected |
| MHI4084 | 15 | 2957 | present | unencapsulated | none detected | none detected |
| MHI4086 | 15 | 2957 | present | unencapsulated | none detected | none detected |
| MHI4087 | 15 | 2957 | present | unencapsulated | none detected | none detected |
| MHI4088 | 15 | 2957 | present | unencapsulated | none detected | none detected |
| MHI4089 | 15 | 2957 | present | unencapsulated | none detected | none detected |
| MHI3361 | 16 | 395 | present | unencapsulated | none detected | none detected |
| MHI3362 | 16 | 395 | present | unencapsulated | none detected | none detected |
| MHI3363 | 16 | 395 | present | unencapsulated | none detected | none detected |
| MHI3364 | 16 | 395 | present | unencapsulated | none detected | none detected |
| MHI3365 | 16 | 395 | present | unencapsulated | none detected | none detected |
| MHI3366 | 16 | 395 | present | unencapsulated | none detected | none detected |
| MHI3367 | 16 | 395 | present | unencapsulated | none detected | none detected |
| MHI3368 | 16 | 395 | present | unencapsulated | none detected | none detected |
| MHI3369 | 16 | 395 | present | unencapsulated | none detected | none detected |
| MHI3370 | 16 | 395 | present | unencapsulated | none detected | none detected |
| MHI3371 | 16 | 395 | present | unencapsulated | none detected | none detected |
| MHI3372 | 16 | 395 | present | unencapsulated | none detected | none detected |
| MHI3373 | 16 | 395 | present | unencapsulated | none detected | none detected |
| MHI3374 | 16 | 395 | present | unencapsulated | none detected | none detected |
| MHI3376 | 16 | 395 | present | unencapsulated | none detected | none detected |
| MHI3377 | 16 | 395 | present | unencapsulated | none detected | none detected |
| MHI3378 | 16 | 395 | present | unencapsulated | none detected | none detected |
| MHI3379 | 16 | 395 | present | unencapsulated | none detected | none detected |
| MHI3380 | 16 | 395 | present | unencapsulated | none detected | none detected |
| MHI3381 | 16 | 395 | present | unencapsulated | none detected | none detected |
| MHI3382 | 16 | 395 | present | unencapsulated | none detected | none detected |
| MHI3383 | 16 | 395 | present | unencapsulated | none detected | none detected |
| MHI3384 | 16 | 395 | present | unencapsulated | none detected | none detected |
| MHI3385 | 16 | 395 | present | unencapsulated | none detected | none detected |
| MHI3386 | 16 | 395 | present | unencapsulated | none detected | none detected |
| MHI3387 | 16 | 395 | present | unencapsulated | none detected | none detected |
| MHI3388 | 16 | 395 | present | unencapsulated | none detected | none detected |
| MHI3389 | 16 | 395 | present | unencapsulated | none detected | none detected |
| MHI3390 | 16 | 395 | present | unencapsulated | none detected | none detected |
| MHI3610 | 16 | 395 | present | unencapsulated | none detected | none detected |
| MHI3611 | 16 | 395 | present | unencapsulated | none detected | none detected |
| MHI4390 | 16 | 395 | present | unencapsulated | none detected | none detected |
| MHI4090 | 20 | 946 | present | unencapsulated | none detected | none detected |
| MHI4091 | 20 | 946 | present | unencapsulated | none detected | none detected |
| MHI4092 | 20 | 946 | present | unencapsulated | none detected | none detected |
| MHI4093 | 20 | 946 | present | unencapsulated | none detected | none detected |
| MHI4094 | 20 | 946 | present | unencapsulated | none detected | none detected |
| MHI4095 | 20 | 946 | present | unencapsulated | none detected | none detected |
| MHI4096 | 20 | 946 | present | unencapsulated | none detected | none detected |
| MHI4097 | 20 | 946 | present | unencapsulated | none detected | none detected |

| | | | | | | |
|---|---|---|---|---|---|---|
| MHI4098 | 20 | 946 | present | unencapsulated | none detected | none detected |
| MHI4099 | 20 | 946 | present | unencapsulated | none detected | none detected |
| MHI4100 | 20 | 946 | present | unencapsulated | none detected | none detected |
| MHI4101 | 20 | 946 | present | unencapsulated | none detected | none detected |
| MHI4102 | 20 | 946 | present | unencapsulated | none detected | none detected |
| MHI4103 | 20 | 946 | present | unencapsulated | none detected | none detected |
| MHI4104 | 20 | 946 | present | unencapsulated | none detected | none detected |
| MHI4105 | 20 | 946 | present | unencapsulated | none detected | none detected |
| MHI4106 | 20 | 946 | present | unencapsulated | none detected | none detected |
| MHI4107 | 20 | 946 | present | unencapsulated | none detected | none detected |
| MHI4108 | 20 | 946 | present | unencapsulated | none detected | none detected |
| MHI4109 | 20 | 946 | present | unencapsulated | none detected | none detected |
| MHI4110 | 20 | 946 | present | unencapsulated | none detected | none detected |
| MHI4111 | 20 | 946 | present | unencapsulated | none detected | none detected |
| MHI4112 | 20 | 946 | present | unencapsulated | none detected | none detected |
| MHI4113 | 20 | 946 | present | unencapsulated | none detected | none detected |
| MHI4114 | 20 | 946 | present | unencapsulated | none detected | none detected |
| MHI4115 | 20 | 946 | present | unencapsulated | none detected | none detected |
| MHI4116 | 20 | 946 | present | unencapsulated | none detected | none detected |
| MHI4117 | 20 | 946 | present | unencapsulated | none detected | none detected |
| MHI4118 | 20 | 946 | present | unencapsulated | none detected | none detected |
| MHI4119 | 20 | 946 | present | unencapsulated | none detected | none detected |
| MHI4120 | 20 | 946 | present | unencapsulated | none detected | none detected |
| MHI4121 | 20 | 946 | present | unencapsulated | none detected | none detected |
| MHI4122 | 20 | 946 | present | unencapsulated | none detected | none detected |
| MHI4123 | 20 | 946 | present | unencapsulated | none detected | none detected |
| MHI4124 | 20 | 946 | present | unencapsulated | none detected | none detected |
| MHI3401 | 27 | 494 | present | unencapsulated | none detected | none detected |
| MHI3402 | 27 | 494 | present | unencapsulated | none detected | none detected |
| MHI3404 | 27 | 494 | present | unencapsulated | none detected | none detected |
| MHI3405 | 27 | 494 | present | unencapsulated | none detected | none detected |
| MHI3406 | 27 | 494 | present | unencapsulated | none detected | none detected |
| MHI3413 | 27 | 494 | present | unencapsulated | none detected | none detected |
| MHI3415 | 27 | 494 | present | unencapsulated | none detected | none detected |
| MHI3416 | 27 | 494 | present | unencapsulated | none detected | none detected |
| MHI3417 | 27 | 494 | present | unencapsulated | none detected | none detected |
| MHI3418 | 27 | 494 | present | unencapsulated | none detected | none detected |
| MHI3419 | 27 | 494 | present | unencapsulated | none detected | none detected |
| MHI3613 | 27 | 494 | present | unencapsulated | none detected | none detected |
| MHI4394 | 27 | 494 | present | unencapsulated | none detected | none detected |
| MHI3512 | 27 | 683 | present | unencapsulated | none detected | TEM |
| MHI3515 | 27 | 683 | present | unencapsulated | none detected | TEM |
| MHI3518 | 27 | 683 | present | unencapsulated | none detected | TEM |
| MHI3519 | 27 | 683 | present | unencapsulated | none detected | TEM |
| MHI3525 | 27 | 683 | present | unencapsulated | none detected | TEM |
| MHI3526 | 27 | 683 | present | unencapsulated | none detected | TEM |
| MHI3527 | 27 | 683 | present | unencapsulated | none detected | TEM |
| MHI3529 | 27 | 683 | present | unencapsulated | none detected | TEM |
| MHI3532 | 27 | 683 | present | unencapsulated | none detected | TEM |
| MHI3533 | 27 | 683 | present | unencapsulated | none detected | TEM |
| MHI3535 | 27 | 683 | present | unencapsulated | none detected | TEM |
| MHI3538 | 27 | 683 | present | unencapsulated | none detected | TEM |
| MHI3540 | 27 | 683 | present | unencapsulated | none detected | TEM |

| | | | | | | |
|---|---|---|---|---|---|---|
| MHI3541 | 27 | 683 | present | unencapsulated | none detected | TEM |
| MHI3511 | 27 | 1034 | present | unencapsulated | none detected | none detected |
| MHI3514 | 27 | 1034 | present | unencapsulated | none detected | none detected |
| MHI3516 | 27 | 1034 | present | unencapsulated | none detected | none detected |
| MHI3517 | 27 | 1034 | present | unencapsulated | none detected | none detected |
| MHI3521 | 27 | 1034 | present | unencapsulated | none detected | none detected |
| MHI3522 | 27 | 1034 | present | unencapsulated | none detected | none detected |
| MHI3530 | 27 | 1034 | present | unencapsulated | none detected | none detected |
| MHI3534 | 27 | 1034 | present | unencapsulated | none detected | none detected |
| MHI3536 | 27 | 1034 | present | unencapsulated | none detected | none detected |
| MHI3537 | 27 | 1034 | present | unencapsulated | none detected | none detected |
| MHI3539 | 27 | 1034 | present | unencapsulated | none detected | none detected |
| MHI4396 | 27 | 1034 | present | unencapsulated | none detected | none detected |
| MHI3391 | 27 | 2954 | present | unencapsulated | none detected | none detected |
| MHI3392 | 27 | 2954 | present | unencapsulated | none detected | none detected |
| MHI3393 | 27 | 2954 | present | unencapsulated | none detected | none detected |
| MHI3394 | 27 | 2954 | present | unencapsulated | none detected | none detected |
| MHI3395 | 27 | 2954 | present | unencapsulated | none detected | none detected |
| MHI3396 | 27 | 2954 | present | unencapsulated | none detected | none detected |
| MHI3397 | 27 | 2954 | present | unencapsulated | none detected | none detected |
| MHI3398 | 27 | 2954 | present | unencapsulated | none detected | none detected |
| MHI3399 | 27 | 2954 | present | unencapsulated | none detected | none detected |
| MHI3400 | 27 | 2954 | present | unencapsulated | none detected | none detected |
| MHI3403 | 27 | 2954 | present | unencapsulated | none detected | none detected |
| MHI3407 | 27 | 2954 | present | unencapsulated | none detected | none detected |
| MHI3408 | 27 | 2954 | present | unencapsulated | none detected | none detected |
| MHI3409 | 27 | 2954 | present | unencapsulated | none detected | none detected |
| MHI3410 | 27 | 2954 | present | unencapsulated | none detected | none detected |
| MHI3411 | 27 | 2954 | present | unencapsulated | none detected | none detected |
| MHI3412 | 27 | 2954 | present | unencapsulated | none detected | none detected |
| MHI3414 | 27 | 2954 | present | unencapsulated | none detected | none detected |
| MHI3420 | 27 | 2954 | present | unencapsulated | none detected | none detected |
| MHI4392 | 27 | 2954 | present | unencapsulated | none detected | none detected |
| MHI3542 | 29 | 199 | present | unencapsulated | none detected | none detected |
| MHI3544 | 29 | 199 | present | unencapsulated | none detected | none detected |
| MHI3547 | 29 | 199 | present | unencapsulated | none detected | none detected |
| MHI3548 | 29 | 199 | present | unencapsulated | none detected | none detected |
| MHI3550 | 29 | 199 | present | unencapsulated | none detected | none detected |
| MHI3552 | 29 | 199 | present | unencapsulated | none detected | none detected |
| MHI3553 | 29 | 199 | present | unencapsulated | none detected | none detected |
| MHI3555 | 29 | 199 | present | unencapsulated | none detected | none detected |
| MHI3556 | 29 | 199 | present | unencapsulated | none detected | none detected |
| MHI3558 | 29 | 199 | present | unencapsulated | none detected | none detected |
| MHI3561 | 29 | 199 | present | unencapsulated | none detected | none detected |
| MHI3563 | 29 | 199 | present | unencapsulated | none detected | none detected |
| MHI3564 | 29 | 199 | present | unencapsulated | none detected | none detected |
| MHI3565 | 29 | 199 | present | unencapsulated | none detected | none detected |
| MHI3566 | 29 | 199 | present | unencapsulated | none detected | none detected |
| MHI3567 | 29 | 199 | present | unencapsulated | none detected | none detected |
| MHI3568 | 29 | 199 | present | unencapsulated | none detected | none detected |
| MHI3569 | 29 | 199 | present | unencapsulated | none detected | none detected |
| MHI3572 | 29 | 199 | present | unencapsulated | none detected | none detected |
| MHI3573 | 29 | 199 | present | unencapsulated | none detected | none detected |

| | | | | | | |
|---|---|---|---|---|---|---|
| MHI4401 | 29 | 199 | present | unencapsulated | none detected | none detected |
| MHI4402 | 29 | 199 | present | unencapsulated | none detected | none detected |
| MHI4403 | 29 | 199 | present | unencapsulated | none detected | none detected |
| MHI3421 | 29 | 266 | present | unencapsulated | none detected | none detected |
| MHI3422 | 29 | 266 | present | unencapsulated | none detected | none detected |
| MHI3423 | 29 | 266 | present | unencapsulated | none detected | none detected |
| MHI3424 | 29 | 266 | present | unencapsulated | none detected | none detected |
| MHI3425 | 29 | 266 | present | unencapsulated | none detected | none detected |
| MHI3426 | 29 | 266 | present | unencapsulated | none detected | none detected |
| MHI3427 | 29 | 266 | present | unencapsulated | none detected | none detected |
| MHI3429 | 29 | 266 | present | unencapsulated | none detected | none detected |
| MHI3430 | 29 | 266 | present | unencapsulated | none detected | none detected |
| MHI3431 | 29 | 266 | present | unencapsulated | none detected | none detected |
| MHI3433 | 29 | 266 | present | unencapsulated | none detected | none detected |
| MHI3434 | 29 | 266 | present | unencapsulated | none detected | none detected |
| MHI3435 | 29 | 266 | present | unencapsulated | none detected | none detected |
| MHI3436 | 29 | 266 | present | unencapsulated | none detected | none detected |
| MHI3437 | 29 | 266 | present | unencapsulated | none detected | none detected |
| MHI3438 | 29 | 266 | present | unencapsulated | none detected | none detected |
| MHI3440 | 29 | 266 | present | unencapsulated | none detected | none detected |
| MHI3441 | 29 | 266 | present | unencapsulated | none detected | none detected |
| MHI3442 | 29 | 266 | present | unencapsulated | none detected | none detected |
| MHI3443 | 29 | 266 | present | unencapsulated | none detected | none detected |
| MHI3444 | 29 | 266 | present | unencapsulated | none detected | none detected |
| MHI3445 | 29 | 266 | present | unencapsulated | none detected | none detected |
| MHI3446 | 29 | 266 | present | unencapsulated | none detected | none detected |
| MHI3447 | 29 | 266 | present | unencapsulated | none detected | none detected |
| MHI3449 | 29 | 266 | present | unencapsulated | none detected | none detected |
| MHI3450 | 29 | 266 | present | unencapsulated | none detected | none detected |
| MHI3614 | 29 | 266 | present | unencapsulated | none detected | none detected |
| MHI3615 | 29 | 266 | present | unencapsulated | none detected | none detected |
| MHI4398 | 29 | 266 | present | unencapsulated | none detected | none detected |
| MHI4399 | 29 | 266 | present | unencapsulated | none detected | none detected |
| MHI4400 | 29 | 266 | present | unencapsulated | none detected | none detected |
| MHI3546 | 29 | 536 | present | unencapsulated | none detected | none detected |
| MHI3551 | 29 | 536 | present | unencapsulated | none detected | none detected |
| MHI3557 | 29 | 536 | present | unencapsulated | none detected | none detected |
| MHI3559 | 29 | 536 | present | unencapsulated | none detected | none detected |
| MHI3562 | 29 | 536 | present | unencapsulated | none detected | none detected |
| MHI3570 | 29 | 536 | present | unencapsulated | none detected | none detected |
| MHI3543 | 29 | 598 | present | unencapsulated | none detected | none detected |
| MHI3545 | 29 | 598 | present | unencapsulated | none detected | none detected |
| MHI3571 | 29 | 598 | present | unencapsulated | none detected | none detected |
| MHI3428 | 29 | 1030 | present | unencapsulated | none detected | none detected |
| MHI3432 | 29 | 1030 | present | unencapsulated | none detected | none detected |
| MHI3448 | 29 | 1030 | present | unencapsulated | none detected | none detected |
| MHI4125 | 36 | 925 | present | unencapsulated | none detected | none detected |
| MHI4126 | 36 | 925 | present | unencapsulated | none detected | none detected |
| MHI4127 | 36 | 925 | present | unencapsulated | none detected | none detected |
| MHI4129 | 36 | 925 | present | unencapsulated | none detected | none detected |
| MHI4130 | 36 | 925 | present | unencapsulated | none detected | none detected |
| MHI4132 | 36 | 925 | present | unencapsulated | none detected | none detected |
| MHI4133 | 36 | 925 | present | unencapsulated | none detected | none detected |

| | | | | | | |
|---|---|---|---|---|---|---|
| MHI4138 | 36 | 925 | present | unencapsulated | none detected | none detected |
| MHI4139 | 36 | 925 | present | unencapsulated | none detected | none detected |
| MHI4140 | 36 | 925 | present | unencapsulated | none detected | none detected |
| MHI4142 | 36 | 925 | present | unencapsulated | none detected | none detected |
| MHI4143 | 36 | 925 | present | unencapsulated | none detected | none detected |
| MHI4146 | 36 | 925 | present | unencapsulated | none detected | none detected |
| MHI4147 | 36 | 925 | present | unencapsulated | none detected | none detected |
| MHI4149 | 36 | 925 | present | unencapsulated | none detected | none detected |
| MHI4152 | 36 | 925 | present | unencapsulated | none detected | none detected |
| MHI4158 | 36 | 925 | present | unencapsulated | none detected | none detected |
| MHI4160 | 39 | 159 | present | unencapsulated | none detected | none detected |
| MHI4161 | 39 | 159 | present | unencapsulated | none detected | none detected |
| MHI4162 | 39 | 159 | present | unencapsulated | none detected | none detected |
| MHI4167 | 39 | 159 | present | unencapsulated | none detected | none detected |
| MHI4168 | 39 | 159 | present | unencapsulated | none detected | none detected |
| MHI4169 | 39 | 159 | present | unencapsulated | none detected | none detected |
| MHI4170 | 39 | 159 | present | unencapsulated | none detected | none detected |
| MHI4171 | 39 | 159 | present | unencapsulated | none detected | none detected |
| MHI4172 | 39 | 159 | present | unencapsulated | none detected | none detected |
| MHI4173 | 39 | 159 | present | unencapsulated | none detected | none detected |
| MHI4184 | 39 | 159 | present | unencapsulated | none detected | none detected |
| MHI4191 | 39 | 159 | present | unencapsulated | none detected | none detected |
| MHI4193 | 39 | 159 | present | unencapsulated | none detected | none detected |
| MHI3451 | 43 | 836 | present | unencapsulated | none detected | none detected |
| MHI3452 | 43 | 836 | present | unencapsulated | none detected | none detected |
| MHI3453 | 43 | 836 | present | unencapsulated | none detected | none detected |
| MHI3454 | 43 | 836 | present | unencapsulated | none detected | none detected |
| MHI3455 | 43 | 836 | present | unencapsulated | none detected | none detected |
| MHI3456 | 43 | 836 | present | unencapsulated | none detected | none detected |
| MHI3457 | 43 | 836 | present | unencapsulated | none detected | none detected |
| MHI3458 | 43 | 836 | present | unencapsulated | none detected | none detected |
| MHI3459 | 43 | 836 | present | unencapsulated | none detected | none detected |
| MHI3460 | 43 | 836 | present | unencapsulated | none detected | none detected |
| MHI3461 | 43 | 836 | present | unencapsulated | none detected | none detected |
| MHI3462 | 43 | 836 | present | unencapsulated | none detected | none detected |
| MHI3463 | 43 | 836 | present | unencapsulated | none detected | none detected |
| MHI3464 | 43 | 836 | present | unencapsulated | none detected | none detected |
| MHI3465 | 43 | 836 | present | unencapsulated | none detected | none detected |
| MHI3466 | 43 | 836 | present | unencapsulated | none detected | none detected |
| MHI3467 | 43 | 836 | present | unencapsulated | none detected | none detected |
| MHI3468 | 43 | 836 | present | unencapsulated | none detected | none detected |
| MHI3469 | 43 | 836 | present | unencapsulated | none detected | none detected |
| MHI3470 | 43 | 836 | present | unencapsulated | none detected | none detected |
| MHI3471 | 43 | 836 | present | unencapsulated | none detected | none detected |
| MHI3472 | 43 | 836 | present | unencapsulated | none detected | none detected |
| MHI3473 | 43 | 836 | present | unencapsulated | none detected | none detected |
| MHI3474 | 43 | 836 | present | unencapsulated | none detected | none detected |
| MHI3475 | 43 | 836 | present | unencapsulated | none detected | none detected |
| MHI3476 | 43 | 836 | present | unencapsulated | none detected | none detected |
| MHI3477 | 43 | 836 | present | unencapsulated | none detected | none detected |
| MHI3478 | 43 | 836 | present | unencapsulated | none detected | none detected |
| MHI3479 | 43 | 836 | present | unencapsulated | none detected | none detected |
| MHI3480 | 43 | 836 | present | unencapsulated | none detected | none detected |

| | | | | | | |
|---|---|---|---|---|---|---|
| MHI3616 | 43 | 836 | present | unencapsulated | none detected | none detected |
| MHI3617 | 43 | 836 | present | unencapsulated | none detected | none detected |
| MHI4404 | 43 | 836 | present | unencapsulated | none detected | none detected |
| MHI4405 | 43 | 836 | present | unencapsulated | none detected | none detected |
| MHI4406 | 43 | 836 | present | unencapsulated | none detected | none detected |
| MHI3481 | 46 | 266 | present | unencapsulated | none detected | none detected |
| MHI3482 | 46 | 266 | present | unencapsulated | none detected | none detected |
| MHI3483 | 46 | 266 | present | unencapsulated | none detected | none detected |
| MHI3484 | 46 | 266 | present | unencapsulated | none detected | none detected |
| MHI3485 | 46 | 266 | present | unencapsulated | none detected | none detected |
| MHI3486 | 46 | 266 | present | unencapsulated | none detected | none detected |
| MHI3487 | 46 | 266 | present | unencapsulated | none detected | none detected |
| MHI3488 | 46 | 266 | present | unencapsulated | none detected | none detected |
| MHI3489 | 46 | 266 | present | unencapsulated | none detected | none detected |
| MHI3490 | 46 | 266 | present | unencapsulated | none detected | none detected |
| MHI3491 | 46 | 266 | present | unencapsulated | none detected | none detected |
| MHI3492 | 46 | 266 | present | unencapsulated | none detected | none detected |
| MHI3493 | 46 | 266 | present | unencapsulated | none detected | none detected |
| MHI3494 | 46 | 266 | present | unencapsulated | none detected | none detected |
| MHI3495 | 46 | 266 | present | unencapsulated | none detected | none detected |
| MHI3496 | 46 | 266 | present | unencapsulated | none detected | none detected |
| MHI3497 | 46 | 266 | present | unencapsulated | none detected | none detected |
| MHI3498 | 46 | 266 | present | unencapsulated | none detected | none detected |
| MHI3499 | 46 | 266 | present | unencapsulated | none detected | none detected |
| MHI3500 | 46 | 266 | present | unencapsulated | none detected | none detected |
| MHI3501 | 46 | 266 | present | unencapsulated | none detected | none detected |
| MHI3502 | 46 | 266 | present | unencapsulated | none detected | none detected |
| MHI3503 | 46 | 266 | present | unencapsulated | none detected | none detected |
| MHI3504 | 46 | 266 | present | unencapsulated | none detected | none detected |
| MHI3505 | 46 | 266 | present | unencapsulated | none detected | none detected |
| MHI3506 | 46 | 266 | present | unencapsulated | none detected | none detected |
| MHI3507 | 46 | 266 | present | unencapsulated | none detected | none detected |
| MHI3508 | 46 | 266 | present | unencapsulated | none detected | none detected |
| MHI3509 | 46 | 266 | present | unencapsulated | none detected | none detected |
| MHI3510 | 46 | 266 | present | unencapsulated | none detected | none detected |
| MHI3618 | 46 | 266 | present | unencapsulated | none detected | none detected |
| MHI3619 | 46 | 266 | present | unencapsulated | none detected | none detected |
| MHI4407 | 46 | 266 | present | unencapsulated | none detected | none detected |
| MHI4408 | 46 | 266 | present | unencapsulated | none detected | none detected |
| MHI4409 | 46 | 266 | present | unencapsulated | none detected | none detected |
| MHI3620 | 52 | 105 | present | unencapsulated | none detected | none detected |
| MHI3621 | 52 | 105 | present | unencapsulated | none detected | none detected |
| MHI3622 | 52 | 105 | present | unencapsulated | none detected | none detected |
| MHI3623 | 52 | 105 | present | unencapsulated | none detected | none detected |
| MHI3624 | 52 | 105 | present | unencapsulated | none detected | none detected |
| MHI3626 | 52 | 105 | present | unencapsulated | none detected | none detected |
| MHI3627 | 52 | 105 | present | unencapsulated | none detected | none detected |
| MHI3628 | 52 | 105 | present | unencapsulated | none detected | none detected |
| MHI3629 | 52 | 105 | present | unencapsulated | none detected | none detected |
| MHI3630 | 52 | 105 | present | unencapsulated | none detected | none detected |
| MHI3631 | 52 | 105 | present | unencapsulated | none detected | none detected |
| MHI3632 | 52 | 105 | present | unencapsulated | none detected | none detected |
| MHI3633 | 52 | 105 | present | unencapsulated | none detected | none detected |

| | | | | | | |
|---|---|---|---|---|---|---|
| MHI3634 | 52 | 105 | present | unencapsulated | none detected | none detected |
| MHI3635 | 52 | 105 | present | unencapsulated | none detected | none detected |
| MHI3636 | 52 | 105 | present | unencapsulated | none detected | none detected |
| MHI3637 | 52 | 105 | present | unencapsulated | none detected | none detected |
| MHI3638 | 52 | 105 | present | unencapsulated | none detected | none detected |
| MHI3639 | 52 | 105 | present | unencapsulated | none detected | none detected |
| MHI3640 | 52 | 105 | present | unencapsulated | none detected | none detected |
| MHI3641 | 52 | 105 | present | unencapsulated | none detected | none detected |
| MHI3642 | 52 | 105 | present | unencapsulated | none detected | none detected |
| MHI3643 | 52 | 105 | present | unencapsulated | none detected | none detected |
| MHI3644 | 52 | 105 | present | unencapsulated | none detected | none detected |
| MHI3645 | 52 | 105 | present | unencapsulated | none detected | none detected |
| MHI3646 | 52 | 105 | present | unencapsulated | none detected | none detected |
| MHI3647 | 52 | 105 | present | unencapsulated | none detected | none detected |
| MHI3648 | 52 | 105 | present | unencapsulated | none detected | none detected |
| MHI3649 | 52 | 105 | present | unencapsulated | none detected | none detected |
| MHI3650 | 52 | 105 | present | unencapsulated | none detected | none detected |
| MHI3651 | 52 | 105 | present | unencapsulated | none detected | none detected |
| MHI4410 | 52 | 105 | present | unencapsulated | none detected | none detected |
| MHI4411 | 52 | 105 | present | unencapsulated | none detected | none detected |
| MHI4194 | 56 | 334 | present | unencapsulated | none detected | none detected |
| MHI4195 | 56 | 334 | present | unencapsulated | none detected | none detected |
| MHI4198 | 56 | 334 | present | unencapsulated | none detected | none detected |
| MHI4210 | 56 | 334 | present | unencapsulated | none detected | none detected |
| MHI4211 | 56 | 334 | present | unencapsulated | none detected | none detected |
| MHI4212 | 56 | 334 | present | unencapsulated | none detected | none detected |
| MHI4213 | 56 | 334 | present | unencapsulated | none detected | none detected |
| MHI4214 | 56 | 334 | present | unencapsulated | none detected | none detected |
| MHI4215 | 56 | 334 | present | unencapsulated | none detected | none detected |
| MHI4216 | 56 | 334 | present | unencapsulated | none detected | none detected |
| MHI4217 | 56 | 334 | present | unencapsulated | none detected | none detected |
| MHI4218 | 56 | 334 | present | unencapsulated | none detected | none detected |
| MHI4219 | 56 | 334 | present | unencapsulated | none detected | none detected |
| MHI4220 | 56 | 334 | present | unencapsulated | none detected | none detected |
| MHI4221 | 56 | 334 | present | unencapsulated | none detected | none detected |
| MHI4222 | 56 | 334 | present | unencapsulated | none detected | none detected |
| MHI4223 | 56 | 334 | present | unencapsulated | none detected | none detected |
| MHI4224 | 56 | 334 | present | unencapsulated | none detected | none detected |
| MHI4225 | 56 | 334 | present | unencapsulated | none detected | none detected |
| MHI4226 | 56 | 334 | present | unencapsulated | none detected | none detected |
| MHI4227 | 57 | 161 | present | unencapsulated | none detected | none detected |
| MHI4228 | 57 | 161 | present | unencapsulated | none detected | none detected |
| MHI4229 | 57 | 161 | present | unencapsulated | none detected | none detected |
| MHI4230 | 57 | 161 | present | unencapsulated | none detected | none detected |
| MHI4231 | 57 | 161 | present | unencapsulated | none detected | none detected |
| MHI4232 | 57 | 161 | present | unencapsulated | none detected | none detected |
| MHI4233 | 57 | 161 | present | unencapsulated | none detected | none detected |
| MHI4234 | 57 | 161 | present | unencapsulated | none detected | none detected |
| MHI4235 | 57 | 161 | present | unencapsulated | none detected | none detected |
| MHI4236 | 57 | 161 | present | unencapsulated | none detected | none detected |
| MHI4237 | 57 | 161 | present | unencapsulated | none detected | none detected |
| MHI4238 | 57 | 161 | present | unencapsulated | none detected | none detected |
| MHI4239 | 57 | 161 | present | unencapsulated | none detected | none detected |

| | | | | | | |
|---|---|---|---|---|---|---|
| MHI4240 | 57 | 161 | present | unencapsulated | none detected | none detected |
| MHI4241 | 57 | 161 | present | unencapsulated | none detected | none detected |
| MHI4242 | 57 | 161 | present | unencapsulated | none detected | none detected |
| MHI4243 | 57 | 161 | present | unencapsulated | none detected | none detected |
| MHI4244 | 57 | 161 | present | unencapsulated | none detected | none detected |
| MHI4245 | 57 | 161 | present | unencapsulated | none detected | none detected |
| MHI4246 | 57 | 161 | present | unencapsulated | none detected | none detected |
| MHI4247 | 57 | 161 | present | unencapsulated | none detected | none detected |
| MHI4248 | 57 | 161 | present | unencapsulated | none detected | none detected |
| MHI4249 | 57 | 161 | present | unencapsulated | none detected | none detected |
| MHI4250 | 57 | 161 | present | unencapsulated | none detected | none detected |
| MHI4251 | 57 | 161 | present | unencapsulated | none detected | none detected |
| MHI4252 | 57 | 161 | present | unencapsulated | none detected | none detected |
| MHI4253 | 57 | 161 | present | unencapsulated | none detected | none detected |
| MHI4254 | 57 | 161 | present | unencapsulated | none detected | none detected |
| MHI4255 | 57 | 161 | present | unencapsulated | none detected | none detected |
| MHI4256 | 57 | 161 | present | unencapsulated | none detected | none detected |
| MHI4257 | 57 | 161 | present | unencapsulated | none detected | none detected |
| MHI4258 | 57 | 161 | present | unencapsulated | none detected | none detected |
| MHI4259 | 57 | 161 | present | unencapsulated | none detected | none detected |
| MHI4260 | 57 | 161 | present | unencapsulated | none detected | none detected |
| MHI4261 | 57 | 161 | present | unencapsulated | none detected | none detected |
| MHI4264 | 59 | 280 | present | unencapsulated | none detected | TEM |
| MHI4265 | 59 | 280 | present | unencapsulated | none detected | TEM |
| MHI4272 | 59 | 280 | present | unencapsulated | none detected | TEM |
| MHI4274 | 59 | 280 | present | unencapsulated | none detected | none detected |
| MHI4277 | 59 | 280 | present | unencapsulated | none detected | none detected |
| MHI4280 | 59 | 280 | present | unencapsulated | none detected | none detected |
| MHI4285 | 59 | 280 | present | unencapsulated | none detected | none detected |
| MHI4287 | 59 | 280 | present | unencapsulated | none detected | none detected |
| MHI4289 | 59 | 280 | present | unencapsulated | none detected | none detected |
| MHI4292 | 59 | 280 | present | unencapsulated | none detected | none detected |
| MHI4293 | 59 | 280 | present | unencapsulated | none detected | none detected |
| MHI4262 | 59 | 1409 | present | unencapsulated | none detected | none detected |
| MHI4263 | 59 | 1409 | present | unencapsulated | none detected | none detected |
| MHI4267 | 59 | 1409 | present | unencapsulated | none detected | TEM |
| MHI4269 | 59 | 1409 | present | unencapsulated | none detected | none detected |
| MHI4270 | 59 | 1409 | present | unencapsulated | none detected | none detected |
| MHI4271 | 59 | 1409 | present | unencapsulated | none detected | none detected |
| MHI4275 | 59 | 1409 | present | unencapsulated | none detected | none detected |
| MHI4276 | 59 | 1409 | present | unencapsulated | none detected | none detected |
| MHI4278 | 59 | 1409 | present | unencapsulated | none detected | none detected |
| MHI4279 | 59 | 1409 | present | unencapsulated | none detected | none detected |
| MHI4281 | 59 | 1409 | present | unencapsulated | none detected | none detected |
| MHI4282 | 59 | 1409 | present | unencapsulated | none detected | none detected |
| MHI4283 | 59 | 1409 | present | unencapsulated | none detected | none detected |
| MHI4284 | 59 | 1409 | present | unencapsulated | none detected | none detected |
| MHI4286 | 59 | 1409 | present | unencapsulated | none detected | none detected |
| MHI4288 | 59 | 1409 | present | unencapsulated | none detected | none detected |
| MHI4290 | 59 | 1409 | present | unencapsulated | none detected | none detected |
| MHI4291 | 59 | 1409 | present | unencapsulated | none detected | none detected |
| MHI4294 | 59 | 1409 | present | unencapsulated | none detected | none detected |
| MHI4295 | 59 | 1409 | present | unencapsulated | none detected | none detected |

| | | | | | | |
|---|---|---|---|---|---|---|
| MHI4296 | 59 | 1409 | present | unencapsulated | none detected | none detected |
| MHI3652 | 61 | 568 | present | unencapsulated | none detected | none detected |
| MHI3653 | 61 | 568 | present | unencapsulated | none detected | none detected |
| MHI3654 | 61 | 568 | present | unencapsulated | none detected | none detected |
| MHI3659 | 61 | 568 | present | unencapsulated | none detected | none detected |
| MHI3664 | 61 | 568 | present | unencapsulated | none detected | none detected |
| MHI3665 | 61 | 568 | present | unencapsulated | none detected | none detected |
| MHI3670 | 61 | 568 | present | unencapsulated | none detected | none detected |
| MHI3672 | 61 | 568 | present | unencapsulated | none detected | none detected |
| MHI3673 | 61 | 568 | present | unencapsulated | none detected | none detected |
| MHI3674 | 61 | 568 | present | unencapsulated | none detected | none detected |
| MHI3675 | 61 | 568 | present | unencapsulated | none detected | none detected |
| MHI3682 | 61 | 568 | present | unencapsulated | none detected | none detected |
| MHI3655 | 61 | 1012 | present | unencapsulated | none detected | none detected |
| MHI3656 | 61 | 1012 | present | unencapsulated | none detected | none detected |
| MHI3657 | 61 | 1012 | present | unencapsulated | none detected | none detected |
| MHI3658 | 61 | 1012 | present | unencapsulated | none detected | none detected |
| MHI3660 | 61 | 1012 | present | unencapsulated | none detected | none detected |
| MHI3661 | 61 | 1012 | present | unencapsulated | none detected | none detected |
| MHI3662 | 61 | 1012 | present | unencapsulated | none detected | none detected |
| MHI3663 | 61 | 1012 | present | unencapsulated | none detected | none detected |
| MHI3667 | 61 | 1012 | present | unencapsulated | none detected | none detected |
| MHI3668 | 61 | 1012 | present | unencapsulated | none detected | none detected |
| MHI3669 | 61 | 1012 | present | unencapsulated | none detected | none detected |
| MHI3671 | 61 | 1012 | present | unencapsulated | none detected | none detected |
| MHI3676 | 61 | 1012 | present | unencapsulated | none detected | none detected |
| MHI3677 | 61 | 1012 | present | unencapsulated | none detected | none detected |
| MHI3678 | 61 | 1012 | present | unencapsulated | none detected | none detected |
| MHI3679 | 61 | 1012 | present | unencapsulated | none detected | none detected |
| MHI3680 | 61 | 1012 | present | unencapsulated | none detected | none detected |
| MHI3681 | 61 | 1012 | present | unencapsulated | none detected | none detected |
| MHI3683 | 61 | 1012 | present | unencapsulated | none detected | none detected |
| MHI4297 | 64 | 2958 | present | unencapsulated | none detected | none detected |
| MHI4298 | 64 | 2958 | present | unencapsulated | none detected | none detected |
| MHI4299 | 64 | 2958 | present | unencapsulated | none detected | none detected |
| MHI4300 | 64 | 2958 | present | unencapsulated | none detected | none detected |
| MHI4301 | 64 | 2958 | present | unencapsulated | none detected | none detected |
| MHI4302 | 64 | 2958 | present | unencapsulated | none detected | none detected |
| MHI4303 | 64 | 2958 | present | unencapsulated | none detected | none detected |
| MHI4304 | 64 | 2958 | present | unencapsulated | none detected | none detected |
| MHI4305 | 64 | 2958 | present | unencapsulated | none detected | none detected |
| MHI4306 | 64 | 2958 | present | unencapsulated | none detected | none detected |
| MHI4307 | 64 | 2958 | present | unencapsulated | none detected | none detected |
| MHI4308 | 64 | 2958 | present | unencapsulated | none detected | none detected |
| MHI4309 | 64 | 2958 | present | unencapsulated | none detected | none detected |
| MHI4310 | 64 | 2958 | present | unencapsulated | none detected | none detected |
| MHI4311 | 64 | 2958 | present | unencapsulated | none detected | none detected |
| MHI4312 | 64 | 2958 | present | unencapsulated | none detected | none detected |
| MHI4313 | 64 | 2958 | present | unencapsulated | none detected | none detected |
| MHI4314 | 64 | 2958 | present | unencapsulated | none detected | none detected |
| MHI4315 | 64 | 2958 | present | unencapsulated | none detected | none detected |
| MHI4316 | 64 | 2958 | present | unencapsulated | none detected | none detected |
| MHI4317 | 64 | 2958 | present | unencapsulated | none detected | none detected |

| | | | | | | |
|---|---|---|---|---|---|---|
| MHI4318 | 64 | 2958 | present | unencapsulated | none detected | none detected |
| MHI4319 | 64 | 2958 | present | unencapsulated | none detected | none detected |
| MHI4320 | 64 | 2958 | present | unencapsulated | none detected | none detected |
| MHI4321 | 64 | 2958 | present | unencapsulated | none detected | none detected |
| MHI4322 | 64 | 2958 | present | unencapsulated | none detected | none detected |
| MHI4323 | 64 | 2958 | present | unencapsulated | none detected | none detected |
| MHI4324 | 64 | 2958 | present | unencapsulated | none detected | none detected |
| MHI4325 | 64 | 2958 | present | unencapsulated | none detected | none detected |
| MHI4326 | 64 | 2958 | present | unencapsulated | none detected | none detected |
| MHI4327 | 64 | 2958 | present | unencapsulated | none detected | none detected |
| MHI4328 | 64 | 2958 | present | unencapsulated | none detected | none detected |
| MHI4329 | 64 | 2958 | present | unencapsulated | none detected | none detected |
| MHI4330 | 64 | 2958 | present | unencapsulated | none detected | none detected |
| MHI4331 | 64 | 2958 | present | unencapsulated | none detected | none detected |
| MHI4332 | 68 | 264 | present | unencapsulated | none detected | none detected |
| MHI4333 | 68 | 264 | present | unencapsulated | none detected | none detected |
| MHI4334 | 68 | 264 | present | unencapsulated | none detected | none detected |
| MHI4335 | 68 | 264 | present | unencapsulated | none detected | none detected |
| MHI4336 | 68 | 264 | present | unencapsulated | none detected | none detected |
| MHI4337 | 68 | 264 | present | unencapsulated | none detected | none detected |
| MHI4339 | 68 | 264 | present | unencapsulated | none detected | none detected |
| MHI4340 | 68 | 264 | present | unencapsulated | none detected | none detected |
| MHI4341 | 68 | 264 | present | unencapsulated | none detected | none detected |
| MHI4342 | 68 | 264 | present | unencapsulated | none detected | none detected |
| MHI4345 | 68 | 264 | present | unencapsulated | none detected | none detected |
| MHI4346 | 68 | 264 | present | unencapsulated | none detected | none detected |
| MHI4347 | 68 | 264 | present | unencapsulated | none detected | none detected |
| MHI4348 | 68 | 264 | present | unencapsulated | none detected | none detected |
| MHI4349 | 68 | 264 | present | unencapsulated | none detected | none detected |
| MHI4350 | 68 | 264 | present | unencapsulated | none detected | none detected |
| MHI4351 | 68 | 264 | present | unencapsulated | none detected | none detected |
| MHI4352 | 68 | 264 | present | unencapsulated | none detected | none detected |
| MHI4353 | 68 | 264 | present | unencapsulated | none detected | none detected |
| MHI4354 | 68 | 264 | present | unencapsulated | none detected | none detected |
| MHI4357 | 68 | 264 | present | unencapsulated | none detected | none detected |
| MHI4358 | 68 | 264 | present | unencapsulated | none detected | none detected |
| MHI4359 | 68 | 264 | present | unencapsulated | none detected | none detected |
| MHI4360 | 68 | 264 | present | unencapsulated | none detected | none detected |
| MHI4361 | 68 | 264 | present | unencapsulated | none detected | none detected |
| MHI4362 | 68 | 264 | present | unencapsulated | none detected | none detected |
| MHI4363 | 68 | 264 | present | unencapsulated | none detected | none detected |
| MHI4364 | 68 | 264 | present | unencapsulated | none detected | none detected |
| MHI4365 | 68 | 264 | present | unencapsulated | none detected | none detected |
| MHI3685 | 73 | 1030 | present | unencapsulated | none detected | none detected |
| MHI3686 | 73 | 1030 | present | unencapsulated | none detected | none detected |
| MHI3687 | 73 | 1030 | present | unencapsulated | none detected | none detected |
| MHI3688 | 73 | 1030 | present | unencapsulated | none detected | none detected |
| MHI3689 | 73 | 1030 | present | unencapsulated | none detected | none detected |
| MHI3690 | 73 | 1030 | present | unencapsulated | none detected | none detected |
| MHI3691 | 73 | 1030 | present | unencapsulated | none detected | none detected |
| MHI3692 | 73 | 1030 | present | unencapsulated | none detected | none detected |
| MHI3693 | 73 | 1030 | present | unencapsulated | none detected | none detected |
| MHI3694 | 73 | 1030 | present | unencapsulated | none detected | none detected |

| | | | | | | |
|---|---|---|---|---|---|---|
| MHI3695 | 73 | 1030 | present | unencapsulated | none detected | none detected |
| MHI3696 | 73 | 1030 | present | unencapsulated | none detected | none detected |
| MHI3697 | 73 | 1030 | present | unencapsulated | none detected | none detected |
| MHI3698 | 73 | 1030 | present | unencapsulated | none detected | none detected |
| MHI3699 | 73 | 1030 | present | unencapsulated | none detected | none detected |
| MHI3700 | 73 | 1030 | present | unencapsulated | none detected | none detected |
| MHI3701 | 73 | 1030 | present | unencapsulated | none detected | none detected |
| MHI3702 | 73 | 1030 | present | unencapsulated | none detected | none detected |
| MHI3703 | 73 | 1030 | present | unencapsulated | none detected | none detected |
| MHI3704 | 73 | 1030 | present | unencapsulated | none detected | none detected |
| MHI3705 | 73 | 1030 | present | unencapsulated | none detected | none detected |
| MHI3706 | 73 | 1030 | present | unencapsulated | none detected | none detected |
| MHI3707 | 73 | 1030 | present | unencapsulated | none detected | none detected |
| MHI3708 | 73 | 1030 | present | unencapsulated | none detected | none detected |
| MHI3709 | 73 | 1030 | present | unencapsulated | none detected | none detected |
| MHI3710 | 73 | 1030 | present | unencapsulated | none detected | none detected |
| MHI3711 | 73 | 1030 | present | unencapsulated | none detected | none detected |
| MHI3712 | 73 | 1030 | present | unencapsulated | none detected | none detected |
| MHI3713 | 73 | 1030 | present | unencapsulated | none detected | none detected |
| MHI3714 | 73 | 1030 | present | unencapsulated | none detected | none detected |
| MHI3715 | 73 | 1030 | present | unencapsulated | none detected | none detected |
| MHI3716 | 75 | 544 | present | unencapsulated | none detected | none detected |
| MHI3717 | 75 | 544 | present | unencapsulated | none detected | none detected |
| MHI3718 | 75 | 544 | present | unencapsulated | none detected | none detected |
| MHI3719 | 75 | 544 | present | unencapsulated | none detected | none detected |
| MHI3720 | 75 | 544 | present | unencapsulated | none detected | none detected |
| MHI3721 | 75 | 544 | present | unencapsulated | none detected | none detected |
| MHI3722 | 75 | 544 | present | unencapsulated | none detected | none detected |
| MHI3723 | 75 | 544 | present | unencapsulated | none detected | none detected |
| MHI3724 | 75 | 544 | present | unencapsulated | none detected | none detected |
| MHI3725 | 75 | 544 | present | unencapsulated | none detected | none detected |
| MHI3726 | 75 | 544 | present | unencapsulated | none detected | none detected |
| MHI3727 | 75 | 544 | present | unencapsulated | none detected | none detected |
| MHI3728 | 75 | 544 | present | unencapsulated | none detected | none detected |
| MHI3729 | 75 | 544 | present | unencapsulated | none detected | none detected |
| MHI3730 | 75 | 544 | present | unencapsulated | none detected | none detected |
| MHI3731 | 75 | 544 | present | unencapsulated | none detected | none detected |
| MHI3732 | 75 | 544 | present | unencapsulated | none detected | none detected |
| MHI3733 | 75 | 544 | present | unencapsulated | none detected | none detected |
| MHI3734 | 75 | 544 | present | unencapsulated | none detected | none detected |
| MHI3735 | 75 | 544 | present | unencapsulated | none detected | none detected |
| MHI3736 | 75 | 544 | present | unencapsulated | none detected | none detected |
| MHI3737 | 75 | 544 | present | unencapsulated | none detected | none detected |
| MHI3738 | 75 | 544 | present | unencapsulated | none detected | none detected |
| MHI3739 | 75 | 544 | present | unencapsulated | none detected | none detected |
| MHI3740 | 75 | 544 | present | unencapsulated | none detected | none detected |
| MHI3741 | 75 | 544 | present | unencapsulated | none detected | none detected |
| MHI3742 | 75 | 544 | present | unencapsulated | none detected | none detected |
| MHI3743 | 75 | 544 | present | unencapsulated | none detected | none detected |
| MHI3744 | 75 | 544 | present | unencapsulated | none detected | none detected |
| MHI3745 | 75 | 544 | present | unencapsulated | none detected | none detected |
| MHI3746 | 75 | 544 | present | unencapsulated | none detected | none detected |
| MHI3747 | 75 | 544 | present | unencapsulated | none detected | none detected |

**Table S2. MLST alleles of the 39 Child-MLST matched reference genomes.**

| Child | Isolate | Year | ST | *adk* | *atpG* | *frdB* | *fucK* | *mdh* | *pgi* | *recA* |
|---|---|---|---|---|---|---|---|---|---|---|
| 1 | MHI3301 | 2018 | 266 | 3 | 18 | 53 | 15 | 86 | 14 | 23 |
| 2 | MHI3331 | 2018 | 1040 | 63 | 8 | 58 | 14 | 92 | 58 | 29 |
| 2 | MHI3332 | 2018 | 544 | 5 | 33 | 7 | 15 | 22 | 1 | 29 |
| 2 | MHI3338 | 2018 | 1191 | 39 | 8 | 35 | 1 | 63 | 160 | 19 |
| 2 | MHI3341 | 2018 | 652 | 6 | 43 | 7 | 7 | 77 | 71 | 43 |
| 7 | MHI3578 | 2018 | 159 | 33 | 8 | 16 | 16 | 17 | 2 | 29 |
| 8 | MHI3941 | 2018 | 431 | 5 | 33 | 7 | 15 | 142 | 117 | 29 |
| 8 | MHI3963 | 2018 | 2955 | 5 | 33 | 7 | 15 | 142 | 117 | 146 |
| 10 | MHI3976 | 2018 | 57 | 14 | 7 | 13 | 7 | 17 | 13 | 17 |
| 13 | MHI4015 | 2018 | 2156 | 4 | 110 | 71 | 4 | 84 | 57 | 35 |
| 15 | MHI4050 | 2018 | 2956 | 166 | 118 | 44 | 0 | 248 | 145 | 99 |
| 15 | MHI4051 | 2018 | 2957 | 4 | 15 | 106 | 87 | 163 | 186 | 41 |
| 15 | MHI4060 | 2018 | 1238 | 35 | 33 | 7 | 15 | 154 | 71 | 29 |
| 16 | MHI3361 | 2018 | 395 | 14 | 51 | 16 | 48 | 15 | 2 | 31 |
| 20 | MHI4090 | 1018 | 946 | 151 | 77 | 53 | 8 | 210 | 2 | 34 |
| 27 | MHI3511 | 2017 | 1034 | 125 | 1 | 1 | 1 | 1 | 2 | 5 |
| 27 | MHI3512 | 2017 | 683 | 93 | 41 | 41 | 8 | 69 | 60 | 98 |
| 27 | MHI3391 | 2018 | 2954 | 1 | 1 | 1 | 1 | 87 | 1 | 5 |
| 27 | MHI3401 | 2018 | 494 | 1 | 1 | 1 | 1 | 148 | 1 | 5 |
| 29 | MHI3542 | 2017 | 199 | 6 | 20 | 41 | 40 | 88 | 63 | 43 |
| 29 | MHI3543 | 2017 | 598 | 22 | 4 | 11 | 11 | 22 | 19 | 15 |
| 29 | MHI3546 | 2017 | 536 | 30 | 1 | 22 | 36 | 51 | 31 | 29 |
| 29 | MHI3421 | 2018 | 266 | 3 | 18 | 53 | 15 | 86 | 14 | 23 |
| 29 | MHI3428 | 2018 | 1030 | 23 | 3 | 5 | 1 | 166 | 66 | 21 |
| 36 | MHI4125 | 2018 | 925 | 37 | 35 | 57 | 8 | 7 | 183 | 13 |
| 39 | MHI4160 | 2018 | 159 | 33 | 8 | 16 | 16 | 17 | 2 | 29 |
| 43 | MHI3451 | 2018 | 836 | 1 | 11 | 18 | 18 | 62 | 1 | 5 |
| 46 | MHI3481 | 2018 | 266 | 3 | 18 | 53 | 15 | 86 | 14 | 23 |
| 52 | MHI3620 | 2018 | 105 | 3 | 9 | 8 | 2 | 14 | 8 | 4 |
| 56 | MHI4194 | 2018 | 334 | 71 | 17 | 58 | 18 | 35 | 58 | 56 |
| 57 | MHI4227 | 2018 | 161 | 41 | 8 | 29 | 9 | 68 | 57 | 8 |
| 59 | MHI4262 | 2018 | 1409 | 5 | 33 | 7 | 15 | 58 | 77 | 29 |
| 59 | MHI4264 | 2018 | 280 | 3 | 9 | 8 | 2 | 86 | 8 | 4 |
| 61 | MHI3652 | 2018 | 568 | 3 | 9 | 8 | 4 | 26 | 8 | 4 |
| 61 | MHI3655 | 2018 | 1012 | 35 | 25 | 16 | 53 | 7 | 40 | 10 |
| 64 | MHI4297 | 2018 | 2958 | 5 | 33 | 7 | 15 | 1 | 4 | 29 |
| 68 | MHI4332 | 2018 | 264 | 5 | 8 | 58 | 14 | 91 | 13 | 29 |
| 73 | MHI3685 | 2018 | 1030 | 23 | 3 | 5 | 1 | 166 | 66 | 21 |
| 75 | MHI3716 | 2018 | 544 | 5 | 33 | 7 | 15 | 22 | 1 | 29 |

**Table S3. The 99 publically available *H. influenzae* genomes.**

| Assembly Name | GeneBank | RefSeq | Bases | Genes | Date | Submitter |
|---|---|---|---|---|---|---|
| 33763_D01 | GCA_900478735.1 | GCF_900478735.1 | 1,865,137 | 1,817 | 06/18/18 | Wellcome Trust Sanger insititute |
| 33962_B02 | GCA_900635795.1 | GCF_900635795.1 | 1,860,106 | 1,828 | 12/20/18 | Wellcome Trust Sanger insititute |
| 33962_G01 | GCA_900478325.1 | GCF_900478325.1 | 1,915,356 | 1,938 | 06/18/18 | Wellcome Trust Sanger insititute |
| 34211_C02 | GCA_900635805.1 | GCF_900635805.1 | 1,879,945 | 1,831 | 12/20/18 | Wellcome Trust Sanger insititute |
| 34211_D02 | GCA_900478275.1 | GCF_900478275.1 | 1,890,469 | 1,861 | 06/18/18 | Wellcome Trust Sanger insititute |
| 35860_G01 | GCA_901472485.1 | GCF_901472485.1 | 1,876,886 | 1,840 | 05/11/19 | Wellcome Trust Sanger insititute |
| 44310_E01 | GCA_900475535.1 | GCF_900475535.1 | 2,044,007 | 2,092 | 06/17/18 | Wellcome Trust Sanger insititute |
| 45214_B02 | GCA_900475755.1 | GCF_900475755.1 | 1,834,484 | 1,772 | 06/17/18 | Wellcome Trust Sanger insititute |
| 47555_F02 | GCA_900475995.1 | GCF_900475995.1 | 1,850,791 | 1,808 | 06/17/18 | Wellcome Trust Sanger insititute |
| 56433_A01 | GCA_900638105.1 | GCF_900638105.1 | 1,948,880 | 1,944 | 12/20/18 | Wellcome Trust Sanger insititute |
| ASM1218v1 | GCA_000012185.1 | GCF_000012185.1 | 1,914,490 | 1,899 | 08/30/07 | Columbus Children's Research Institute |
| ASM1339440v1 | GCA_013394405.1 | GCF_013394405.1 | 1,957,393 | 2,003 | 06/09/20 | Tokyo University of Pharmacy and Life Sciences |
| ASM1470121v1 | GCA_014701215.1 | GCF_014701215.1 | 1,809,645 | 1,773 | 09/15/20 | National Institute of Infectious Diseases |
| ASM1493149v1 | GCA_014931495.1 | GCF_014931495.1 | 1,894,676 | 1,878 | 10/26/20 | Monash University |
| ASM1646v1 | GCA_000016465.1 | GCF_000016465.1 | 1,813,033 | 1,695 | 06/07/07 | Allegheny-Singer Research Institute |
| ASM16552v1 | GCA_000165525.1 | GCF_000165525.1 | 1,932,306 | 1,892 | 10/22/10 | Seattle Biomedical Research Institute |
| ASM16557v1 | GCA_000165575.1 | GCF_000165575.1 | 1,819,370 | 1,744 | 10/25/10 | Seattle Biomedical Research Institute |
| ASM1686128v1 | GCA_016861285.1 | GCF_016861285.1 | 1,852,308 | 1,842 | 01/28/21 | Tokyo Metropolitan Institute of Public Health |
| ASM185672v1 | GCA_001856725.1 | GCF_001856725.1 | 1,829,217 | 1,781 | 10/31/16 | Public Health Agency of Canada |
| ASM1970336v1 | GCA_019703365.1 | GCF_019703365.1 | 1,979,718 | 2,002 | 05/22/21 | Chiba University |
| ASM1970338v1 | GCA_019703385.1 | GCF_019703385.1 | 1,979,786 | 2,002 | 05/22/21 | Chiba University |
| ASM1970340v1 | GCA_019703405.1 | GCF_019703405.1 | 1,853,393 | 1,825 | 05/22/21 | Chiba University |
| ASM1970342v1 | GCA_019703425.1 | GCF_019703425.1 | 1,832,515 | 1,806 | 05/22/21 | Chiba University |
| ASM1970345v1 | GCA_019703455.1 | GCF_019703455.1 | 1,805,925 | 1,769 | 05/22/21 | Chiba University |
| ASM1970349v1 | GCA_019703495.1 | GCF_019703495.1 | 1,819,364 | 1,783 | 05/22/21 | Chiba University |
| ASM1970352v1 | GCA_019703525.1 | GCF_019703525.1 | 1,886,480 | 1,883 | 05/22/21 | Chiba University |
| ASM1970354v1 | GCA_019703545.1 | GCF_019703545.1 | 1,902,536 | 1,909 | 05/22/21 | Chiba University |
| ASM1970357v1 | GCA_019703575.1 | GCF_019703575.1 | 1,816,076 | 1,760 | 05/22/21 | Chiba University |
| ASM1970359v1 | GCA_019703595.1 | GCF_019703595.1 | 1,914,722 | 1,899 | 05/22/21 | Chiba University |
| ASM1970361v1 | GCA_019703615.1 | GCF_019703615.1 | 1,810,977 | 1,753 | 05/22/21 | Chiba University |
| ASM1970363v1 | GCA_019703635.1 | GCF_019703635.1 | 1,818,266 | 1,780 | 05/22/21 | Chiba University |
| ASM1970365v1 | GCA_019703655.1 | GCF_019703655.1 | 1,822,115 | 1,779 | 05/22/21 | Chiba University |
| ASM1970367v1 | GCA_019703675.1 | GCF_019703675.1 | 1,884,503 | 1,879 | 05/22/21 | Chiba University |
| ASM1970369v1 | GCA_019703695.1 | GCF_019703695.1 | 1,911,890 | 1,929 | 05/22/21 | Chiba University |
| ASM1970371v1 | GCA_019703715.1 | GCF_019703715.1 | 1,791,343 | 1,749 | 05/22/21 | Chiba University |
| ASM1970373v1 | GCA_019703735.1 | GCF_019703735.1 | 1,882,642 | 1,866 | 05/22/21 | Chiba University |
| ASM1970375v1 | GCA_019703755.1 | GCF_019703755.1 | 1,953,394 | 1,949 | 05/22/21 | Chiba University |
| ASM1970377v1 | GCA_019703775.1 | GCF_019703775.1 | 1,790,924 | 1,768 | 05/22/21 | Chiba University |
| ASM1970379v1 | GCA_019703795.1 | GCF_019703795.1 | 1,936,686 | 1,941 | 05/22/21 | Chiba University |
| ASM19787v1 | GCA_000197875.1 | GCF_000197875.1 | 1,985,832 | 1,810 | 01/11/11 | Wellcome Trust Sanger Institute |
| ASM1993070v1 | GCA_019930705.1 | GCF_019930705.1 | 1,902,597 | 1,899 | 09/12/21 | US Food and Drug Administration |
| ASM20047v1 | GCA_000200475.1 | GCF_000200475.1 | 2,007,018 | 1,820 | 01/11/11 | Wellcome Trust Sanger insititute |
| ASM207347v2 | GCA_002073475.2 | GCF_002073475.2 | 1,830,681 | 1,823 | 02/23/18 | University of Maryland |
| ASM2073602v1 | GCA_020736025.1 | GCF_020736025.1 | 1,834,285 | 1,789 | 11/03/21 | US Food and Drug Administration |
| ASM2073604v1 | GCA_020736045.1 | GCF_020736045.1 | 1,890,662 | 1,896 | 11/03/21 | US Food and Drug Administration |
| ASM21087v1 | GCA_000210875.1 | GCF_000210875.1 | 1,981,535 | 1,914 | 07/23/10 | Wellcome Trust Sanger Institute |
| ASM2853503v1 | GCA_028535035.1 | Not assigned | 1,812,138 | 1,775 | 02/07/23 | University of Maryland |
| ASM296657v1 | GCA_002966575.1 | GCF_002966575.1 | 1,857,175 | 1,850 | 03/02/18 | University of Maryland |
| ASM296659v1 | GCA_002966595.1 | GCF_002966595.1 | 1,857,048 | 1,851 | 03/02/18 | University of Maryland |
| ASM296661v1 | GCA_002966615.1 | GCF_002966615.1 | 1,886,450 | 1,922 | 03/02/18 | University of Maryland |
| ASM296663v1 | GCA_002966635.1 | GCF_002966635.1 | 1,886,411 | 1,932 | 03/02/18 | University of Maryland |
| ASM296665v1 | GCA_002966655.1 | GCF_002966655.1 | 1,823,096 | 1,818 | 03/02/18 | University of Maryland |
| ASM296667v1 | GCA_002966675.1 | GCF_002966675.1 | 1,823,172 | 1,818 | 03/02/18 | University of Maryland |
| ASM296669v1 | GCA_002966695.1 | GCF_002966695.1 | 1,901,558 | 1,908 | 03/02/18 | University of Maryland |
| ASM296671v1 | GCA_002966715.1 | GCF_002966715.1 | 1,901,533 | 1,911 | 03/02/18 | University of Maryland |
| ASM296673v1 | GCA_002966735.1 | GCF_002966735.1 | 1,813,865 | 1,799 | 03/02/18 | University of Maryland |
| ASM298821v2 | GCA_002988215.2 | GCF_002988215.2 | 1,812,418 | 1,777 | 02/06/23 | University of Maryland |
| ASM298888v2 | GCA_002988885.2 | GCF_002988885.2 | 1,812,134 | 1,772 | 02/06/23 | University of Maryland |
| ASM318438v1 | GCA_003184385.1 | GCF_003184385.1 | 2,047,595 | 2,079 | 06/04/18 | Griffith University |
| ASM318440v1 | GCA_003184405.1 | GCF_003184405.1 | 2,025,527 | 2,115 | 06/04/18 | Griffith University |
| ASM335142v1 | GCF_003351425.1 | GCF_003351425.1 | 1,817,261 | 1,761 | 08/01/18 | Centers for Disease Control and Prevention |
| ASM335144v1 | GCF_003351445.1 | GCF_003351445.1 | 1,860,196 | 1,850 | 08/01/18 | Centers for Disease Control and Prevention |
| ASM335146v1 | GCF_003351465.1 | GCF_003351465.1 | 1,816,295 | 1,783 | 08/01/18 | Centers for Disease Control and Prevention |
| ASM335158v1 | GCF_003351585.1 | GCF_003351585.1 | 1,848,871 | 1,846 | 08/01/18 | Centers for Disease Control and Prevention |
| ASM335160v1 | GCF_003351605.1 | GCF_003351605.1 | 1,804,746 | 1,772 | 08/01/18 | Centers for Disease Control and Prevention |
| ASM335234v1 | GCF_003352345.1 | GCF_003352345.1 | 1,914,782 | 1,895 | 08/01/18 | Centers for Disease Control and Prevention |
| ASM335236v1 | GCF_003352365.1 | GCF_003352365.1 | 1,887,933 | 1,862 | 08/01/18 | Centers for Disease Control and Prevention |

| | | | | | | |
|---|---|---|---|---|---|---|
| ASM335240v1 | GCF_003352405.1 | GCF_003352405.1 | 1,919,901 | 1,922 | 08/01/18 | Centers for Disease Control and Prevention |
| ASM342544v1 | GCF_003425445.1 | GCF_003425445.1 | 1,838,740 | 1,820 | 08/23/18 | Institute of Agrobiotechnology |
| ASM342546v1 | GCF_003425465.1 | GCF_003425465.1 | 1,833,710 | 1,817 | 08/23/18 | Institute of Agrobiotechnology |
| ASM342548v1 | GCF_003425485.1 | GCF_003425485.1 | 1,811,303 | 1,767 | 08/23/18 | Institute of Agrobiotechnology |
| ASM342550v1 | GCF_003425505.1 | GCF_003425505.1 | 2,013,003 | 2,061 | 08/23/18 | Institute of Agrobiotechnology |
| ASM342552v1 | GCF_003425525.1 | GCF_003425525.1 | 1,897,311 | 1,905 | 08/23/18 | Institute of Agrobiotechnology |
| ASM342556v1 | GCF_003425565.1 | GCF_003425565.1 | 1,858,634 | 1,839 | 08/23/18 | Institute of Agrobiotechnology |
| ASM342558v1 | GCF_003425585.1 | GCF_003425585.1 | 1,840,498 | 1,817 | 08/23/18 | Institute of Agrobiotechnology |
| ASM342560v1 | GCF_003425605.1 | GCF_003425605.1 | 1,858,630 | 1,839 | 08/23/18 | Institute of Agrobiotechnology |
| ASM342562v1 | GCF_003425625.1 | GCF_003425625.1 | 1,833,305 | 1,799 | 08/23/18 | Institute of Agrobiotechnology |
| ASM342564v1 | GCF_003425645.1 | GCF_003425645.1 | 1,908,143 | 1,941 | 08/23/18 | Institute of Agrobiotechnology |
| ASM342571v1 | GCF_003425715.1 | GCF_003425715.1 | 1,848,210 | 1,832 | 08/23/18 | Institute of Agrobiotechnology |
| ASM342576v1 | GCF_003425765.1 | GCF_003425765.1 | 1,833,864 | 1,819 | 08/23/18 | Institute of Agrobiotechnology |
| ASM342581v1 | GCF_003425815.1 | GCF_003425815.1 | 1,840,062 | 1,824 | 08/23/18 | Institute of Agrobiotechnology |
| ASM342593v1 | GCF_003425935.1 | GCF_003425935.1 | 1,849,483 | 1,836 | 08/23/18 | Institute of Agrobiotechnology |
| ASM342595v1 | GCF_003425955.1 | GCF_003425955.1 | 1,904,311 | 1,903 | 08/23/18 | Institute of Agrobiotechnology |
| ASM46525v1 | GCF_000465255.1 | GCF_000465255.1 | 1,856,176 | 1,812 | 09/09/13 | Lund University |
| ASM69836v1 | GCF_000698365.1 | GCF_000698365.1 | 1,811,802 | 1,773 | 06/05/14 | Center for Genomic Science |
| ASM76707v1 | GCF_000767075.1 | GCF_000767075.1 | 1,850,897 | 1,842 | 10/16/14 | Drexel University |
| ASM858674v1 | GCF_008586745.1 | GCF_008586745.1 | 1,985,844 | 2,024 | 09/19/19 | Griffith University |
| ASM858676v1 | GCF_008586765.1 | GCF_008586765.1 | 2,000,194 | 2,038 | 09/19/19 | Griffith University |
| ASM858678v1 | GCF_008586785.1 | GCF_008586785.1 | 1,877,864 | 1,896 | 09/19/19 | Griffith University |
| ASM858680v1 | GCF_008586805.1 | GCF_008586805.1 | 1,987,687 | 2,030 | 09/19/19 | Griffith University |
| ASM858682v1 | GCF_008586825.1 | GCF_008586825.1 | 1,984,979 | 2,024 | 09/19/19 | Griffith University |
| ASM883152v1 | GCF_008831525.1 | GCF_008831525.1 | 1,887,343 | 1,879 | 10/07/19 | Drexel University |
| ASM93157v1 | GCF_000931575.1 | GCF_000931575.1 | 1,846,259 | 1,834 | 02/23/15 | Griffith University |
| ASM93160v1 | GCF_000931605.1 | GCF_000931605.1 | 1,846,503 | 1,813 | 02/23/15 | Griffith University |
| ASM93162v1 | GCF_000931625.1 | GCF_000931625.1 | 1,887,620 | 1,899 | 02/23/15 | Griffith University |
| ASM96833v1 | GCF_000968335.1 | GCF_000968335.1 | 1,969,659 | 2,009 | 04/02/15 | Research Institute at Nationwide Children's Hospital |
| KRLund_NTHi_Asse | GCF_919946725.1 | GCF_919946725.1 | 1,805,497 | 1,785 | 12/03/21 | Lund University |
| KRLund_NTHi_Asse | GCF_919949215.1 | GCF_919949215.1 | 1,917,048 | 1,928 | 12/03/21 | Lund University |
| NCTC8143 | GCF_001457655.1 | GCF_001457655.1 | 1,890,645 | 1,890 | 03/22/15 | Wellcome Trust Sanger insititute |

**Table S4. Polymorphisms identified in 805 *H. influenzae* asymptomatic carrier isolates.**

| Gene | Protein | Function | Polymorphism | Substitution | Position in codon | Child-MLST | Isolate |
|---|---|---|---|---|---|---|---|
| *adk* | Adenylate kinase | transcription/translation | C640G | G214R | 640 | 1-266 | MHI4370, |
| *adk* | Adenylate kinase | transcription/translation | C640G | G214R | 640 | 52-105 | MHI3632, MHI3633, |
| *adk* | Adenylate kinase | transcription/translation | C641A | G214V | 641 | 1-266 | MHI4370, |
| *adk* | Adenylate kinase | transcription/translation | C641A | G214V | 641 | 29-199 | MHI3563, |
| *adk* | Adenylate kinase | transcription/translation | C641A | G214V | 641 | 52-105 | MHI3632, MHI3633, |
| *adk* | Adenylate kinase | transcription/translation | T643C | *215Q | 643 | 15-1238 | MHI4074, |
| *adk* | Adenylate kinase | transcription/translation | A643G | *215Q | 643 | 68-264 | MHI4345, |
| *adk* | Adenylate kinase | transcription/translation | T644A | *215L | 644 | 1-266 | MHI4370, |
| *adk* | Adenylate kinase | transcription/translation | T644A | *215L | 644 | 7-159 | MHI4382, |
| *adk* | Adenylate kinase | transcription/translation | A644T | *215L | 644 | 20-946 | MHI4092, |
| *adk* | Adenylate kinase | transcription/translation | T644A | *215L | 644 | 15-2957 | MHI4087, |
| *adk* | Adenylate kinase | transcription/translation | T644A | *215L | 644 | 29-199 | MHI3547, MHI3558, MHI3561, MHI3563, |
| *adk* | Adenylate kinase | transcription/translation | T644A | *215L | 644 | 52-105 | MHI3632, MHI3633, MHI3639, |
| *adk* | Adenylate kinase | transcription/translation | A644T | *215L | 644 | 61-1012 | MHI3667, |
| *adk* | Adenylate kinase | transcription/translation | T644A | *215L | 644 | 8-431 | MHI3953, |
| *adk* | Adenylate kinase | transcription/translation | A644T | *215L | 644 | 8-2955 | MHI3965, MHI3978, |
| *argG* | Aerobic respiration control sensor protein ArcB | regulator | T1321A | L441I | 1321 | 39-159 | MHI4172, |
| *argG* | Aerobic respiration control sensor protein ArcB | regulator | T1321A | L441I | 1321 | 15-2957 | MHI4075, MHI4081, |
| *aroA* | 3-phosphoshikimate 1-carboxyvinyltransferase | transcription/translation | T1294C | N432D | 1294 | 29-266 | MHI3431, |
| *aroA* | 3-phosphoshikimate 1-carboxyvinyltransferase | transcription/translation | T1295C | N432S | 1295 | 1-266 | MHI3575, |
| *aroA* | 3-phosphoshikimate 1-carboxyvinyltransferase | transcription/translation | A1297G | *433Q | 1297 | 1-266 | MHI3575, |
| *aroA* | 3-phosphoshikimate 1-carboxyvinyltransferase | transcription/translation | A1297G | *433Q | 1297 | 29-199 | MHI3556, |
| *aroA* | 3-phosphoshikimate 1-carboxyvinyltransferase | transcription/translation | A1298C | *433S | 1298 | 7-159 | MHI4382, |
| *artP* | Arginine transport ATP-binding protein | transporter | T109A | S37C | 109 | 10-57 | MHI4003, |
| *artP* | Arginine transport ATP-binding protein | transporter | C110G | S37T | 110 | 10-57 | MHI4003, |
| *artP* | Arginine transport ATP-binding protein | transporter | T139A | T47S | 139 | 10-57 | MHI4003, |
| *artP* | Arginine transport ATP-binding protein | transporter | G140C | T47R | 140 | 10-57 | MHI4003, |
| *aspA* | Aspartate ammonia-lyase | amino acid metabolism | A1346C | E449A | 1346 | 61-1012 | MHI3658, |
| *atsB* | Serine-type anaerobic sulfatase-maturating enzyme | transcription/translation | T218A | Y73F | 218 | 43-836 | MHI3454, MHI3467, MHI3478, |
| *atsB* | Serine-type anaerobic sulfatase-maturating enzyme | transcription/translation | T225A | R75S | 225 | 43-836 | MHI3454, MHI3467, MHI3478, |
| *atsB* | Serine-type anaerobic sulfatase-maturating enzyme | transcription/translation | T240G | *80Y | 240 | 43-836 | MHI3454, |
| *bacA* | Prephenate decarboxylase | antibiotic biosynthesis | C36G | N12K | 36 | 56-344 | MHI4213, |
| *bacA* | Prephenate decarboxylase | antibiotic biosynthesis | T40A | I14F | 40 | 10-57 | MHI4002, |
| *bacA* | Prephenate decarboxylase | antibiotic biosynthesis | T57A | F19L | 57 | 56-344 | MHI4213, |
| *bacA* | Prephenate decarboxylase | antibiotic biosynthesis | A40T | I14F | 40 | 27-683 | MHI3518, |
| *bacA* | Prephenate decarboxylase | antibiotic biosynthesis | T56A | F19Y | 56 | 27-683 | MHI3518, |
| *bacA* | Prephenate decarboxylase | antibiotic biosynthesis | T58G | W20G | 58 | 27-683 | MHI3518, MHI4015, |
| *bacA* | Prephenate decarboxylase | antibiotic biosynthesis | G60T | W20C | 60 | 27-683 | MHI3518, |
| *bacA* | Prephenate decarboxylase | antibiotic biosynthesis | G61A | V21I | 61 | 27-683 | MHI3518, |
| *bacA* | Prephenate decarboxylase | antibiotic biosynthesis | C518G | T173S | 518 | 13-2156 | MHI4017, MHI4022, MHI4031, MHI4033, MHI4040, |
| *bacA* | Prephenate decarboxylase | antibiotic biosynthesis | G556A | V186I | 556 | 13-2156 | MHI4017, MHI4022, MHI4028, MHI4031, MHI4032, MHI4033, MHI4040, MHI4046, |
| *bamA* | Outer membrane protein assembly factor BamA | transporter | C1982G | G661A | 1982 | 1-266 | MHI3320, |
| *bamA* | Outer membrane protein assembly factor BamA | transporter | T2322A | K774N | 2322 | 68-264 | MHI4358, |
| *bamA* | Outer membrane protein assembly factor BamA | transporter | A2323T | Y775N | 2323 | 68-264 | MHI4358, |
| *bcp* | ATP-dependent dethiobiotin synthetase BioD 1 | transcription/translation | T469C | Y157H | 469 | 75-544 | MHI3721, |
| *bioD1* | ATP-dependent dethiobiotin synthetase BioD 1 | transcription/translation | A715G | *239Q | 715 | 16-396 | MHI4390, |
| *bioD1* | ATP-dependent dethiobiotin synthetase BioD 1 | transcription/translation | T724C | K242E | 724 | 20-946 | MHI4122, |
| *bioD1* | ATP-dependent dethiobiotin synthetase BioD 1 | transcription/translation | A727C | *243E | 727 | 20-946 | MHI4093, MHI4100, MHI4115, |
| *bioD1* | ATP-dependent dethiobiotin synthetase BioD 1 | transcription/translation | T727G | *243E | 727 | 75-544 | MHI3726, |
| *bioD1* | ATP-dependent dethiobiotin synthetase BioD 1 | transcription/translation | T706G | M236L | 706 | 15-2957 | MHI4088, MHI4089, |
| *bioD1* | ATP-dependent dethiobiotin synthetase BioD 1 | transcription/translation | T706G | M236L | 706 | 8-431 | MHI3954, |
| *bioD1* | ATP-dependent dethiobiotin synthetase BioD 1 | transcription/translation | A707T | M236K | 707 | 15-2957 | MHI4088, MHI4089, |
| *bioD1* | ATP-dependent dethiobiotin synthetase BioD 1 | transcription/translation | A707T | M236K | 707 | 8-431 | MHI3954, |
| *bioD1* | ATP-dependent dethiobiotin synthetase BioD 1 | transcription/translation | C708T | M236I | 708 | 15-2957 | MHI4088, MHI4089, |
| *bioD1* | ATP-dependent dethiobiotin synthetase BioD 1 | transcription/translation | C708T | M236I | 708 | 8-431 | MHI3954, |
| *bioD1* | ATP-dependent dethiobiotin synthetase BioD 1 | transcription/translation | T710G | Q237P | 710 | 15-2957 | MHI4082, MHI4088, MHI4089, |
| *bioD1* | ATP-dependent dethiobiotin synthetase BioD 1 | transcription/translation | T710G | Q237P | 710 | 8-431 | MHI3954, |
| *bioD1* | ATP-dependent dethiobiotin synthetase BioD 1 | transcription/translation | A715G | *239Q | 715 | 15-2957 | MHI4053, MHI4054, MHI4055, MHI4057, MHI4063, MHI4065, MHI4066, MHI4067, MHI4069, MHI4070, MHI4071, MHI4072, MHI4073, MHI4075, MHI4076, MHI4078, MHI4081, MHI4084, MHI4086, MHI4087, |
| *bioD1* | ATP-dependent dethiobiotin synthetase BioD 1 | transcription/translation | T715C | *239Q | 715 | 2-1040 | MHI3343, |
| *bioD1* | ATP-dependent dethiobiotin synthetase BioD 1 | transcription/translation | A715G | *239Q | 715 | 68-264 | MHI3532, MHI4333, MHI4342, MHI4353, |
| *bioD1* | ATP-dependent dethiobiotin synthetase BioD 1 | transcription/translation | A715G | *239Q | 715 | 8-431 | MHI3954, MHI3974, |
| *bioD1* | ATP-dependent dethiobiotin synthetase BioD 1 | transcription/translation | T727G | *243E | 727 | 52-105 | MHI3650, |
| *bolA* | DNA-binding transcriptional regulator | regulator | A251T | E84V | 251 | 73-1030 | MHI3714, |
| *bolA* | DNA-binding transcriptional regulator | regulator | T251A | E84V | 251 | 75-544 | MHI3730, MHI3737, |
| *bolA* | DNA-binding transcriptional regulator | regulator | A251T | E84V | 251 | 15-2957 | MHI4076, |
| *bolA* | DNA-binding transcriptional regulator | regulator | A251T | E84V | 251 | 27-683 | MHI3532, |
| *bolA* | DNA-binding transcriptional regulator | regulator | A269T | E90V | 269 | 27-683 | MHI3515, |
| *bolA* | DNA-binding transcriptional regulator | regulator | C272T | T91I | 272 | 27-683 | MHI3515, |
| *bspRIM* | Cytosine-specific methyltransferase | transcription/translation | T660A | K220N | 660 | 56-344 | MHI4198, |
| *bspRIM* | Cytosine-specific methyltransferase | transcription/translation | C661G | A221P | 661 | 56-344 | MHI4198, |
| *bspRIM* | Cytosine-specific methyltransferase | transcription/translation | G662C | A221G | 662 | 56-344 | MHI3419, MHI4198, |
| *bspRIM* | Cytosine-specific methyltransferase | transcription/translation | A664C | L222V | 664 | 2-1191 | MHI4378, |
| *bspRIM* | Cytosine-specific methyltransferase | transcription/translation | A664C | L222V | 664 | 56-344 | MHI4198, |
| *bspRIM* | Cytosine-specific methyltransferase | transcription/translation | A665G | L222S | 665 | 2-1191 | MHI4378, |
| *bspRIM* | Cytosine-specific methyltransferase | transcription/translation | A665G | L222S | 665 | 29-199 | MHI3556, |
| *bspRIM* | Cytosine-specific methyltransferase | transcription/translation | A665G | L222S | 665 | 56-344 | MHI4198, |
| *bspRIM* | Cytosine-specific methyltransferase | transcription/translation | T665C | L222S | 665 | 61-1012 | MHI3656, MHI3661, MHI3669, |
| *bspRIM* | Cytosine-specific methyltransferase | transcription/translation | A687T | N229K | 687 | 29-199 | MHI3561, |
| *bspRIM* | Cytosine-specific methyltransferase | transcription/translation | A687T | N229K | 687 | 56-344 | MHI4198, |
| *bspRIM* | Cytosine-specific methyltransferase | transcription/translation | T687A | N229K | 687 | 61-1012 | MHI3656, MHI3658, MHI3661, MHI3668, MHI3669, |
| *bspRIM* | Cytosine-specific methyltransferase | transcription/translation | T690G | I230M | 690 | 61-1012 | MHI3656, MHI3658, MHI3661, MHI3668, MHI3669, |
| *bspRIM* | Cytosine-specific methyltransferase | transcription/translation | G694T | E232* | 694 | 61-1012 | MHI3658, MHI3661, MHI3668, MHI3669, |
| *bspRIM* | Cytosine-specific methyltransferase | transcription/translation | A695C | E232A | 695 | 61-1012 | MHI3658, MHI3661, MHI3668, MHI3669, |
| *bspRIM* | Cytosine-specific methyltransferase | transcription/translation | A700C | L234V | 700 | 27-683 | MHI3535, |
| *bspRIM* | Cytosine-specific methyltransferase | transcription/translation | A701G | L234S | 701 | 15-2957 | MHI4075, |
| *bspRIM* | Cytosine-specific methyltransferase | transcription/translation | A701G | L234S | 701 | 27-683 | MHI3525, MHI3535, |
| *bspRIM* | Cytosine-specific methyltransferase | transcription/translation | A723T | N241K | 723 | 15-2957 | MHI4075, MHI4076, |
| *bspRIM* | Cytosine-specific methyltransferase | transcription/translation | A723T | N241K | 723 | 27-683 | MHI3525, MHI3541, |
| *bspRIM* | Cytosine-specific methyltransferase | transcription/translation | A726C | I242M | 726 | 15-2957 | MHI4075, MHI4076, |
| *bspRIM* | Cytosine-specific methyltransferase | transcription/translation | A726C | I242M | 726 | 27-683 | MHI3525, MHI3541, |
| *can* | Carbonic anhydrase | environmental stress response | T685G | T229P | 685 | 73-1030 | MHI3705, |
| *can* | Carbonic anhydrase | environmental stress response | G685T | T229P | 685 | 29-1030 | MHI3432, MHI3448, |
| *can* | Carbonic anhydrase | environmental stress response | T685G | T229P | 685 | 61-1012 | MHI3658, |
| *can* | Carbonic anhydrase | environmental stress response | G686A | T229I | 686 | 52-105 | MHI3629, |
| *can* | Carbonic anhydrase | environmental stress response | G686A | T229I | 686 | 61-1012 | MHI3658, |
| *can* | Carbonic anhydrase | environmental stress response | A688T | *230K | 688 | 10-57 | MHI4003, |
| *can* | Carbonic anhydrase | environmental stress response | A688T | *230K | 688 | 46-266 | MHI3489, |
| *can* | Carbonic anhydrase | environmental stress response | A688G | *230Q | 688 | 73-1030 | MHI3698, MHI3705, |
| *can* | Carbonic anhydrase | environmental stress response | A688G | *230Q | 688 | 29-1030 | MHI3432, MHI3448, |
| *can* | Carbonic anhydrase | environmental stress response | A688T | *230K | 688 | 52-105 | MHI3628, MHI3629, |
| *can* | Carbonic anhydrase | environmental stress response | A688T | *230K | 688 | 61-1012 | MHI3658, |
| *can* | Carbonic anhydrase | environmental stress response | T690A | *230Y | 690 | 46-266 | MHI3489, MHI3491, |
| *can* | Carbonic anhydrase | environmental stress response | T690A | *230Y | 690 | 75-544 | MHI3724, |
| *can* | Carbonic anhydrase | environmental stress response | A690T | *230Y | 690 | 16-396 | MHI4390, |
| *can* | Carbonic anhydrase | environmental stress response | A690T | *230Y | 690 | 29-199 | MHI3544, MHI3547, |
| *can* | Carbonic anhydrase | environmental stress response | T690A | *230Y | 690 | 52-105 | MHI3628, MHI3629, MHI3633, |

| | | | | | | | |
|---|---|---|---|---|---|---|---|
| *can* | Carbonic anhydrase | environmental stress response | T690A | *230Y | 690 | 61-1012 | MHI3658, |
| *can* | Carbonic anhydrase | environmental stress response | T690A | *230Y | 690 | 385545 | MHI3965, |
| *can* | Carbonic anhydrase | environmental stress response | G704T | S235* | 704 | 29-1030 | MHI3448, |
| *can* | Carbonic anhydrase | environmental stress response | T707C | D236G | 707 | 29-1030 | MHI3432, MHI3448, |
| *cbiM* | Cobalt transport protein | transporter | G130A | A44T | 130 | 52-105 | MHI3634, |
| *ccmC* | Heme exporter protein C | transporter | T361A | T121S | 361 | 36-36 | MHI4152, |
| *ccmC* | Heme exporter protein C | transporter | G731A | T244I | 731 | 27-683 | MHI3535, |
| *ccmC* | Heme exporter protein C | transporter | T734A | L245Q | 734 | 52-105 | MHI3637, |
| *ccmC* | Heme exporter protein C | transporter | A736T | K246* | 736 | 52-105 | MHI3637, |
| *chuR 1* | Regulator of arylsulfatase activity | environmental stress response | A225T | R75S | 225 | 20-946 | MHI4092, |
| *chuR 1* | Regulator of arylsulfatase activity | environmental stress response | A240C | *80Y | 240 | 20-946 | MHI4092, |
| *chuR 2* | Regulator of arylsulfatase activity | environmental stress response | T218A | Y73F | 218 | 46-266 | MHI3489, |
| *chuR 2* | Regulator of arylsulfatase activity | environmental stress response | T218A | Y73F | 218 | 29-266 | MHI3441, |
| *chuR 2* | Regulator of arylsulfatase activity | environmental stress response | T225A | R75S | 225 | 46-266 | MHI3489, |
| *chuR 2* | Regulator of arylsulfatase activity | environmental stress response | T225A | R75S | 225 | 29-266 | MHI3441, |
| *citC* | [Citrate [pro-3S]-lyase] ligase | transcription/translation | C1001A | R334L | 1001 | 43-836 | MHI3466, MHI3467, MHI3478, MHI3479, MHI4406, |
| *citG* | 2-(5''-triphosphoribosyl)-3'-dephosphocoenzyme-A synthase | transcription/translation | G3T | L1F | 3 | 27-683 | MHI3526, |
| *citG* | 2-(5''-triphosphoribosyl)-3'-dephosphocoenzyme-A synthase | transcription/translation | C3G | L1F | 3 | 52-105 | MHI3636, MHI3637, MHI3640, MHI3645, MHI3646, MHI3648, |
| *cntO* | Metal-pseudopaline receptor | transporter | A142C | L48V | 142 | 13-2156 | MHI4025, |
| *cntO* | Metal-pseudopaline receptor | transporter | T144A | L48F | 144 | 13-2156 | MHI4025, |
| *cntO* | Metal-pseudopaline receptor | transporter | G148C | L50V | 148 | 13-2156 | MHI4025, MHI4049, |
| *cntO* | Metal-pseudopaline receptor | transporter | T160C | N54D | 160 | 13-2156 | MHI4049, |
| *cntO* | Metal-pseudopaline receptor | transporter | T166C | T56A | 166 | 13-2156 | MHI4043, MHI4049, |
| *coaBC* | Coenzyme A biosynthesis bifunctional protein | carbohydrate metabilism | T785C | L262P | 785 | 52-105 | MHI3636, |
| *coaBC* | Coenzyme A biosynthesis bifunctional protein | carbohydrate metabilism | G787C | E263Q | 787 | 52-105 | MHI3636, |
| *copA* | Copper-exporting P-type ATPase | transporter | T5C | L2P | 5 | 2-1040 | MHI4375, |
| *copA* | Copper-exporting P-type ATPase | transporter | T55C | T19A | 55 | 1-266 | MHI3302, |
| *copA* | Copper-exporting P-type ATPase | transporter | T55C | T19A | 55 | 15-2957 | MHI4072, |
| *copA* | Copper-exporting P-type ATPase | transporter | T55C | T19A | 55 | 27-494 | MHI4394, |
| *copA* | Copper-exporting P-type ATPase | transporter | T55C | T19A | 55 | 2954 | MHI3408, |
| *copA* | Copper-exporting P-type ATPase | transporter | A2165C | F722C | 2165 | 39-159 | MHI4161, |
| *copA* | Copper-exporting P-type ATPase | transporter | G2166T | F722L | 2166 | 39-159 | MHI4161, |
| *cpxR* | Transcriptional regulatory protein | regulator | A475T | F159I | 475 | 52-105 | MHI3645, |
| *cpxR* | Transcriptional regulatory protein | regulator | A476T | F159Y | 476 | 52-105 | MHI3645, |
| *csdA* | Cysteine desulfurase | environmental stress response | C417A | E139D | 417 | 13-2156 | MHI4029, MHI4040, MHI4045, |
| *csdE* | Cysteine desulfurase | environmental stress response | G281A | T94I | 281 | 57-161 | MHI4257, MHI4258, MHI4259, |
| *csdE* | Cysteine desulfurase | environmental stress response | A284G | V95A | 284 | 57-161 | MHI4257, MHI4258, MHI4259, |
| *csdE* | Cysteine desulfurase | environmental stress response | T314C | H105R | 314 | 57-161 | MHI4259, |
| *cydD* | ATP-binding/permease protein CydD | transporter | T115G | F39V | 115 | 1-266 | MHI4370, MHI4371, MHI4373, |
| *cydD* | ATP-binding/permease protein CydD | transporter | A115C | F39V | 115 | 64-2958 | MHI4331, |
| *cydD* | ATP-binding/permease protein CydD | transporter | A115C | F39V | 115 | 59-280 | MHI4265, |
| *cydD* | ATP-binding/permease protein CydD | transporter | T115G | F39V | 115 | 7-159 | MHI4382, |
| *cydD* | ATP-binding/permease protein CydD | transporter | T115G | F39V | 115 | 46-266 | MHI4407, |
| *cydD* | ATP-binding/permease protein CydD | transporter | T115G | F39V | 115 | 68-264 | MHI4345, MHI4351, MHI4357, |
| *dacA* | D-alanyl-D-alanine carboxypeptidase DacA | transporter | T1177C | S393P | 1177 | 61-1012 | MHI3656, MHI3661, MHI3667, |
| *dacA* | D-alanyl-D-alanine carboxypeptidase DacA | transporter | C1178A | S393* | 1178 | 61-1012 | MHI3656, MHI3661, MHI3667, MHI3669, |
| *dacA* | D-alanyl-D-alanine carboxypeptidase DacA | transporter | T1180A | *394K | 1180 | 61-1012 | MHI3656, MHI3661, MHI3667, MHI3669, |
| *dam* | DNA adenine methylase | recombination | T854C | H285R | 854 | 64-2958 | MHI4331, |
| *dam* | DNA adenine methylase | recombination | T854C | H285R | 854 | 75-544 | MHI3730, MHI3737, MHI3740, |
| *dam* | DNA adenine methylase | recombination | T854C | H285R | 854 | 15-1238 | MHI4068, |
| *dam* | DNA adenine methylase | recombination | A854G | H285R | 854 | 15-2957 | MHI4061, MHI4067, MHI4075, MHI4076, MHI4086, |
| *dam* | DNA adenine methylase | recombination | T854C | H285R | 854 | 2-1040 | MHI3357, |
| *dam* | DNA adenine methylase | recombination | A854G | H285R | 854 | 2-544 | MHI3351, |
| *dam* | DNA adenine methylase | recombination | A854G | H285R | 854 | 29-536 | MHI3557, MHI3559, MHI3570, |
| *dam* | DNA adenine methylase | recombination | A854G | H285R | 854 | 59-1409 | MHI4267, |
| *dam* | DNA adenine methylase | recombination | A854G | H285R | 854 | 68-264 | MHI4333, MHI4345, MHI4353, MHI4354, MHI4357, |
| *dam* | DNA adenine methylase | recombination | A854G | H285R | 854 | 385545 | MHI3964, MHI3975, |
| *dapD* | 2,3,4,5-tetrahydropyridine-2,6-dicarboxylate N-succinyltransferase | transcription/translation | T829A | *277K | 829 | 52-105 | MHI3647, |
| *dcuD* | Putative cryptic C4-dicarboxylate transporter | transporter | A1292G | L431S | 1292 | 36-36 | MHI4152, |
| *dcuD* | Putative cryptic C4-dicarboxylate transporter | transporter | A1294T | *432K | 1294 | 36-36 | MHI4127, MHI4152, MHI4158, |
| *dmsD* | Tat proofreading chaperone | transcription/translation | A157T | Y53N | 157 | 52-105 | MHI3632, |
| *dmsD* | Tat proofreading chaperone | transcription/translation | C181T | E61K | 181 | 52-105 | MHI3632, |
| *dtd* | D-aminoacyl-tRNA deacylase | transcription/translation | A327T | F109L | 327 | 75-544 | MHI3730, |
| *dtd* | D-aminoacyl-tRNA deacylase | transcription/translation | T348A | K116N | 348 | 75-544 | MHI3730, |
| *dtd* | D-aminoacyl-tRNA deacylase | transcription/translation | A348T | K116N | 348 | 2-544 | MHI3351, |
| *dtd* | D-aminoacyl-tRNA deacylase | transcription/translation | T351A | L117F | 351 | 75-544 | MHI3730, |
| *dtd* | D-aminoacyl-tRNA deacylase | transcription/translation | G352C | P118A | 352 | 75-544 | MHI3730, |
| *dtd* | D-aminoacyl-tRNA deacylase | transcription/translation | G353A | P118L | 353 | 75-544 | MHI3730, |
| *envC* | Protease with a role in cell division | cell wall biosynthesis | A28T | T10S | 28 | 61-1012 | MHI3667, |
| *era* | GTPase Era | transcription/translation | T896G | Y299S | 896 | 2-544 | MHI3351, |
| *era* | GTPase Era | transcription/translation | A896C | Y299S | 896 | 61-1012 | MHI3656, |
| *era* | GTPase Era | transcription/translation | A899T | M300K | 899 | 2-544 | MHI3351, |
| *era* | GTPase Era | transcription/translation | T899A | M300K | 899 | 61-1012 | MHI3656, |
| *era* | GTPase Era | transcription/translation | G901A | D301N | 901 | 75-544 | MHI3740, |
| *era* | GTPase Era | transcription/translation | C901T | D301N | 901 | 2-544 | MHI3351, |
| *era* | GTPase Era | transcription/translation | G901A | D301N | 901 | 61-1012 | MHI3656, |
| *esiB* | 3-phosphoshikimate 1-carboxyvinyltransferase (IgA Bingind) | virulence | A24T | L8F | 24 | 1-266 | MHI3574, MHI4370, |
| *esiB* | 3-phosphoshikimate 1-carboxyvinyltransferase (IgA Bingind) | virulence | A24T | L8F | 24 | 46-266 | MHI3489, MHI3490, MHI3494, |
| *esiB* | 3-phosphoshikimate 1-carboxyvinyltransferase (IgA Bingind) | virulence | A24T | L8F | 24 | 29-266 | MHI3430, MHI3431, MHI3443, |
| *esiB* | 3-phosphoshikimate 1-carboxyvinyltransferase (IgA Bingind) | virulence | C180A | E60D | 180 | 29-536 | MHI3557, |
| *esiB* | 3-phosphoshikimate 1-carboxyvinyltransferase (IgA Bingind) | virulence | A193C | F65V | 193 | 29-536 | MHI3557, |
| *esiB* | 3-phosphoshikimate 1-carboxyvinyltransferase (IgA Bingind) | virulence | C520A | D174Y | 520 | 16-396 | MHI3385, |
| *esiB* | 3-phosphoshikimate 1-carboxyvinyltransferase (IgA Bingind) | virulence | G565A | H189Y | 565 | 39-159 | MHI4184, |
| *esiB* | 3-phosphoshikimate 1-carboxyvinyltransferase (IgA Bingind) | virulence | T571C | K191E | 571 | 39-159 | MHI3963, MHI3965, MHI4184, |
| *esiB* | 3-phosphoshikimate 1-carboxyvinyltransferase (IgA Bingind) | virulence | C573A | K191N | 573 | 39-159 | MHI4184, |
| *esiB* | 3-phosphoshikimate 1-carboxyvinyltransferase (IgA Bingind) | virulence | T581A | Y194F | 581 | 39-159 | MHI4184, |
| *esiB* | 3-phosphoshikimate 1-carboxyvinyltransferase (IgA Bingind) | virulence | C601A | A201S | 601 | 39-159 | MHI4184, |
| *esiB* | 3-phosphoshikimate 1-carboxyvinyltransferase (IgA Bingind) | virulence | G602T | A201D | 602 | 39-159 | MHI4184, |
| *esiB* | 3-phosphoshikimate 1-carboxyvinyltransferase (IgA Bingind) | virulence | T610A | K204* | 610 | 73-1030 | MHI3686, |
| *esiB* | 3-phosphoshikimate 1-carboxyvinyltransferase (IgA Bingind) | virulence | C611T | R204H | 611 | 39-159 | MHI4184, |
| *esiB* | 3-phosphoshikimate 1-carboxyvinyltransferase (IgA Bingind) | virulence | C612A | K204N | 612 | 73-1030 | MHI3686, |
| *esiB* | 3-phosphoshikimate 1-carboxyvinyltransferase (IgA Bingind) | virulence | A618T | N206K | 618 | 73-1030 | MHI3686, |
| *esiB* | 3-phosphoshikimate 1-carboxyvinyltransferase (IgA Bingind) | virulence | T626C | D209G | 626 | 73-1030 | MHI3686, |
| *esiB* | 3-phosphoshikimate 1-carboxyvinyltransferase (IgA Bingind) | virulence | A628C | Y210D | 628 | 39-159 | MHI4184, |
| *esiB* | 3-phosphoshikimate 1-carboxyvinyltransferase (IgA Bingind) | virulence | T628G | S210R | 628 | 73-1030 | MHI3686, |
| *esiB* | 3-phosphoshikimate 1-carboxyvinyltransferase (IgA Bingind) | virulence | C629T | S210N | 629 | 73-1030 | MHI3686, |
| *esiB* | 3-phosphoshikimate 1-carboxyvinyltransferase (IgA Bingind) | virulence | T637C | I213V | 637 | 73-1030 | MHI3686, |
| *esiB* | 3-phosphoshikimate 1-carboxyvinyltransferase (IgA Bingind) | virulence | T649C | S217G | 649 | 73-1030 | MHI3686, |
| *esiB* | 3-phosphoshikimate 1-carboxyvinyltransferase (IgA Bingind) | virulence | C892A | V298F | 892 | 73-1030 | MHI3686, |
| *esiB* | 3-phosphoshikimate 1-carboxyvinyltransferase (IgA Bingind) | virulence | G906T | N302K | 906 | 73-1030 | MHI3686, |
| *esiB* | 3-phosphoshikimate 1-carboxyvinyltransferase (IgA Bingind) | virulence | C917T | R306K | 917 | 73-1030 | MHI3686, |
| *esiB* | 3-phosphoshikimate 1-carboxyvinyltransferase (IgA Bingind) | virulence | T14C | K5R | 14 | 20-946 | MHI4091, MHI4092, |
| *esiB* | 3-phosphoshikimate 1-carboxyvinyltransferase (IgA Bingind) | virulence | T24A | L8F | 24 | 20-946 | MHI4091, |
| *esiB* | 3-phosphoshikimate 1-carboxyvinyltransferase (IgA Bingind) | virulence | T39G | L13F | 39 | 20-946 | MHI4091, |
| *esiB* | 3-phosphoshikimate 1-carboxyvinyltransferase (IgA Bingind) | virulence | T610G | N204H | 610 | 20-946 | MHI4101, |
| *esiB* | 3-phosphoshikimate 1-carboxyvinyltransferase (IgA Bingind) | virulence | T616C | K206E | 616 | 20-946 | MHI4101, |
| *esiB* | 3-phosphoshikimate 1-carboxyvinyltransferase (IgA Bingind) | virulence | C618A | K206N | 618 | 20-946 | MHI4101, |
| *fbp* | Fructose-1,6-bisphosphatase class 1 | carbohydrate metabilism | G514T | H172N | 514 | 29-199 | MHI3547, |
| *fbp* | Fructose-1,6-bisphosphatase class 1 | carbohydrate metabilism | T691G | N231H | 691 | 29-199 | MHI3547, |
| *fbp* | Fructose-1,6-bisphosphatase class 1 | carbohydrate metabilism | G704A | A235V | 704 | 29-199 | MHI3547, |

| | | | | | | | |
|---|---|---|---|---|---|---|---|
| *fbp* | Fructose-1,6-bisphosphatase class 1 | carbohydrate metabilism | C898G | E300Q | 898 | 29-199 | MHI3547, |
| *fbp* | Fructose-1,6-bisphosphatase class 1 | carbohydrate metabilism | G908T | A303E | 908 | 29-199 | MHI3547, |
| *fbp* | Fructose-1,6-bisphosphatase class 1 | carbohydrate metabilism | C916G | E306Q | 916 | 29-199 | MHI3547, |
| *fbp* | Fructose-1,6-bisphosphatase class 1 | carbohydrate metabilism | G923A | A308V | 923 | 29-199 | MHI3547, MHI3635, MHI3646, MHI3647, |
| *fbp* | Fructose-1,6-bisphosphatase class 1 | carbohydrate metabilism | G3A | M1I | 3 | 73-1030 | MHI3704, MHI3705, |
| *fbp* | Fructose-1,6-bisphosphatase class 1 | carbohydrate metabilism | C3T | M1I | 3 | 29-199 | MHI3548, |
| *fbp* | Fructose-1,6-bisphosphatase class 1 | carbohydrate metabilism | C3T | M1I | 3 | 52-105 | MHI3626, MHI3627, MHI3631, |
| *fbp* | Fructose-1,6-bisphosphatase class 1 | carbohydrate metabilism | G3A | M1I | 3 | 61-568 | MHI3653, MHI3659, |
| *fbpC2* | Fe(3+) ions import ATP-binding protein FbpC 2 | iron metabolism | T63A | F21L | 63 | 10 | MHI3997, |
| *fbpC2* | Fe(3+) ions import ATP-binding protein FbpC 2 | iron metabolism | A63T | F21L | 63 | 2-1040 | MHI3346, MHI3352, |
| *fbpC2* | Fe(3+) ions import ATP-binding protein FbpC 2 | iron metabolism | A64C | N22H | 64 | 10 | MHI3997, |
| *fbpC2* | Fe(3+) ions import ATP-binding protein FbpC 2 | iron metabolism | T64G | N22H | 64 | 2-1040 | MHI3346, |
| *fbpC2* | Fe(3+) ions import ATP-binding protein FbpC 2 | iron metabolism | A65C | N22T | 65 | 10-57 | MHI3997, |
| *fbpC2* | Fe(3+) ions import ATP-binding protein FbpC 2 | iron metabolism | T66G | N22K | 66 | 10-57 | MHI3997, |
| *fbpC2* | Fe(3+) ions import ATP-binding protein FbpC 2 | iron metabolism | C887A | S296* | 887 | 46-266 | MHI3491, |
| *fbpC2* | Fe(3+) ions import ATP-binding protein FbpC 2 | iron metabolism | C887A | S296* | 887 | 29-199 | MHI3544, MHI3547, MHI3558, MHI3563, |
| *fbpC2* | Fe(3+) ions import ATP-binding protein FbpC 2 | iron metabolism | C887A | S296* | 887 | 29-266 | MHI3449, |
| *fbpC2* | Fe(3+) ions import ATP-binding protein FbpC 2 | iron metabolism | C895T | P299S | 895 | 46-266 | MHI3491, |
| *fbpC2* | Fe(3+) ions import ATP-binding protein FbpC 2 | iron metabolism | C895T | P299S | 895 | 73-1030 | MHI3710, MHI3711, MHI3713, |
| *fbpC2* | Fe(3+) ions import ATP-binding protein FbpC 2 | iron metabolism | G895A | P299S | 895 | 29-1030 | MHI3448, |
| *fbpC2* | Fe(3+) ions import ATP-binding protein FbpC 2 | iron metabolism | C895T | P299S | 895 | 29-199 | MHI3544, MHI3547, MHI3552, MHI3558, MHI3561, MHI3563, |
| *fbpC2* | Fe(3+) ions import ATP-binding protein FbpC 2 | iron metabolism | C895T | P299S | 895 | 29-266 | MHI3438, MHI3441, MHI3443, MHI3449, |
| *fbpC2* | Fe(3+) ions import ATP-binding protein FbpC 2 | iron metabolism | T911G | N304T | 911 | 20-946 | MHI4091, MHI4092, |
| *fhuD* | Fumarate and nitrate reduction regulatory protein | regulator | T8A | K3I | 8 | 27-683 | MHI3525, MHI3532, |
| *fnr* | Fumarate and nitrate reduction regulatory protein | regulator | T746G | N249T | 746 | 52-105 | MHI3647, |
| *fnr* | Fumarate and nitrate reduction regulatory protein | regulator | T756G | K252N | 756 | 52-105 | MHI3647, |
| *fnr* | Fumarate and nitrate reduction regulatory protein | regulator | T767A | V256D | 767 | 56-344 | MHI4213, |
| *ftsH_1* | ATP-dependent zinc metalloprotease | cell wall biosynthesis | G1870A | E624K | 1870 | 61-1012 | MHI3661, |
| *ftsH_1* | ATP-dependent zinc metalloprotease | cell wall biosynthesis | A1892C | D631A | 1892 | 61-1012 | MHI3661, |
| *ftsH_1* | ATP-dependent zinc metalloprotease | cell wall biosynthesis | A1894G | N632D | 1894 | 61-1012 | MHI3661, MHI4390, |
| *ftsH_1* | ATP-dependent zinc metalloprotease | cell wall biosynthesis | A1895T | N632I | 1895 | 61-1012 | MHI3661, |
| *ftsH_2* | ATP-dependent zinc metalloprotease | cell wall biosynthesis | A1627C | S543A | 1627 | 13-2156 | MHI4025, |
| *ftsH_2* | ATP-dependent zinc metalloprotease | cell wall biosynthesis | G1870A | E624K | 1870 | 10-57 | MHI4003, |
| *ftsH_2* | ATP-dependent zinc metalloprotease | cell wall biosynthesis | A1892C | D631A | 1892 | 10-57 | MHI4003, |
| *ftsI* | Peptidoglycan D,D-transpeptidase | cell wall biosynthesis | G166T | G56C | 166 | 39-159 | MHI4162, MHI4171, MHI4184, MHI4191, |
| *ftsI* | Peptidoglycan D,D-transpeptidase | cell wall biosynthesis | G224A | S75N | 224 | 39-159 | MHI4162, MHI4171, MHI4184, MHI4191, |
| *ftsI* | Peptidoglycan D,D-transpeptidase | cell wall biosynthesis | C364A | H122N | 364 | 39-159 | MHI4162, MHI4171, MHI4191, |
| *ftsI* | Peptidoglycan D,D-transpeptidase | cell wall biosynthesis | T374C | V125A | 374 | 39-159 | MHI4162, MHI4171, MHI4191, |
| *ftsI* | Peptidoglycan D,D-transpeptidase | cell wall biosynthesis | T391G | S131A | 391 | 39-159 | MHI4162, MHI4171, MHI4191, |
| *ftsI* | Peptidoglycan D,D-transpeptidase | cell wall biosynthesis | C403G | Q135E | 403 | 39-159 | MHI4162, MHI4171, MHI4191, |
| *ftsI* | Peptidoglycan D,D-transpeptidase | cell wall biosynthesis | G454T | A152S | 454 | 39-159 | MHI4162, MHI4171, MHI4191, |
| *ftsI* | Peptidoglycan D,D-transpeptidase | cell wall biosynthesis | C494T | S165L | 494 | 39-159 | MHI4162, MHI4171, MHI4191, |
| *ftsI* | Peptidoglycan D,D-transpeptidase | cell wall biosynthesis | A497G | N166S | 497 | 39-159 | MHI4162, MHI4171, MHI4191, |
| *ftsI* | Peptidoglycan D,D-transpeptidase | cell wall biosynthesis | G1578T | K526N | 1578 | 39-159 | MHI4162, MHI4171, MHI4184, MHI4191, |
| *ftsI* | Peptidoglycan D,D-transpeptidase | cell wall biosynthesis | G1652A | G551D | 1652 | 39-159 | MHI4162, MHI4171, MHI4184, MHI4191, |
| *ftsN* | Cell division protein | cell wall biosynthesis | A41C | N14T | 41 | 10-57 | MHI3988, |
| *glmM* | Phosphoglucosamine mutase | carbohydrate metabilism | A1T | M1L | 1 | 2-1040 | MHI3352, MHI3357, |
| *glmM* | Phosphoglucosamine mutase | carbohydrate metabilism | T2C | M1T | 2 | 2-1040 | MHI3352, |
| *glnE* | Bifunctional glutamine synthetase adenylyltransferase/adenylyl-removing enzyme | environmental stress response | C1363T | D455N | 1363 | 27-683 | MHI3541, |
| *glnE* | Bifunctional glutamine synthetase adenylyltransferase/adenylyl-removing enzyme | environmental stress response | G1380T | L460F | 1380 | 29-199 | MHI3563, |
| *glpQ* | Glycerophosphodiester phosphodiesterase | glycolipid metabolism | T1094G | *365S | 1094 | 61-1012 | MHI3656, MHI3661, |
| *glpQ* | Glycerophosphodiester phosphodiesterase | glycolipid metabolism | G1066T | G356C | 1066 | 52-105 | MHI3623, MHI3640, |
| *glpQ* | Glycerophosphodiester phosphodiesterase | glycolipid metabolism | T238G | C80G | 238 | 52-105 | MHI3642, |
| *gltC* | HTH-type transcriptional regulator GltC | regulator | C4T | E2K | 4 | 27-494 | MHI3417, |
| *gmhA_2* | Phosphoheptose isomerase | carbohydrate metabilism | C471A | K157N | 471 | 61-568 | MHI3673, MHI3682, |
| *gor* | Glutathione reductase | environmental stress response | G310T | A104S | 310 | 20-946 | MHI4092, |
| *gor* | Glutathione reductase | environmental stress response | T321A | N107K | 321 | 20-946 | MHI4091, MHI4092, |
| *gpt_1* | Xanthine-guanine phosphoribosyltransferase | transcription/translation | T53C | K18R | 53 | 52-105 | MHI3635, |
| *gpt_2* | Xanthine-guanine phosphoribosyltransferase | transcription/translation | C53T | R18K | 53 | 52-105 | MHI3640, MHI3644, MHI3649, MHI3650, |
| *gpt_2* | Xanthine-guanine phosphoribosyltransferase | transcription/translation | A468T | *156Y | 468 | 27-683 | MHI3526, MHI3533, MHI3535, |
| *gpt_2* | Xanthine-guanine phosphoribosyltransferase | transcription/translation | T469C | F157L | 469 | 27-683 | MHI3526, MHI3535, |
| *greA* | Transcription elongation factor GreA | transcription/translation | T467C | N156S | 467 | 29-199 | MHI3568, |
| *greA* | Transcription elongation factor GreA | transcription/translation | A469T | Y157N | 469 | 36-36 | MHI4158, |
| *greA* | Transcription elongation factor GreA | transcription/translation | A469T | Y157N | 469 | 15-2957 | MHI4088, |
| *greA* | Transcription elongation factor GreA | transcription/translation | T470G | Y157S | 470 | 29-199 | MHI3568, |
| *greA* | Transcription elongation factor GreA | transcription/translation | A471T | Y157* | 471 | 36-36 | MHI4158, |
| *greA* | Transcription elongation factor GreA | transcription/translation | A471T | Y157* | 471 | 15-2957 | MHI4088, |
| *greA* | Transcription elongation factor GreA | transcription/translation | T471A | Y157* | 471 | 2-1040 | MHI3346, |
| *greA* | Transcription elongation factor GreA | transcription/translation | A473G | I158T | 473 | 36-36 | MHI4158, |
| *greA* | Transcription elongation factor GreA | transcription/translation | A473G | I158T | 473 | 15-2957 | MHI4077, MHI4088, |
| *greA* | Transcription elongation factor GreA | transcription/translation | T473C | I158T | 473 | 2-1040 | MHI3346, |
| *greA* | Transcription elongation factor GreA | transcription/translation | T473A | I158N | 473 | 29-266 | MHI3450, |
| *greA* | Transcription elongation factor GreA | transcription/translation | A475C | *159E | 475 | 36-36 | MHI4158, |
| *greA* | Transcription elongation factor GreA | transcription/translation | A475C | *159E | 475 | 15-2957 | MHI4065, MHI4077, MHI4088, |
| *greA* | Transcription elongation factor GreA | transcription/translation | T475G | *159E | 475 | 2-1040 | MHI3346, |
| *greA* | Transcription elongation factor GreA | transcription/translation | T475G | *159E | 475 | 29-266 | MHI3450, |
| *greA* | Transcription elongation factor GreA | transcription/translation | A475C | *159E | 475 | 59-1409 | MHI4283, |
| *greA* | Transcription elongation factor GreA | transcription/translation | T477A | *159Y | 477 | 20-946 | MHI4104, |
| *greA* | Transcription elongation factor GreA | transcription/translation | C493A | G165* | 493 | 68-264 | MHI4334, MHI4335, MHI4336, MHI4337, MHI4339, MHI4340, MHI4341, MHI4346, MHI4347, MHI4348, MHI4349, MHI4350, MHI4351, MHI4352, MHI4359, MHI4360, MHI4361, MHI4362, MHI4363, MHI4364, |
| *greA* | Transcription elongation factor GreA | transcription/translation | C494A | G165V | 494 | 68-264 | MHI4357, |
| *hap* | Adhesion and penetration protein autotransporter | transporter | C3203T | P1068L | 3203 | 385545 | MHI3965, |
| *hindIIIM* | Type II methyltransferase | recombination | A53T | D18V | 53 | 57-161 | MHI4244, |
| *hindIIIR* | Type-2 restriction enzyme HindIII | recombination | T632A | N211I | 632 | 20-946 | MHI4091, |
| *hindIIIR* | Type-2 restriction enzyme HindIII | recombination | T710G | K237T | 710 | 7-159 | MHI3605, |
| *hindIIIR* | Type-2 restriction enzyme HindIII | recombination | A733G | R245G | 733 | 29-266 | MHI3615, |
| *hindIIIR* | Type-2 restriction enzyme HindIII | recombination | C734A | R245I | 734 | 20-946 | MHI4092, |
| *hsdR* | Type I restriction enzyme EcoR124II R protein | recombination | T566G | F189C | 566 | 13-2156 | MHI4025, |
| *hsdR* | Type I restriction enzyme EcoR124II R protein | recombination | T566G | F189C | 566 | 46-266 | MHI3489, |
| *hsdR* | Type I restriction enzyme EcoR124II R protein | recombination | T566G | F189C | 566 | 2-544 | MHI3354, |
| *hsdR* | Type I restriction enzyme EcoR124II R protein | recombination | T566G | F189C | 566 | 29-266 | MHI3431, MHI3441, |
| *hsdR* | Type I restriction enzyme EcoR124II R protein | recombination | T566G | F189C | 566 | 56-344 | MHI4198, |
| *hsdR* | Type I restriction enzyme EcoR124II R protein | recombination | A568T | N190Y | 568 | 27-494 | MHI3418, |
| *hsdR* | Type I restriction enzyme EcoR124II R protein | recombination | A568T | N190Y | 568 | 59-1409 | MHI4269, |
| *hsdR* | Type I restriction enzyme EcoR124II R protein | recombination | A568T | N190Y | 568 | 68-264 | MHI4353, |
| *hsdR* | Type I restriction enzyme EcoR124II R protein | recombination | T590A | K197I | 590 | 57-161 | MHI4255, |
| *hsdR* | Type I restriction enzyme EcoR124II R protein | recombination | T590A | K197I | 590 | 75-544 | MHI3729, |
| *hsdR* | Type I restriction enzyme EcoR124II R protein | recombination | A590G | K197R | 590 | 27-494 | MHI3418, |
| *hsdR* | Type I restriction enzyme EcoR124II R protein | recombination | A590T | K197I | 590 | 29-266 | MHI3440, |
| *hsdR* | Type I restriction enzyme EcoR124II R protein | recombination | A590G | K197R | 590 | 56-344 | MHI4213, |
| *hsdR* | Type I restriction enzyme EcoR124II R protein | recombination | A590G | K197R | 590 | 68-264 | MHI4353, |
| *hsdR* | Type I restriction enzyme EcoR124II R protein | recombination | A592T | Y198N | 592 | 75-544 | MHI3729, |
| *hsdR* | Type I restriction enzyme EcoR124II R protein | recombination | T592A | Y198N | 592 | 29-266 | MHI3440, |
| *hsdR* | Type I restriction enzyme EcoR124II R protein | recombination | T592A | Y198N | 592 | 59-1409 | MHI4269, |
| *hxuC* | Heme/hemopexin utilization protein C | iron metabolism | A1673G | N558S | 1673 | 1-266 | MHI4370, |
| *hxuC* | Heme/hemopexin utilization protein C | iron metabolism | A1678G | S560G | 1678 | 1-266 | MHI4370, |
| *hxuC* | Heme/hemopexin utilization protein C | iron metabolism | A1678G | S560G | 1678 | 46-266 | MHI3494, |
| *hxuC* | Heme/hemopexin utilization protein C | iron metabolism | A1684G | S562G | 1684 | 1-266 | MHI4370, |
| *hxuC* | Heme/hemopexin utilization protein C | iron metabolism | A1684G | S562G | 1684 | 46-266 | MHI3490, MHI3494, |
| *iaaH* | aminobenzoyl-glutamate utilization protein | transcription/translation | T5C | N2S | 5 | 29-199 | MHI3552, |
| *iaaH* | aminobenzoyl-glutamate utilization protein | transcription/translation | A5G | N2S | 5 | 61-1012 | MHI3671, |

| | | | | | | | |
|---|---|---|---|---|---|---|---|
| *iaaH* | aminobenzoyl-glutamate utilization protein | transcription/translation | T7A | L3I | 7 | 61-1012 | MHI3671, |
| *iaaH* | aminobenzoyl-glutamate utilization protein | transcription/translation | T8G | L3* | 8 | 61-1012 | MHI3671, |
| *iaaH* | aminobenzoyl-glutamate utilization protein | transcription/translation | A9C | L3F | 9 | 61-1012 | MHI3671, |
| *iga* | Immunoglobulin A1 protease autotransporter | virulence | A1012T | S338T | 1012 | 16-396 | MHI3611, |
| *iga* | Immunoglobulin A1 protease autotransporter | virulence | G1013T | S338Y | 1013 | 16-396 | MHI3611, |
| *iga* | Immunoglobulin A1 protease autotransporter | virulence | G1015C | Q339E | 1015 | 16-396 | MHI3611, |
| *ihfB* | Integration host factor subunit beta | recombination | G244A | V82M | 244 | 73-1030 | MHI3698, MHI3705, MHI3707, |
| *kdgK* | 2-dehydro-3-deoxygluconokinase | carbohydrate metabilism | T423G | L141F | 423 | 52-105 | MHI3634, MHI3635, MHI3645, MHI3646, MHI3649, |
| *kdgK* | 2-dehydro-3-deoxygluconokinase | carbohydrate metabilism | T427C | K143E | 427 | 61-568 | MHI3670, |
| *kdgK* | 2-dehydro-3-deoxygluconokinase | carbohydrate metabilism | T430A | N144Y | 430 | 52-105 | MHI3645, |
| *kdgK* | 2-dehydro-3-deoxygluconokinase | carbohydrate metabilism | T431A | N144I | 431 | 52-105 | MHI3634, MHI3635, MHI3645, MHI3649, |
| *kdgK* | 2-dehydro-3-deoxygluconokinase | carbohydrate metabilism | T431G | N144T | 431 | 52-105 | MHI3634, MHI3635, MHI3645, MHI3649, |
| *kdgK* | 2-dehydro-3-deoxygluconokinase | carbohydrate metabilism | T431G | N144T | 431 | 61-568 | MHI3670, |
| *kdgK* | 2-dehydro-3-deoxygluconokinase | carbohydrate metabilism | C451A | E151* | 451 | 52-105 | MHI3634, MHI3635, MHI3645, MHI3646, MHI3648, |
| *kdgK* | 2-dehydro-3-deoxygluconokinase | carbohydrate metabilism | C451A | E151* | 451 | 61-568 | MHI3670, |
| *kdgK* | 2-dehydro-3-deoxygluconokinase | carbohydrate metabilism | T455G | Q152P | 455 | 61-568 | MHI3670, |
| *kdgK* | 2-dehydro-3-deoxygluconokinase | carbohydrate metabilism | T456A | Q152H | 456 | 61-568 | MHI3670, |
| *kdgK* | 2-dehydro-3-deoxygluconokinase | carbohydrate metabilism | T459A | L153F | 459 | 61-568 | MHI3670, |
| *kdkA* | 3-deoxy-D-manno-octulosonic acid kinase | cell wall biosynthesis | T39A | F13L | 39 | 15-1238 | MHI4074, |
| *kdkA* | 3-deoxy-D-manno-octulosonic acid kinase | cell wall biosynthesis | A39T | F13L | 39 | 2-544 | MHI3351, |
| *lapA* | Lipopolysaccharide assembly protein A | glycolipid metabolism | T50G | V17G | 50 | 15-1238 | MHI4074, |
| *lapA* | Lipopolysaccharide assembly protein A | glycolipid metabolism | A50C | V17G | 50 | 29-199 | MHI3544, MHI3555, |
| *lapA* | Lipopolysaccharide assembly protein A | glycolipid metabolism | G59A | T20I | 59 | 29-199 | MHI3555, |
| *lapA* | Lipopolysaccharide assembly protein A | glycolipid metabolism | T61C | I21V | 61 | 29-199 | MHI3555, |
| *lapA* | Lipopolysaccharide assembly protein A | glycolipid metabolism | A62C | I21S | 62 | 29-199 | MHI3555, |
| *lepB* | Signal peptidase I | transcription/translation | T98G | L33* | 98 | 27-683 | MHI3535, |
| *lepB* | Signal peptidase I | transcription/translation | G119A | S40L | 119 | 10-57 | MHI4003, |
| *lepB* | Signal peptidase I | transcription/translation | C119T | S40L | 119 | 27-683 | MHI3535, |
| *lepB* | Signal peptidase I | transcription/translation | G119A | S40L | 119 | 29-199 | MHI3558, |
| *lepB* | Signal peptidase I | transcription/translation | C124A | V42L | 124 | 10 | MHI4003, |
| *lepB* | Signal peptidase I | transcription/translation | C124A | V42L | 124 | 29-199 | MHI3558, |
| *lldP* | L-lactate permease | transporter | A1550G | I517T | 1550 | 16-396 | MHI4390, |
| *lldP* | L-lactate permease | transporter | G1594T | L532I | 1594 | 27-494 | MHI3406, |
| *lolB* | Outer-membrane lipoprotein LolB | cell wall biosynthesis | G28A | L10F | 28 | 15-2957 | MHI4078, MHI4081, MHI4086, |
| *lolB* | Outer-membrane lipoprotein LolB | cell wall biosynthesis | A37G | T13A | 37 | 57-161 | MHI4247, MHI4255, MHI4258, MHI4261, |
| *lolB* | Outer-membrane lipoprotein LolB | cell wall biosynthesis | T46A | T16S | 46 | 15-2957 | MHI4078, |
| *lolB* | Outer-membrane lipoprotein LolB | cell wall biosynthesis | G46A | A16T | 46 | 29-536 | MHI3551, MHI3557, MHI3559, MHI3562, MHI3570, |
| *lst* | N-acetyllactosaminide alpha-2,3-sialyltransferase | cell wall biosynthesis | T410A | I137N | 410 | 64-2958 | MHI4304, |
| *lst* | N-acetyllactosaminide alpha-2,3-sialyltransferase | cell wall biosynthesis | T456A | K152N | 456 | 59-1409 | MHI4295, |
| *lst* | N-acetyllactosaminide alpha-2,3-sialyltransferase | cell wall biosynthesis | A460T | I154L | 460 | 57-161 | MHI4261, |
| *lst* | N-acetyllactosaminide alpha-2,3-sialyltransferase | cell wall biosynthesis | A468C | K156N | 468 | 57-161 | MHI4261, |
| *lst* | N-acetyllactosaminide alpha-2,3-sialyltransferase | cell wall biosynthesis | C469A | H157N | 469 | 57-161 | MHI4253, MHI4261, |
| *lst* | N-acetyllactosaminide alpha-2,3-sialyltransferase | cell wall biosynthesis | A475C | T159P | 475 | 57-161 | MHI4261, |
| *lst* | N-acetyllactosaminide alpha-2,3-sialyltransferase | cell wall biosynthesis | C476G | T159R | 476 | 57-161 | MHI4261, |
| *lst* | N-acetyllactosaminide alpha-2,3-sialyltransferase | cell wall biosynthesis | A491T | Y164F | 491 | 57-161 | MHI4261, MHI4261, |
| *lst* | N-acetyllactosaminide alpha-2,3-sialyltransferase | cell wall biosynthesis | A491G | Y164C | 491 | 64-2958 | MHI4304, |
| *lst* | N-acetyllactosaminide alpha-2,3-sialyltransferase | cell wall biosynthesis | T494G | K165T | 494 | 61-568 | MHI3670, |
| *lst* | N-acetyllactosaminide alpha-2,3-sialyltransferase | cell wall biosynthesis | A499T | I167F | 499 | 57-161 | MHI4261, |
| *lst* | N-acetyllactosaminide alpha-2,3-sialyltransferase | cell wall biosynthesis | T594A | L198F | 594 | 15-2957 | MHI4078, |
| *lst* | N-acetyllactosaminide alpha-2,3-sialyltransferase | cell wall biosynthesis | T613C | I205V | 613 | 15-2957 | MHI4078, |
| *lst* | N-acetyllactosaminide alpha-2,3-sialyltransferase | cell wall biosynthesis | A646G | S216P | 646 | 15-2957 | MHI4078, |
| *lst* | N-acetyllactosaminide alpha-2,3-sialyltransferase | cell wall biosynthesis | T898A | I300L | 898 | 27-683 | MHI3529, |
| *lst* | N-acetyllactosaminide alpha-2,3-sialyltransferase | cell wall biosynthesis | A899G | I300T | 899 | 27-683 | MHI3529, |
| *lst* | N-acetyllactosaminide alpha-2,3-sialyltransferase | cell wall biosynthesis | A901T | Y301N | 901 | 27-683 | MHI3529, |
| *lst* | N-acetyllactosaminide alpha-2,3-sialyltransferase | cell wall biosynthesis | A903T | Y301* | 903 | 27-683 | MHI3529, |
| *mdh* | Malate dehydrogenase | carbohydrate metabilism | T918G | E306D | 918 | 57-161 | MHI4243, |
| *mdh* | Malate dehydrogenase | carbohydrate metabilism | T919A | K307* | 919 | 57-161 | MHI4243, |
| *mdh* | Malate dehydrogenase | carbohydrate metabilism | T920A | K307I | 920 | 57-161 | MHI4243, |
| *menE* | 2-succinylbenzoate--CoA ligase | environmental stress response | G1196A | S399N | 1196 | 15-2957 | MHI4075, |
| *menE* | 2-succinylbenzoate--CoA ligase | environmental stress response | T1203A | S401R | 1203 | 15-2957 | MHI4067, MHI4075, |
| *menE* | 2-succinylbenzoate--CoA ligase | environmental stress response | G1221T | M407I | 1221 | 15-2957 | MHI4075, |
| *menE* | 2-succinylbenzoate--CoA ligase | environmental stress response | T1222G | F408V | 1222 | 15-2957 | MHI4075, |
| *mepA* | Penicillin-insensitive murein endopeptidase | cell wall biosynthesis | T25G | T9P | 25 | 57-161 | MHI4255, |
| *mepA* | Penicillin-insensitive murein endopeptidase | cell wall biosynthesis | A25C | T9P | 25 | 73-1030 | MHI3712, |
| *mepA* | Penicillin-insensitive murein endopeptidase | cell wall biosynthesis | C26G | T9S | 26 | 39-159 | MHI4172, |
| *mepA* | Penicillin-insensitive murein endopeptidase | cell wall biosynthesis | G26C | T9S | 26 | 57-161 | MHI4255, MHI4260, |
| *mepA* | Penicillin-insensitive murein endopeptidase | cell wall biosynthesis | C26G | T9S | 26 | 73-1030 | MHI3712, |
| *mepA* | Penicillin-insensitive murein endopeptidase | cell wall biosynthesis | G26C | T9S | 26 | 75-544 | MHI3728, |
| *mepA* | Penicillin-insensitive murein endopeptidase | cell wall biosynthesis | G26C | T9S | 26 | 2-1040 | MHI3353, |
| *mepA* | Penicillin-insensitive murein endopeptidase | cell wall biosynthesis | C26G | T9S | 26 | 52-105 | MHI3635, MHI3646, MHI3647, |
| *mepA* | Penicillin-insensitive murein endopeptidase | cell wall biosynthesis | C50A | S17Y | 50 | 73-1030 | MHI3712, |
| *mepA* | Penicillin-insensitive murein endopeptidase | cell wall biosynthesis | G52T | L18M | 52 | 64-2958 | MHI4331, |
| *mepA* | Penicillin-insensitive murein endopeptidase | cell wall biosynthesis | G52T | L18M | 52 | 2-1040 | MHI3346, |
| *mepA* | Penicillin-insensitive murein endopeptidase | cell wall biosynthesis | A52C | L18M | 52 | 29-598 | MHI3545, MHI3571, |
| *mepA* | Penicillin-insensitive murein endopeptidase | cell wall biosynthesis | C52A | L18M | 52 | 52-105 | MHI3645, |
| *mepA* | Penicillin-insensitive murein endopeptidase | cell wall biosynthesis | G52T | L18M | 52 | 61-1012 | MHI3656, |
| *mglA* | Galactose/methyl galactoside import ATP-binding protein | transporter | T654A | D218E | 654 | 29-598 | MHI3571, |
| *mglA* | Galactose/methyl galactoside import ATP-binding protein | transporter | G655A | V219M | 655 | 29-598 | MHI3571, |
| *mglA* | Galactose/methyl galactoside import ATP-binding protein | transporter | A661G | I221V | 661 | 29-598 | MHI3545, |
| *mlaA* | Intermembrane phospholipid transport system lipoprotein | transporter | C14A | A5E | 14 | 52-105 | MHI3644, MHI3649, |
| *mlaA* | Intermembrane phospholipid transport system lipoprotein | transporter | G14T | A5E | 14 | 59-280 | MHI4272, |
| *mlaA* | Intermembrane phospholipid transport system lipoprotein | transporter | G37T | A13S | 37 | 39-159 | MHI4172, |
| *mlaA* | Intermembrane phospholipid transport system lipoprotein | transporter | G37T | A13S | 37 | 52-105 | MHI3634, MHI3644, MHI3649, |
| *mlaA* | Intermembrane phospholipid transport system lipoprotein | transporter | C37A | A13S | 37 | 59-280 | MHI4272, |
| *mlaA* | Intermembrane phospholipid transport system lipoprotein | transporter | A752T | *251L | 752 | 43-836 | MHI3479, |
| *mlaA* | Intermembrane phospholipid transport system lipoprotein | transporter | T754A | *252K | 754 | 10-57 | MHI3990, MHI4011, |
| *mlaC* | Intermembrane phospholipid transport system binding protein | transporter | G169C | D57H | 169 | 64-2958 | MHI4304, |
| *mntH* | Divalent metal cation transporter | transporter | A1189T | S397C | 1189 | 27-683 | MHI3538, |
| *moaC* | Cyclic pyranopterin monophosphate synthase | carbohydrate metabilism | A1T | M1L | 1 | 2-1191 | MHI4378, |
| *moaC* | Cyclic pyranopterin monophosphate synthase | carbohydrate metabilism | G448A | G150R | 448 | 10-57 | MHI3987, MHI4002, |
| *modA* | Molybdate-binding protein ModA | transporter | A13C | T5P | 13 | 20-946 | MHI4122, |
| *modA* | Molybdate-binding protein ModA | transporter | C14T | T5I | 14 | 20-946 | MHI4100, MHI4122, |
| *modA* | Molybdate-binding protein ModA | transporter | G14A | T5I | 14 | 73-1030 | MHI3710, MHI3711, |
| *modA* | Molybdate-binding protein ModA | transporter | C14T | T5I | 14 | 29-1030 | MHI3448, |
| *modA* | Molybdate-binding protein ModA | transporter | T19A | I7F | 19 | 1-266 | MHI4370, |
| *modA* | Molybdate-binding protein ModA | transporter | T19A | I7F | 19 | 10 | MHI4002, MHI4003, |
| *modA* | Molybdate-binding protein ModA | transporter | A19T | I7F | 19 | 20-946 | MHI4100, |
| *modA* | Molybdate-binding protein ModA | transporter | T19A | I7F | 19 | 16-396 | MHI4390, |
| *modA* | Molybdate-binding protein ModA | transporter | A19T | I7F | 19 | 61-1012 | MHI3668, |
| *modA* | Molybdate-binding protein ModA | transporter | G23A | S8L | 23 | 27-683 | MHI3535, |
| *modA* | Molybdate-binding protein ModA | transporter | A23G | S8L | 23 | 29-536 | MHI3559, MHI3562, MHI3570, |
| *modA* | Molybdate-binding protein ModA | transporter | T35C | L12S | 35 | 57-161 | MHI4259, |
| *modA* | Molybdate-binding protein ModA | transporter | T36A | L12F | 36 | 43-836 | MHI4406, |
| *modA* | Molybdate-binding protein ModA | transporter | A38C | I13S | 38 | 43-836 | MHI4406, |
| *modA* | Molybdate-binding protein ModA | transporter | G39C | I13M | 39 | 10-57 | MHI4002, MHI4003, |
| *modA* | Molybdate-binding protein ModA | transporter | C39G | I13M | 39 | 20-946 | MHI4100, |
| *modA* | Molybdate-binding protein ModA | transporter | G39C | I13M | 39 | 16-396 | MHI4390, |
| *modA* | Molybdate-binding protein ModA | transporter | C40T | A14T | 40 | 27-683 | MHI3535, |
| *modA* | Molybdate-binding protein ModA | transporter | G41T | A14E | 41 | 27-683 | MHI3535, |
| *modA* | Molybdate-binding protein ModA | transporter | G43A | G15R | 43 | 20-946 | MHI4100, |

| | | | | | | | |
|---|---|---|---|---|---|---|---|
| *modA* | Molybdate-binding protein ModA | transporter | C43T | G15R | 43 | 27-683 | MHI3535, |
| *mog* | Molybdopterin adenylyltransferase | environmental stress response | T1A | M1L | 1 | 64-2958 | MHI4331, |
| *mog* | Molybdopterin adenylyltransferase | environmental stress response | A1T | M1L | 1 | 27-683 | MHI3525, MHI3533, MHI3535, |
| *mog* | Molybdopterin adenylyltransferase | environmental stress response | A19T | I7F | 19 | 68-264 | MHI4354, |
| *mog* | Molybdopterin adenylyltransferase | environmental stress response | T20A | I7N | 20 | 68-264 | MHI4354, |
| *mpl* | UDP-N-acetylmuramate--L-alanyl-gamma-D-glutamyl-meso-2,6-diaminoheptandioate ligase | cell wall biosynthesis | T1357A | K453* | 1357 | 15-2957 | MHI3656, MHI3658, MHI3660, MHI3661, MHI3667, MHI3668, MHI3669, MHI4075, |
| *mpl* | UDP-N-acetylmuramate--L-alanyl-gamma-D-glutamyl-meso-2,6-diaminoheptandioate ligase | cell wall biosynthesis | T1357C | K453E | 1357 | 52-105 | MHI3649, |
| *mpl* | UDP-N-acetylmuramate--L-alanyl-gamma-D-glutamyl-meso-2,6-diaminoheptandioate ligase | cell wall biosynthesis | T1358G | K453T | 1358 | 52-105 | MHI3649, |
| *mpl* | UDP-N-acetylmuramate--L-alanyl-gamma-D-glutamyl-meso-2,6-diaminoheptandioate ligase | cell wall biosynthesis | T1359G | K453N | 1359 | 15-2957 | MHI4075, |
| *mpl* | UDP-N-acetylmuramate--L-alanyl-gamma-D-glutamyl-meso-2,6-diaminoheptandioate ligase | cell wall biosynthesis | A1360T | *454K | 1360 | 15-2957 | MHI4075, |
| *mpl* | UDP-N-acetylmuramate--L-alanyl-gamma-D-glutamyl-meso-2,6-diaminoheptandioate ligase | cell wall biosynthesis | T1360A | *454K | 1360 | 2-1191 | MHI3358, |
| *mpl* | UDP-N-acetylmuramate--L-alanyl-gamma-D-glutamyl-meso-2,6-diaminoheptandioate ligase | cell wall biosynthesis | A1360T | *454K | 1360 | 52-105 | MHI3644, MHI3649, |
| *mpl* | UDP-N-acetylmuramate--L-alanyl-gamma-D-glutamyl-meso-2,6-diaminoheptandioate ligase | cell wall biosynthesis | A1361C | *454S | 1361 | 2-1191 | MHI3358, |
| *mpl* | UDP-N-acetylmuramate--L-alanyl-gamma-D-glutamyl-meso-2,6-diaminoheptandioate ligase | cell wall biosynthesis | T1361G | *454S | 1361 | 52-105 | MHI3644, MHI3649, |
| *msbA 1* | ATP-dependent lipid A-core flippase | glycolipid metabolism | G35A | A12V | 35 | 75-544 | MHI3724, |
| *msbA 1* | ATP-dependent lipid A-core flippase | glycolipid metabolism | A115C | F39V | 115 | 57-161 | MHI4254, |
| *msbA 1* | ATP-dependent lipid A-core flippase | glycolipid metabolism | G119C | S40C | 119 | 57-161 | MHI4254, |
| *mtr* | Response regulator | regulator | C914T | T305I | 914 | 43-836 | MHI3452, |
| *murE* | UDP-N-acetylmuramoyl-L-alanyl-D-glutamate--2,6-diaminopimelate ligase | cell wall biosynthesis | T2C | M1T | 2 | 10-57 | MHI4012, MHI4013, |
| *murE* | UDP-N-acetylmuramoyl-L-alanyl-D-glutamate--2,6-diaminopimelate ligase | #N/A | A221C | E74A | 221 | 39-159 | MHI4162, MHI4171, MHI4184, MHI4191, |
| *murE* | UDP-N-acetylmuramoyl-L-alanyl-D-glutamate--2,6-diaminopimelate ligase | #N/A | C253T | H85Y | 253 | 39-159 | MHI4162, MHI4171, MHI4184, MHI4191, |
| *murE* | UDP-N-acetylmuramoyl-L-alanyl-D-glutamate--2,6-diaminopimelate ligase | #N/A | T383C | V128A | 383 | 39-159 | MHI4162, MHI4171, MHI4184, MHI4191, |
| *murE* | UDP-N-acetylmuramoyl-L-alanyl-D-glutamate--2,6-diaminopimelate ligase | #N/A | A503C | N168T | 503 | 39-159 | MHI4162, MHI4171, MHI4184, MHI4191, |
| *murE* | UDP-N-acetylmuramoyl-L-alanyl-D-glutamate--2,6-diaminopimelate ligase | #N/A | A513C | Q171H | 513 | 39-159 | MHI4162, MHI4171, MHI4184, MHI4191, |
| *murE* | UDP-N-acetylmuramoyl-L-alanyl-D-glutamate--2,6-diaminopimelate ligase | #N/A | G529A | A177T | 529 | 39-159 | MHI4162, MHI4171, MHI4184, MHI4191, |
| *murE* | UDP-N-acetylmuramoyl-L-alanyl-D-glutamate--2,6-diaminopimelate ligase | #N/A | T565C | Y189H | 565 | 39-159 | MHI4162, MHI4171, MHI4184, MHI4191, |
| *murE* | UDP-N-acetylmuramoyl-L-alanyl-D-glutamate--2,6-diaminopimelate ligase | #N/A | C572T | A191V | 572 | 39-159 | MHI4162, MHI4171, MHI4184, MHI4191, |
| *murE* | UDP-N-acetylmuramoyl-L-alanyl-D-glutamate--2,6-diaminopimelate ligase | #N/A | C593T | A198V | 593 | 39-159 | MHI4162, MHI4171, MHI4184, MHI4191, |
| *murF* | UDP-N-acetylmuramoyl-tripeptide--D-alanyl-D-alanine ligase | cell wall biosynthesis | A1195G | I399V | 1195 | 1-266 | MHI4370, |
| *murF* | UDP-N-acetylmuramoyl-tripeptide--D-alanyl-D-alanine ligase | cell wall biosynthesis | A1195G | I399V | 1195 | 2-544 | MHI3351, |
| *murF* | UDP-N-acetylmuramoyl-tripeptide--D-alanyl-D-alanine ligase | cell wall biosynthesis | A1195G | I399V | 1195 | 29-199 | MHI3558, |
| *murF* | UDP-N-acetylmuramoyl-tripeptide--D-alanyl-D-alanine ligase | cell wall biosynthesis | A1195G | I399V | 1195 | 52-105 | MHI3630, MHI3635, MHI3642, MHI3645, MHI3649, |
| *murF* | UDP-N-acetylmuramoyl-tripeptide--D-alanyl-D-alanine ligase | cell wall biosynthesis | T1195C | I399V | 1195 | 61-1012 | MHI3669, |
| *mutM* | Formamidopyrimidine-DNA glycosylase | transcription/translation | T438A | K146N | 438 | 43-836 | MHI3478, |
| *mutM* | Formamidopyrimidine-DNA glycosylase | transcription/translation | C443G | R148P | 443 | 43-836 | MHI3478, |
| *mutM* | Formamidopyrimidine-DNA glycosylase | transcription/translation | A277C | I93L | 277 | 29-536 | MHI3551, MHI3557, MHI3559, MHI3562, MHI3570, |
| *mutM* | Formamidopyrimidine-DNA glycosylase | transcription/translation | A277C | I93L | 277 | 59-280 | MHI4272, |
| *mutM* | Formamidopyrimidine-DNA glycosylase | transcription/translation | T277G | I93L | 277 | 61-568 | MHI3654, MHI3670, |
| *mutM* | Formamidopyrimidine-DNA glycosylase | transcription/translation | T289C | M97V | 289 | 52-105 | MHI3622, MHI3640, MHI3644, |
| *nagB* | Glucosamine-6-phosphate deaminase | carbohydrate metabilism | A13G | S5P | 13 | 61-568 | MHI3682, |
| *nagB* | Glucosamine-6-phosphate deaminase | carbohydrate metabilism | A35G | V12A | 35 | 52-105 | MHI3621, MHI3626, MHI3640, MHI3651, |
| *nagB* | Glucosamine-6-phosphate deaminase | carbohydrate metabilism | T67C | T23A | 67 | 61-568 | MHI3659, MHI3682, |
| *nagB* | Glucosamine-6-phosphate deaminase | carbohydrate metabilism | T274A | N92Y | 274 | 10-57 | MHI3988, |
| *nagB* | Glucosamine-6-phosphate deaminase | carbohydrate metabilism | G788A | A263V | 788 | 13-2156 | MHI4023, MHI4042, |
| *nagK* | N-acetyl-D-glucosamine kinase | cell wall biosynthesis | A53G | N18S | 53 | 8-431 | MHI3953, |
| *nanK* | N-acetylmannosamine kinase | transporter | T627G | K209N | 627 | 1-266 | MHI4369, |
| *nanK* | N-acetylmannosamine kinase | transporter | A627C | K209N | 627 | 64-2958 | MHI4321, |
| *nanK* | N-acetylmannosamine kinase | transporter | T627G | K209N | 627 | 59-1409 | MHI4296, |
| *nanK* | N-acetylmannosamine kinase | transporter | T627G | K209N | 627 | 68-264 | MHI4350, MHI4364, |
| *neuA* | N-acylneuraminate cytidylyltransferase | cell wall biosynthesis | C674T | W225* | 674 | 2-1191 | MHI3358, MHI3577, |
| *neuA* | N-acylneuraminate cytidylyltransferase | cell wall biosynthesis | C675T | W225* | 675 | 2-1191 | MHI3358, MHI3577, |
| *nhaA* | Na(+)/H(+) antiporter | transporter | T1202A | *401L | 1202 | 29-199 | MHI3558, |
| *norM* | Multidrug resistance protein | transporter | T16G | L6V | 16 | 56-344 | MHI4213, MHI4214, |
| *norM* | Multidrug resistance protein | transporter | T1163G | D388A | 1163 | 20-946 | MHI4091, |
| *ompP1* | Outer membrane protease | cell wall biosynthesis | A436G | N146D | 436 | 56-344 | MHI4195, MHI4198, MHI4210, MHI4212, MHI4213, MHI4214, MHI4215, MHI4217, MHI4219, MHI4221, MHI4222, MHI4224, |
| *opgE* | Phosphoethanolamine transferase | cell wall biosynthesis | T354A | S118R | 354 | 10-57 | MHI3978, |
| *opgE* | Phosphoethanolamine transferase | cell wall biosynthesis | T1124C | F375S | 1124 | 10-57 | MHI3989, |
| *oppA* | Periplasmic oligopeptide-binding protein | cell wall biosynthesis | G29A | R10Q | 29 | 61-1012 | MHI3658, |
| *patB* | Cystathionine beta-lyase PatB | iron metabolism | T4A | N2Y | 4 | 56-344 | MHI4213, |
| *patB* | Cystathionine beta-lyase PatB | iron metabolism | T23G | D8A | 23 | 43-836 | MHI3454, |
| *patB* | Cystathionine beta-lyase PatB | iron metabolism | G119C | P40R | 119 | 2954 | MHI4392, |
| *patB* | Cystathionine beta-lyase PatB | iron metabolism | A148C | F50V | 148 | 52-105 | MHI3649, |
| *patB* | Cystathionine beta-lyase PatB | iron metabolism | A457C | C153G | 457 | 43-836 | MHI3454, |
| *patB* | Cystathionine beta-lyase PatB | iron metabolism | A560G | K187R | 560 | 75-544 | MHI3723, |
| *patB* | Cystathionine beta-lyase PatB | iron metabolism | T773A | M258K | 773 | 15-1238 | MHI4074, |
| *patB* | Cystathionine beta-lyase PatB | iron metabolism | A831T | Y277* | 831 | 56-344 | MHI4214, |
| *patB* | Cystathionine beta-lyase PatB | iron metabolism | C991G | E331Q | 991 | 2954 | MHI4392, |
| *pcp* | Pyrrolidone-carboxylate peptidase | transcription/translation | G367C | E123Q | 367 | 59-1409 | MHI4281, |
| *pe* | Surface-adhesin protein E | cell wall biosynthesis | T479A | K160I | 479 | 1-266 | MHI4076, MHI4367, MHI4370, |
| *pe* | Surface-adhesin protein E | cell wall biosynthesis | A479C | K160T | 479 | 20-946 | MHI4091, MHI4104, MHI4114, |
| *pe* | Surface-adhesin protein E | cell wall biosynthesis | A479C | K160T | 479 | 36-36 | MHI4139, MHI4152, |
| *pe* | Surface-adhesin protein E | cell wall biosynthesis | T479A | K160I | 479 | 46-266 | MHI3490, MHI3494, |
| *pe* | Surface-adhesin protein E | cell wall biosynthesis | A479C | K160T | 479 | 57-161 | MHI4256, MHI4257, MHI4258, MHI4260, |
| *pe* | Surface-adhesin protein E | cell wall biosynthesis | A479C | K160T | 479 | 15-1238 | MHI4068, MHI4074, |
| *pe* | Surface-adhesin protein E | cell wall biosynthesis | T479A | K160I | 479 | 15-2957 | MHI4061, MHI4062, MHI4064, MHI4065, MHI4066, MHI4067, MHI4077, MHI4081, MHI4086, MHI4087, |
| *pe* | Surface-adhesin protein E | cell wall biosynthesis | A479C | K160T | 479 | 2-1040 | MHI3343, MHI3352, MHI3353, MHI3357, |
| *pe* | Surface-adhesin protein E | cell wall biosynthesis | T479A | K160I | 479 | 2-544 | MHI3354, |
| *pe* | Surface-adhesin protein E | cell wall biosynthesis | T479A | K160I | 479 | 29-199 | MHI3547, MHI3550, MHI3555, MHI3556, MHI3561, MHI3563, MHI3564, MHI3568, MHI3569, |
| *pe* | Surface-adhesin protein E | cell wall biosynthesis | T479A | K160I | 479 | 29-266 | MHI3429, MHI3440, MHI3441, MHI3443, MHI3444, |
| *pe* | Surface-adhesin protein E | cell wall biosynthesis | T479A | K160I | 479 | 52-105 | MHI3628, MHI3638, MHI3641, MHI3650, |
| *pe* | Surface-adhesin protein E | cell wall biosynthesis | T479A | K160I | 479 | 56-344 | MHI4198, MHI4211, |
| *pe* | Surface-adhesin protein E | cell wall biosynthesis | T479A | K160I | 479 | 59-1409 | MHI4267, MHI4271, |
| *pe* | Surface-adhesin protein E | cell wall biosynthesis | A479C | K160T | 479 | 61-1012 | MHI3671, |
| *pe* | Surface-adhesin protein E | cell wall biosynthesis | T479A | K160I | 479 | 61-568 | MHI3653, MHI3659, |
| *pe* | Surface-adhesin protein E | cell wall biosynthesis | T479A | K160I | 479 | 68-264 | MHI4345, MHI4353, MHI4354, MHI4358, |
| *pe* | Surface-adhesin protein E | cell wall biosynthesis | T479A | K160I | 479 | 8-431 | MHI3954, MHI3967, |
| *pe* | Surface-adhesin protein E | cell wall biosynthesis | A479C | K160T | 479 | 385545 | MHI3964, MHI3975, |
| *pe* | Surface-adhesin protein E | cell wall biosynthesis | T482A | *161L | 482 | 29-199 | MHI3544, |
| *pe* | Surface-adhesin protein E | cell wall biosynthesis | T482A | *161L | 482 | 8-431 | MHI3974, |
| *pepB* | Peptidase B | amino acid metabolism | A89C | H30P | 89 | 16-396 | MHI3611, |
| *pepB* | Peptidase B | amino acid metabolism | T92C | L31S | 92 | 10-57 | MHI3977, |
| *pepB* | Peptidase B | amino acid metabolism | T92C | L31S | 92 | 16-396 | MHI3611, |
| *pepB* | Peptidase B | amino acid metabolism | A93T | L31F | 93 | 10-57 | MHI3977, |
| *pepB* | Peptidase B | amino acid metabolism | A93T | L31F | 93 | 16-396 | MHI3611, |
| *pepB* | Peptidase B | amino acid metabolism | T104A | E35V | 104 | 75-544 | MHI3730, |
| *pepB* | Peptidase B | amino acid metabolism | A104T | E35V | 104 | 2-544 | MHI3354, |
| *pepB* | Peptidase B | amino acid metabolism | C110T | T37I | 110 | 46-266 | MHI3490, |
| *pepB* | Peptidase B | amino acid metabolism | G110A | T37I | 110 | 75-544 | MHI3717, MHI3730, |
| *pepB* | Peptidase B | amino acid metabolism | C110T | T37I | 110 | 16-396 | MHI3611, |
| *pepB* | Peptidase B | amino acid metabolism | C110T | T37I | 110 | 2-544 | MHI3354, |
| *pepB* | Peptidase B | amino acid metabolism | C110T | T37I | 110 | 29-266 | MHI3441, |
| *pepB* | Peptidase B | amino acid metabolism | G110A | T37I | 110 | 29-536 | MHI3551, |
| *pepB* | Peptidase B | amino acid metabolism | C110T | T37I | 110 | 52-105 | MHI3630, MHI3639, |
| *pepB* | Peptidase B | amino acid metabolism | C110T | T37I | 110 | 56-344 | MHI4213, |
| *pepB* | Peptidase B | amino acid metabolism | C110T | T37I | 110 | 68-264 | MHI4365, |
| *pepB* | Peptidase B | amino acid metabolism | A113C | D38A | 113 | 43-836 | MHI3454, MHI3466, MHI3467, MHI3476, MHI3478, MHI4406, |
| *pepB* | Peptidase B | amino acid metabolism | G124C | V42L | 124 | 52-105 | MHI3630, MHI3639, |
| *pepB* | Peptidase B | amino acid metabolism | C125T | T42I | 125 | 39-159 | MHI4170, MHI4172, |
| *pepB* | Peptidase B | amino acid metabolism | T125G | V42G | 125 | 52-105 | MHI3630, MHI3639, |
| *pepB* | Peptidase B | amino acid metabolism | A420T | N140K | 420 | 61-1012 | MHI3658, |

| | | | | | | | |
|---|---|---|---|---|---|---|---|
| *pepT* | Peptidase T | amino acid metabolism | G19A | K7E | 19 | 39-159 | MHI4167, MHI4168, MHI4169, MHI4171, MHI4172, MHI4173, MHI4184, MHI4191, MHI4193, |
| *pepT* | Peptidase T | amino acid metabolism | A19G | K7E | 19 | 16-396 | MHI4390, |
| *pepT* | Peptidase T | amino acid metabolism | A19G | K7E | 19 | 2-1191 | MHI3338, MHI3358, |
| *pepT* | Peptidase T | amino acid metabolism | T19C | K7E | 19 | 27-683 | MHI3515, MHI3519, MHI3527, MHI3535, MHI3541, |
| *phoR* | Phosphate regulon sensor protein PhoR | transporter | T20A | F7Y | 20 | 52-105 | MHI3648, |
| *phoR* | Phosphate regulon sensor protein PhoR | transporter | A37C | L13V | 37 | 20-946 | MHI4092, |
| *phoR* | Phosphate regulon sensor protein PhoR | transporter | C39G | L13F | 39 | 20-946 | MHI4092, |
| *phoR* | Phosphate regulon sensor protein PhoR | transporter | A279T | K93N | 279 | 75-544 | MHI3728, |
| *phoR* | Phosphate regulon sensor protein PhoR | transporter | A389C | V130G | 389 | 20-946 | MHI4092, |
| *phoR* | Phosphate regulon sensor protein PhoR | transporter | T389G | V130G | 389 | 8-2955 | MHI3966, |
| *phoR* | Phosphate regulon sensor protein PhoR | transporter | C393G | Q131H | 393 | 20-946 | MHI4092, |
| *phoR* | Phosphate regulon sensor protein PhoR | transporter | A408C | D136E | 408 | 20-946 | MHI4092, |
| *phoR* | Phosphate regulon sensor protein PhoR | transporter | G421C | E141Q | 421 | 75-544 | MHI3717, |
| *phoR* | Phosphate regulon sensor protein PhoR | transporter | A422C | E141A | 422 | 10-57 | MHI3990, |
| *phoR* | Phosphate regulon sensor protein PhoR | transporter | T429A | F143L | 429 | 15-1238 | MHI4074, |
| *phoR* | Phosphate regulon sensor protein PhoR | transporter | A437T | Y146F | 437 | 52-105 | MHI3649, |
| *phoR* | Phosphate regulon sensor protein PhoR | transporter | T439G | F147V | 439 | 29-1030 | MHI3432, |
| *phoR* | Phosphate regulon sensor protein PhoR | transporter | C444A | F148L | 444 | 52-105 | MHI3632, |
| *phoR* | Phosphate regulon sensor protein PhoR | transporter | T445A | S149C | 445 | 39-159 | MHI4162, |
| *phoR* | Phosphate regulon sensor protein PhoR | transporter | A445C | S149R | 445 | 43-836 | MHI3478, |
| *phoR* | Phosphate regulon sensor protein PhoR | transporter | T447A | S149R | 447 | 43-836 | MHI3478, |
| *phoR* | Phosphate regulon sensor protein PhoR | transporter | C448A | Q150K | 448 | 43-836 | MHI3478, |
| *phoR* | Phosphate regulon sensor protein PhoR | transporter | G458C | R153T | 458 | 1-266 | MHI4368, |
| *phoR* | Phosphate regulon sensor protein PhoR | transporter | G458C | R153T | 458 | 57-161 | MHI4257, |
| *phoR* | Phosphate regulon sensor protein PhoR | transporter | C458G | R153T | 458 | 56-344 | MHI4216, |
| *phoR* | Phosphate regulon sensor protein PhoR | transporter | T854A | N285I | 854 | 29-536 | MHI3551, |
| *pldB* | Probable lysophospholipase | glycolipid metabolism | A793C | N265H | 793 | 20-946 | MHI4092, |
| *pldB* | Probable lysophospholipase | glycolipid metabolism | G793A | Q265* | 793 | 29-536 | MHI3551, MHI3557, MHI3559, MHI3562, MHI3570, |
| *pldB* | Probable lysophospholipase | glycolipid metabolism | C795A | N265K | 795 | 20-946 | MHI4092, |
| *pldB* | Probable lysophospholipase | glycolipid metabolism | G795T | N265K | 795 | 73-1030 | MHI3712, MHI3712, |
| *pldB* | Probable lysophospholipase | glycolipid metabolism | C795A | N265K | 795 | 15-1238 | MHI4068, |
| *pldB* | Probable lysophospholipase | glycolipid metabolism | C795A | N265K | 795 | 61-1012 | MHI3661, MHI3669, |
| *pldB* | Probable lysophospholipase | glycolipid metabolism | T796G | K266Q | 796 | 29-536 | MHI3551, MHI3557, MHI3559, MHI3562, MHI3570, |
| *pldB* | Probable lysophospholipase | glycolipid metabolism | T801A | N267K | 801 | 20-946 | MHI4092, |
| *pldB* | Probable lysophospholipase | glycolipid metabolism | A801T | N267K | 801 | 73-1030 | MHI3712, |
| *pldB* | Probable lysophospholipase | glycolipid metabolism | T801A | N267K | 801 | 15-1238 | MHI4068, MHI4074, |
| *pldB* | Probable lysophospholipase | glycolipid metabolism | T801A | N267K | 801 | 61-1012 | MHI3658, MHI3661, MHI3669, |
| *plsY* | Probable glycerol-3-phosphate acyltransferase | glycolipid metabolism | A589T | K197* | 589 | 52-105 | MHI3636, |
| *plsY* | Probable glycerol-3-phosphate acyltransferase | glycolipid metabolism | G594T | K198N | 594 | 29-199 | MHI3558, |
| *plsY* | Probable glycerol-3-phosphate acyltransferase | glycolipid metabolism | G594T | K198N | 594 | 52-105 | MHI3636, MHI3644, |
| *plsY* | Probable glycerol-3-phosphate acyltransferase | glycolipid metabolism | A595C | K199Q | 595 | 29-199 | MHI3558, |
| *plsY* | Probable glycerol-3-phosphate acyltransferase | glycolipid metabolism | A595T | K199* | 595 | 52-105 | MHI3636, MHI3644, |
| *plsY* | Probable glycerol-3-phosphate acyltransferase | glycolipid metabolism | A595C | K199Q | 595 | 52-105 | MHI3636, MHI3644, |
| *plsY* | Probable glycerol-3-phosphate acyltransferase | glycolipid metabolism | A597C | K199N | 597 | 29-199 | MHI3558, |
| *plsY* | Probable glycerol-3-phosphate acyltransferase | glycolipid metabolism | A597T | K199N | 597 | 52-105 | MHI3636, MHI3643, MHI3644, |
| *plsY* | Probable glycerol-3-phosphate acyltransferase | glycolipid metabolism | A597C | K199N | 597 | 52-105 | MHI3636, MHI3643, MHI3644, |
| *potB* | Spermidine/putrescine transport system permease | transporter | C301A | L101I | 301 | 10-57 | MHI3982, |
| *purR* | HTH-type transcriptional repressor PurR | regulator | A1009T | *337K | 1009 | 1-266 | MHI3574, |
| *sixA* | Phosphohistidine phosphatase | transcription/translation | G46A | D16N | 46 | 57-161 | MHI4259, |
| *sixA* | Phosphohistidine phosphatase | transcription/translation | G473T | S158Y | 473 | 39-159 | MHI4169, |
| *slp* | Outer membrane protein | cell wall biosynthesis | C21A | F7L | 21 | 29-536 | MHI3551, MHI3557, MHI3559, MHI3562, MHI3570, |
| *slp* | Outer membrane protein | cell wall biosynthesis | C21A | F7L | 21 | 29-598 | MHI3545, |
| *sotB* | Sugar efflux transporter | transporter | C1183A | Q395K | 1183 | 43-836 | MHI3466, MHI3478, |
| *sotB* | Sugar efflux transporter | transporter | T1187G | K396T | 1187 | 61-1012 | MHI3658, MHI3661, |
| *sotB* | Sugar efflux transporter | transporter | A1190C | K397T | 1190 | 43-836 | MHI3456, |
| *sotB* | Sugar efflux transporter | transporter | A1190C | K397T | 1190 | 15-2957 | MHI4067, |
| *ssb_1* | Single-stranded DNA-binding protein | transcription/translation | C128T | R43K | 128 | 52-105 | MHI3627, MHI3631, MHI3651, |
| *ssb_1* | Single-stranded DNA-binding protein | transcription/translation | T263A | Q88L | 263 | 52-105 | MHI3627, MHI3631, MHI3651, |
| *ssb_1* | Single-stranded DNA-binding protein | transcription/translation | T338C | N113S | 338 | 52-105 | MHI3627, MHI3631, MHI3651, |
| *ssb_1* | Single-stranded DNA-binding protein | transcription/translation | G359C | T120S | 359 | 52-105 | MHI3627, MHI3631, MHI3651, |
| *ssb_1* | Single-stranded DNA-binding protein | transcription/translation | T389C | N130S | 389 | 52-105 | MHI3627, MHI3631, MHI3651, |
| *ssb_1* | Single-stranded DNA-binding protein | transcription/translation | C424T | V142I | 424 | 52-105 | MHI3627, MHI3631, MHI3651, |
| *tamB* | Translocation and assembly module subunit TamB | transporter | A527T | N176I | 527 | 385545 | MHI3975, |
| *tamB* | Translocation and assembly module subunit TamB | transporter | T549A | F183L | 549 | 61-1012 | MHI3657, MHI3658, MHI3660, MHI3661, MHI3662, MHI3663, MHI3669, MHI3671, MHI3676, MHI3677, MHI3678, MHI3679, MHI3680, MHI3681, MHI3683, |
| *tamB* | Translocation and assembly module subunit TamB | transporter | T549A | L183F | 549 | 8-431 | MHI3952, MHI3953, |
| *tamB* | Translocation and assembly module subunit TamB | transporter | T549A | F183L | 549 | 8-2955 | MHI3964, MHI3965, |
| *tamB* | Translocation and assembly module subunit TamB | transporter | T551C | N184S | 551 | 8-431 | MHI3952, |
| *tamB* | Translocation and assembly module subunit TamB | transporter | A551G | N184S | 551 | 385545 | MHI3975, |
| *tamB* | Translocation and assembly module subunit TamB | transporter | A552C | N184K | 552 | 8-431 | MHI3952, |
| *tamB* | Translocation and assembly module subunit TamB | transporter | T552G | N184K | 552 | 385545 | MHI3975, |
| *tamB* | Translocation and assembly module subunit TamB | transporter | T554C | N185S | 554 | 8-431 | MHI3952, |
| *tamB* | Translocation and assembly module subunit TamB | transporter | A554G | N185S | 554 | 385545 | MHI3975, |
| *tbpB* | Transferrin-binding protein 2 | iron metabolism | A315C | N105K | 315 | 2-544 | MHI3337, |
| *tbpB* | Transferrin-binding protein 2 | iron metabolism | T323C | K108R | 323 | 2-544 | MHI3337, |
| *tbpB* | Transferrin-binding protein 2 | iron metabolism | C328T | G110R | 328 | 2-544 | MHI3337, |
| *tbpB* | Transferrin-binding protein 2 | iron metabolism | C329T | G110E | 329 | 2-544 | MHI3337, |
| *tbpB* | Transferrin-binding protein 2 | iron metabolism | G338T | A113D | 338 | 2-544 | MHI3334, MHI3337, |
| *tbpB* | Transferrin-binding protein 2 | iron metabolism | G340T | L114I | 340 | 2-544 | MHI3337, MHI3348, |
| *tbpB* | Transferrin-binding protein 2 | iron metabolism | T427A | I143F | 427 | 27-494 | MHI3417, |
| *tbpB* | Transferrin-binding protein 2 | iron metabolism | A1702T | T568S | 1702 | 36-36 | MHI4152, |
| *tbpB* | Transferrin-binding protein 2 | iron metabolism | C1703T | T568I | 1703 | 36-36 | MHI4152, |
| *tbpB* | Transferrin-binding protein 2 | iron metabolism | C1705G | Q569E | 1705 | 36-36 | MHI4152, |
| *tdk* | Thymidine kinase | transcription/translation | G574C | Q192E | 574 | 1-266 | MHI3574, MHI4370, |
| *tdk* | Thymidine kinase | transcription/translation | G574C | Q192E | 574 | 46-266 | MHI3489, MHI3494, |
| *tdk* | Thymidine kinase | transcription/translation | C574G | Q192E | 574 | 15-1238 | MHI4074, |
| *tdk* | Thymidine kinase | transcription/translation | G574C | Q192E | 574 | 16-396 | MHI4390, |
| *tdk* | Thymidine kinase | transcription/translation | G574C | Q192E | 574 | 27-683 | MHI3525, MHI3532, MHI3533, MHI3535, |
| *tdk* | Thymidine kinase | transcription/translation | G574C | Q192E | 574 | 29-266 | MHI3443, |
| *tdk* | Thymidine kinase | transcription/translation | T577A | I193F | 577 | 1-266 | MHI3574, MHI4370, |
| *tdk* | Thymidine kinase | transcription/translation | T577A | I193F | 577 | 46-266 | MHI3489, MHI3494, |
| *tdk* | Thymidine kinase | transcription/translation | T577A | I193F | 577 | 29-266 | MHI3443, |
| *tdk* | Thymidine kinase | transcription/translation | T578A | I193N | 578 | 57-161 | MHI4255, |
| *tdk* | Thymidine kinase | transcription/translation | A581C | *194S | 581 | 57-161 | MHI4255, |
| *tdk* | Thymidine kinase | transcription/translation | A583C | I195L | 583 | 57-161 | MHI4255, |
| *trpC* | Tryptophan biosynthesis protein | transcription/translation | A1391G | D464G | 1391 | 27-683 | MHI3518, |
| *trpC* | Tryptophan biosynthesis protein | transcription/translation | C1393T | Q465* | 1393 | 27-683 | MHI3515, MHI3518, |
| *trpC* | Tryptophan biosynthesis protein | transcription/translation | A1395T | Q465H | 1395 | 27-683 | MHI3515, MHI3518, |
| *trpC* | Tryptophan biosynthesis protein | transcription/translation | G1396T | E466* | 1396 | 27-683 | MHI3515, MHI3518, |
| *trpC* | Tryptophan biosynthesis protein | transcription/translation | A1411C | F471V | 1411 | 61-1012 | MHI3657, MHI3660, MHI3662, MHI3663, MHI3667, MHI3671, MHI3676, MHI3677, MHI3678, MHI3679, MHI3680, MHI3681, MHI3683, |
| *trpC* | Tryptophan biosynthesis protein | transcription/translation | C1416A | L472F | 1416 | 2-1191 | MHI3577, MHI4378, |
| *trpC* | Tryptophan biosynthesis protein | transcription/translation | T1416G | F472L | 1416 | 27-683 | MHI3515, MHI3518, MHI3527, MHI3529, |
| *trpC* | Tryptophan biosynthesis protein | transcription/translation | A1420T | N474Y | 1420 | 27-683 | MHI3518, |
| *trpE* | Anthranilate synthase component 1 | transcription/translation | T257C | Y86C | 257 | 61-1012 | MHI3658, |
| *trpE* | Anthranilate synthase component 1 | transcription/translation | T280A | K94* | 280 | 61-1012 | MHI3658, MHI3668, |
| *tuaG* | Teichuronic acid biosynthesis glycosyltransferase | cell wall biosynthesis | A454T | K152* | 454 | 73-1030 | MHI3713, |
| *tuaG* | Teichuronic acid biosynthesis glycosyltransferase | cell wall biosynthesis | T461A | I154K | 461 | 10-57 | MHI3977, |
| *tusB* | Protein TusB | transcription/translation | T286C | *96Q | 286 | 73-1030 | MHI3694, |
| *tusB* | Protein TusB | transcription/translation | T286C | *96Q | 286 | 8-431 | MHI3969, |
| *tusB* | Protein TusB | transcription/translation | A287C | *96S | 287 | 73-1030 | MHI3561, MHI3690, MHI3694, |
| *tusB* | Protein TusB | transcription/translation | A287C | *96S | 287 | 15-2957 | MHI4063, MHI4069, MHI4071, |

| | | | | | | | |
|---|---|---|---|---|---|---|---|
| *tusB* | Protein TusB | transcription/translation | A287C | *96S | 287 | 29-266 | MHI3435, MHI3447, MHI3450, |
| *tusB* | Protein TusB | transcription/translation | A287C | *96S | 287 | 8-431 | MHI3969, |
| *tusC* | Protein TusC | transcription/translation | G293C | C98S | 293 | 29-199 | MHI3564, |
| *tusC* | Protein TusC | transcription/translation | T310A | S104T | 310 | 1-266 | MHI3574, |
| *tusC* | Protein TusC | transcription/translation | A310T | S104T | 310 | 20-946 | MHI4092, |
| *tusC* | Protein TusC | transcription/translation | T310A | S104T | 310 | 39-159 | MHI4172, |
| *tusC* | Protein TusC | transcription/translation | T310A | S104T | 310 | 73-1030 | MHI3711, |
| *tusC* | Protein TusC | transcription/translation | T310A | S104T | 310 | 15-2957 | MHI4067, MHI4086, MHI4087, |
| *tusC* | Protein TusC | transcription/translation | T310A | S104T | 310 | 29-199 | MHI3564, |
| *tusC* | Protein TusC | transcription/translation | T310A | S104T | 310 | 52-105 | MHI3646, |
| *tusC* | Protein TusC | transcription/translation | G322A | E108K | 322 | 1-266 | MHI3574, |
| *tusC* | Protein TusC | transcription/translation | G322A | E108K | 322 | 73-1030 | MHI3714, |
| *tusC* | Protein TusC | transcription/translation | A325T | K109* | 325 | 1-266 | MHI3574, |
| *tusC* | Protein TusC | transcription/translation | A325T | K109* | 325 | 73-1030 | MHI3714, |
| *tusC* | Protein TusC | transcription/translation | A326T | K109M | 326 | 1-266 | MHI3574, |
| *tusC* | Protein TusC | transcription/translation | A326T | K109M | 326 | 73-1030 | MHI3714, |
| *tusE* | Protein TusE | transcription/translation | C325G | L109V | 325 | 46-266 | MHI3494, |
| *tusE* | Protein TusE | transcription/translation | T326A | L109Q | 326 | 10-57 | MHI3991, MHI4003, MHI4013, |
| *tusE* | Protein TusE | transcription/translation | A326T | L109Q | 326 | 8-431 | MHI3952, MHI3953, |
| *tusE* | Protein TusE | transcription/translation | T328A | *110K | 328 | 46-266 | MHI3494, |
| *tyrP_1* | Tyrosine-specific transport protein | transporter | T1201A | *401K | 1201 | 1-266 | MHI3574, |
| *tyrP_1* | Tyrosine-specific transport protein | transporter | T1201A | *401K | 1201 | 7-159 | MHI4382, |
| *tyrP_1* | Tyrosine-specific transport protein | transporter | T1201G | *401E | 1201 | 73-1030 | MHI3710, |
| *tyrP_1* | Tyrosine-specific transport protein | transporter | T1201G | *401E | 1201 | 2-544 | MHI3354, |
| *tyrP_1* | Tyrosine-specific transport protein | transporter | T1201G | *401E | 1201 | 59-1409 | MHI4269, MHI4283, |
| *tyrP_1* | Tyrosine-specific transport protein | transporter | A1202T | *401L | 1202 | 1-266 | MHI3574, |
| *tyrP_1* | Tyrosine-specific transport protein | transporter | A1202T | *401L | 1202 | 7-159 | MHI4382, |
| *ulaE* | L-ribulose-5-phosphate 3-epimerase | carbohydrate metabilism | C10T | H4Y | 10 | 75-544 | MHI3737, |
| *ulaE* | L-ribulose-5-phosphate 3-epimerase | carbohydrate metabilism | C10T | H4Y | 10 | 8-2955 | MHI3975, |
| *ulaE* | L-ribulose-5-phosphate 3-epimerase | carbohydrate metabilism | A11T | H4L | 11 | 75-544 | MHI3737, |
| *ulaE* | L-ribulose-5-phosphate 3-epimerase | carbohydrate metabilism | A11T | H4L | 11 | 8-2955 | MHI3975, |
| *ulaE* | L-ribulose-5-phosphate 3-epimerase | carbohydrate metabilism | A13G | K5E | 13 | 75-544 | MHI3737, |
| *ulaE* | L-ribulose-5-phosphate 3-epimerase | carbohydrate metabilism | A13G | K5E | 13 | 8-2955 | MHI3975, |
| *ulaE* | L-ribulose-5-phosphate 3-epimerase | carbohydrate metabilism | A14C | K5T | 14 | 8-2955 | MHI3975, |
| *ung* | Uracil-DNA glycosylase | transcription/translation | T8G | N3T | 8 | 20-946 | MHI4124, |
| *yejM* | Inner membrane protein | transporter | A1135G | N379D | 1135 | 52-105 | MHI3645, |
| *yejM* | Inner membrane protein | transporter | A1138G | N380D | 1138 | 52-105 | MHI3644, MHI4265, |
| *yejM* | Inner membrane protein | transporter | T1160G | L387R | 1160 | 52-105 | MHI3645, MHI3646, MHI3648, |
| *yejM* | Inner membrane protein | transporter | T1160A | L387H | 1160 | 52-105 | MHI3645, MHI3646, MHI3648, |
| *yejM* | Inner membrane protein | transporter | A1162C | N388H | 1162 | 52-105 | MHI3645, MHI3648, |
| *yejM* | Inner membrane protein | transporter | A1163T | N388I | 1163 | 52-105 | MHI3646, |
| *yejM* | Inner membrane protein | transporter | A1166T | D389V | 1166 | 52-105 | MHI3645, |
| *yejM* | Inner membrane protein | transporter | T1170A | F390L | 1170 | 52-105 | MHI3645, |
| *ytfE* | Iron-sulfur cluster repair protein | iron metabolism | T673A | F225I | 673 | 46-266 | MHI3490, |
| *znuA* | High-affinity zinc uptake system protein | transporter | G47C | S16T | 47 | 52-105 | MHI3624, MHI3630, MHI3632, MHI3644, MHI3649, |

**Table S5. Presence of each polymorphic gene in the 39 Child-MLST matched reference genoems.**

| Gene | MHI3301 | MHI3331 | MHI3332 | MHI3338 | MHI3341 | MHI3361 | MHI3391 | MHI3401 | MHI3421 | MHI3428 | MHI3451 | MHI3481 | MHI3511 | MHI3512 | MHI3542 | MHI3543 | MHI3546 | MHI3578 | MHI3620 | MHI3652 | MHI3655 | MHI3685 | MHI3716 | MHI3941 | MHI3963 | MHI3976 | MHI4015 | MHI4050 | MHI4051 | MHI4060 | MHI4090 | MHI4125 | MHI4160 | MHI4194 | MHI4227 | MHI4262 | MHI4264 | MHI4297 | MHI4332 |
|---|---|---|---|---|---|---|---|---|---|---|---|---|---|---|---|---|---|---|---|---|---|---|---|---|---|---|---|---|---|---|---|---|---|---|---|---|---|---|---|
| *adk* | present | present | present | present | present | present | present | present | present | present | present | present | present | present | present | present | present | present | present | present | present | present | present | present | present | present | present | present | present | present | present | present | present | present | present | present | present | present | present |
| *argG* | present | present | present | present | present | present | present | present | present | present | present | present | present | present | present | present | present | present | present | present | present | present | present | present | present | present | present | present | present | present | present | present | present | present | present | present | present | present | present |
| *aroA* | present | present | present | present | present | present | present | present | present | present | present | present | present | present | present | present | present | present | present | present | present | present | present | present | present | present | present | present | present | present | present | present | present | present | present | present | present | present | present |
| *artP* | present | present | present | present | present | present | present | present | present | present | present | present | present | present | present | present | present | present | present | present | present | present | present | present | present | present | present | present | present | present | present | present | present | present | present | present | present | present | present |
| *aspA* | present | present | present | present | present | present | present | present | present | present | present | present | present | present | present | present | present | present | present | present | present | present | present | present | present | present | present | present | present | present | present | present | present | present | present | present | present | present | present |
| *atsB* | absent | absent | absent | absent | absent | absent | present | present | absent | absent | present | absent | present | absent | absent | absent | absent | absent | absent | absent | absent | absent | absent | absent | absent | absent | present | absent | absent | absent | absent | absent | absent | absent | absent | absent | absent | absent | absent |
| *bacA* | present | present | present | present | present | present* | present | present | present | present* | present | present | present | present* | present | present* | present | present* | present | present | present* | present* | present | present | present | present* | present* | present | present | present* | present | present | present | present* | present* | present | present | present | present |
| *bamA* | present | present | present | present | present | present | present | present | present | present | present | present | present | present | present | present | present | present | present | present | present | present | present | present | present | present | present | present | present | present | present | present | present | present | present | present | present | present | present |
| *bcp* | present | present | present | present | present | present | present | present | present | present | present | present | present | present | present | present | present | present | present | present | present | present | present | present | present | present | present | present | present | present | present | present | present | present | present | present | present | present | present |
| *bioD1* | present | present | present | present | present | present | present | present | present | present | present | present | present | present | present | present | present | present | present | present | present | present | present | present | present | present | present | present | present | present | present | present | present | present | present | present | present | present | present |
| *bolA* | present | present | present | present | present | present | present | present | present | present | present | present | present | present | present | present | present | present | present | present | present | present | present | present | present | present | present | present | present | present | present | present | present | present | present | present | present | present | present |
| *bspRIM* | absent | absent | absent | present | present | absent | absent | absent | absent | absent | absent | absent | present | present | present | absent | absent | present | absent | absent | present | absent | absent | absent | absent | absent | absent | absent | present | absent | absent | absent | present | present | absent | absent | absent | absent | absent |
| *can* | present | present | present | present | present | present | present | present | present | present | present | present | present | present | present | present | present | present | present | present | present | present | present | present | present | present | present | present | present | present | present | present | present | present | present | present | present | present | present |
| *cbiM* | present | present | present | present | present | present | present | present | present | present | present | present | present | present | present | present | present | present | present | present | present | present | present | present | present | present | present | present | present | present | present | present | present | present | present | present | present | present | present |
| *ccmC* | present | present | present | present | present | present | present | present | present | present | present | present | present | present | present | present | present | present | present | present | present | present | present | present | present | present | present | present | present | present | present | present | present | present | present | present | present | present | present |
| *chuR_1*** | present | present | present | present | present | present | present | present | present | present | present | present | present | present | present | present | present | present | present | present | present | present | present | present | present | present | present | present | present | present | present | present | present | present | present | present | present | present | present |
| *chuR_2*** | present | absent | absent | absent | absent | absent | absent | absent | present | absent | absent | present | absent | absent | absent | absent | absent | absent | absent | absent | absent | absent | absent | absent | absent | absent | absent | absent | absent | absent | present | absent | absent | absent | absent | absent | absent | absent | absent |
| *citC* | present | present | present | present | present | present | present | present | present | present | present | present | present | present | present | present | present | present | present | present | present | present | present | present | present | absent | present | absent | present | present | present | absent | present | present | present | present | present | present | present |
| *citG* | present | present | present | present | present | present | present | present | present | present | present | present | present | present | present | present | present | present | present | present | present | present | present | present | present | absent | present | absent | present | present | present | absent | present | present | present | present | present | present | present |
| *cntO* | absent | absent | absent | absent | absent | present | absent | absent | absent | present | absent | absent | absent | present | absent | present | absent | present | absent | absent | present | present | absent | absent | absent | present | present | absent | absent | absent | absent | absent | present | present | present | absent | absent | absent | absent |
| *coaBC* | present | present | present | present | present | present | present | present | present | present | present | present | present | present | present | present | present | present | present | present | present | present | present | present | present | present | present | present | present | present | present | present | present | present | present | present | present | present | present |
| *copA* | present | present | present | present | present | present | present | present | present | present | present | present | present | present | present | present | present | present | present | present | present | present | present | present | present | present | present | present | present | present | present | present | present | present | present | present | present | present | present |
| *cpxR* | present | present | present | present | present | present | present | present | present | present | present | present | present | present | present | present | present | present | present | present | present | present | present | present | present | present | present | present | present | present | present | present | present | present | present | present | present | present | present |
| *csdA* | present | present | present | present | present | present | present | present | present | present | present | present | present | present | present | present | present | present | present | present | present | present | present | present | present | present | present | present | present | present | absent | absent | absent | present | present | present | present | present | present |
| *csdE* | present | present | present | present | present | present | present | present | present | present | present | present | present | present | present | present | present | present | present | present | present | present | present | present | present | present | present | present | present | present | present | present | present | present | present | present | present | present | present |
| *cydD* | present* | present* | present | present* | present | present* | present | present | present* | present | present | present* | present | present* | present* | present* | present | present* | present | present* | present | present | present | present | present | present* | present | present | present* | present* | present* | present | present* | present* | present | present | present* | present* | present* |
| *dacA* | present | present | present | present | present | present | present | present | present | present | present | present | present | present | present | present | present | present | present | present | present | present | present | present | present | present | present | present | present | present | present | present | present | present | present | present | present | present | present |
| *dam* | present | present | present | present | present | present | present | present | present | present | present | present | present | present | present | present | present | present | present | present | present | present | present | present | present | present | present | present | present | present | present | present | present | present | present | present | present | present | present |
| *dapD* | present | present | present | present | present | present | present | present | present | present | present | present | present | present | present | present | present | present | present | present | present | present | present | present | present | present | present | present | present | present | present | present | present | present | present | present | present | present | present |
| *dcuD* | present | present | present | present | present | present | present | present | present | present | present | present | present | present | present | present | present | present | present | present | present | present | present | present | present | present | present | present | present | present | present | present | present | present | present | present | present | present | present |
| *dmsD* | present | present | present | present | present | present | present | present | present | present | present | present | present | present | present | present | present | present | present | present | present | present | present | present | present | present | present | present | present | present | present | present | present | present | present | present | present | present | present |
| *dtd* | present | present | present | present | present | present | present | present | present | present | absent | present | present | present | present | present | present | present | present | present | present | present | present | present | present | present | present | present | present | present | present | present | present | present | present | absent | present | present | present |
| *envC* | present | present | present | present | present | present | present | present | present | present | present | present | present | present | present | present | present | present | present | present | present | present | present | present | present | present | present | present | present | present | present | present | present | present | present | present | present | present | present |
| *era* | present | present | present | present | present | present | present | present | present | present | present | present | present | present | present | present | present | present | present | present | present | present | present | present | present | present | present | present | present | present | present | present | present | present | present | present | present | present | present |
| *esiB* | present | present | present | present | present | present | present | present | present | present | present | present | present | present | present | present | present | present | present | present | present | present | present | present | present | present | present | present | present | present | present* | present | present | present* | present* | present | present | present | present |
| *fbp* | present | present | present | present | present* | present | present | present | present | present* | present | present | present | present | present | present | present | present | present* | present* | present | present* | present | present | present | present | present | present | present | present | present | present | present | present | present | present | present* | present | present |
| *fbpC2* | present | present | present | present | present | present | present | present | present | present | present | present | present | present | present | present | present | present | present | present | present | present | present | absent | present | present | absent | present | present | present | present | present | present | present | present | present | present | present | present |
| *fhuD* | absent | absent | absent | present | present | present | absent | absent | absent | present | absent | absent | absent | present | present | absent | absent | present | present | present | present | present | absent | absent | absent | absent | present | absent | present | absent | absent | absent | present | present | present | absent | absent | absent | absent |
| *fnr* | present | present | present | present | present | present | present | present | present | present | present | present | present | present | present | present | present | present | present | present | present | present | present | present | present | present* | present | present | present | present | present | present | present | present | present | present | present | present | present |
| *ftsH_1*** | present | present | present | present | present | present | present | present | present | present | present | present | present | present | present | present | present | present | present | present | present | present | present | present | present | present | present | present | present | present | present | present | present | present | present | present | present | present | present |
| *ftsH_2*** | present | present | present | present | absent | present | present | present | present | absent | absent | present | present | absent | absent | absent | present | present | absent | absent | absent | absent | present | present | present | present | present | absent | absent | present | present | absent | present | present | present | present | absent | present | present |
| *ftsI* | present | present | present | present | present | present | present | present | present | present | present | present | present | present | present | present | present | present | present | present | present | present | present | present | present | present | present | present | present | present | present | present | present | present | present | present | present | present | present |
| *ftsN* | present | present | present | present | present | present | present | present | present | present | present | present | present | present | present | present | present | present | present | present | present | present | present | present | present | present | present | present | present | present | present | present | present | present | present | present | present | present | present |
| *glmM* | present | present | present | present | present | present* | present | present | present | present | present | present | present | present | present | present | present | present* | present | present | present | present | present | present | present | present | present* | present | present | present | present | present | present* | present | present* | present | present | present | present |
| *glnE* | present | present | present | present | present | present | present | present | present | present | present | present | present | present | present | present | present | present | present | present | present | present | present | present | present | present | present | present | present | present | present | present | present | present | present | present | present | present | present |
| *glpQ* | present | present | present | present | present | present | present | present | present | present | absent | present | present | present | present | present | present | present | present | present | present | present | present | present | present | present | present | present | present | present | present | absent | present | present | present | absent | absent | present | present |
| *gltC* | present | present | present | present | present | present | present | present | present | present | absent | present | present | present | present | present | present | present | present | present | present | present | present | present | present | present | present | present | present | present | present | present | present | present | present | absent | present | present | present |
| *gmhA_2* | present | present | present | present | present | present | present | present | present | present | present | present | present | present | present | present | present | present | present | present | present | present | present | present | present | present | present | present | present | present | present | present | present | present | present | present | present | present | present |
| *gor* | present | present | present | present | present | present | present | present | present | present | present | present | present | present | present | present | present | present | present | present | present | present | present | present | present | present | present | present | present | present | present | present | present | present | present | present | present | present | present |
| *gpt_1* | present | present | present | present | present | present | present | present | present | present | absent | present | present | present | present | present | present | present | present | present | present | present | present | present | present | absent | present | absent | present | present | present | absent | present | present | present | absent | absent | present | present |
| *gpt_2* | present | present | present | present | present | present | present | present | present | present | absent | present | present | present | present | present | present | present | present | present | present | present | present | present | present | absent | present | absent | present | present | present | absent | present | present | present | absent | absent | present | present |
| *greA* | present | present | present | present | present | present | present | present | present | present | present | present | present | present | present | present | present | present | present | present | present | present | present | present | present | present | present | present | present | present | present | present | present | present | present | present | present | present | present |
| *hap* | present | present | present | present | present | present | present | present | present | present | present | present | present | present | present | present | present | present | present | present | present | present | present | present | present | present | present | absent | present | present | present | present | present | present | present | present | present | present | present |
| *hindIIIM* | present | absent | absent | absent | absent | present | absent | absent | present | absent | absent | present | absent | absent | absent | absent | absent | present | absent | absent | absent | absent | absent | absent | absent | absent | present | absent | absent | absent | present | absent | present | absent | present | absent | absent | absent | absent |
| *hindIIIR* | present | absent | absent | absent | absent | present | absent | absent | present | absent | absent | present | absent | absent | absent | absent | absent | present | absent | absent | absent | absent | absent | absent | absent | absent | present | absent | absent | absent | present | absent | present | absent | present | absent | absent | absent | absent |
| *hsdR* | present | present | present | present | present | present | present | present | present | present | present | present | present | present | present | present | present | present | present | present | present | present | present | present | present | present | present | absent | present | present | present | present | present | present | present | present | present | present | present |
| *hxuC* | present | present | present | present | present | present | present | present | present | present | present | present | present | present | present | present | present | present | present | present | present | present | present | present | present | present | present | present | present | present | present | present | present | present | present | present | present | present | present |
| *iaaH* | present | present | present | absent | present | present | present | present | present | present | present | present | present | present | present | absent | present | present | present | present | present | present | present | present | present | present | absent | absent | present | present | present | absent | present | present | present | present | present | present | present |
| *iga* | present | present | present | present | present | present | present | present | present | present | present | present | absent | present | present | present | present | present | present | present | present | present | present | present | present | present | present | absent | present | present | present | present | present | present | present | present | present | present | present |
| *ihfB* | present | present | present | present | present | present | present | present | present | present | present | present | present | present | present | present | present | present | present | present | present | present | present | present | present | present | present | present | present | present | present | present | present | present | present | present | present | present | present |
| *kdgK* | present | present | present | present | present | present | present | present | present | present | present | present | present | present | present | present | present | present | present | present | absent | present | present | present | present | present | present | absent | present | present | present | absent | present | present | present | present | present | present | present |
| *lapA* | present | present | present | present | present | present | present | present | present | present | present | present | present | present | present | present | present | present | present | present | present | present | present | present | present | present | present | present | present | present | present | present | present | present | present | present | present | present | present |
| *lepB* | present | present | present | present | present | present | present | present | present | present | present | present | present | present | present | present | present | present | present | present | present | present | present | present | present | present | present | present | present | present | present | present | present | present | present | present | present | present | present |
| *lldP* | present | present | present | present | present | present | present | present | present | present | present | present | present | present | present | present | present | present | present | present | present | present | present | present | present | present | present | present | present | present | present | present | present | present | present | present | present | present | present |
| *lolB* | present | present | present | present | present | present | present | present | present | present | present | present | present | present | present | present | present | present | present | present | present | present | present | present | present | present | present | present | present | present | present | present | present | present | present | present | present | present | present |
| *lst* | present | present | present | present | present | present | present | present | present | present | present | present | present | present | present | present | present | present | present | present | present | present | present | present | present | present | present | present | present | present | present | present | present | present | present | present | present | present | present |
| *mdh* | present | present | present | present | present | present | present | present | present | present | present | present | present | present | present | present | present | present | present | present | present | present | present | present | present | present | present | present | present | present | present | present | present | present | present | present | present | present | present |
| *menE* | present | present | present | present | present | present | present | present | present | present | present | present | present | present | present | present | present | present | present | present | present | present | present | present | present | present | present | present | present | present | present | present | present | present | present | present | present | present | present |
| *mepA* | present | present | present | present | present | present | present | present | present | present | present | present | present | present | present | present | present | present | present | present | present | present | present | present | present | present | present | present | present | present | present | present | present | present | present | present | present | present | present |
| *mglA* | present | present | present | present | present | present | present | present | present | present | present | present | present | present | present | present | present | present | present | present | present | present | present | present | present | present | present | present | present | present | present | present | present | present | present | present | present | present | present |
| *mlaA* | present | present | present | present | present | present | present | present | present | present | present | present | present | present | present | present | present | present | present | present | present | present | present | present | present | present | present | present | present | present | present | present | present | present | present | present | present | present | present |
| *mlaC* | present | present | present | present | present | present | present | present | present | present | present | present | present | present | present | present | present | present | present | present | present | present | present | present | present | present | present | present | present | present | present | present | present | present | present | present | present | present | present |
| *mntH* | present | present | present | present | present | present | present | present | present | present | present | present | present | present | present | present | present | present | present | present | present | present | present | present | present | present | present | present | present | present | absent | absent | present | present | present | present | present | present | present |
| *moaC* | present | present | present | present | present | present | present | present | present | present | present | present | present | present | present | present | present | present | present | present | present | present | present | present | present | present | present | present | present | present | present | present | present | present | present | present | present | present | present |
| *modA* | present | present | present | present | present | present | present | present | present | present | present | present | present | present | present | present | present | present | present | present | present | present | present | present | present | present | present | present | present | present | present | present | present | present | present | present | present | present | present |
| *mog* | present | present | present | present | present | present | present | present | present | present | present | present | present | present | present | present | present | present | present | present | present | present | present | present | present | present | present | present | present | present | present | present | present | present | present | present | present | present | present |
| *mpl* | present | present | present | present | present | present | present | present | present | present | present | present | present | present | present | present | present | present | present | present | present | present | present | absent | present | present | present | present | present | present | present | present | present | present | present | present | present | present | present |
| *msbA_1* | present | present | present* | present | present* | present | present* | present* | present | present* | present | present | present* | present* | present | present* | present* | present | present* | present | present* | present* | present* | present* | present* | present | present* | present* | present | present | present | present* | present | present | present* | present | present | present | present |
| *mtr* | present* | present* | present* | present | present* | present | present* | present* | present* | present* | present | present* | present* | present* | present* | present* | present* | present | present* | present* | present | present* | present* | present | present | present* | present* | present | present* | present* | present* | present* | present | present* | present | present | present* | present* | present* |
| *murE* | present | present | present | present | present | present | present | present | present | present | present | present | present | present | present | present | present | present | present | present | present | present | present | present | present | present | present | present | present | present | present | present | present | present | present | present | present | present | present |
| *murF* | present | present | present | present | present | present | present | present | present | present | present | present | present | present | present | present | present | present | present | present | present | present | present | present | present | present | present | present | present | present | present | present | present | present | present | present | present | present | present |
| *mutM* | present | present | present | present | present | present | present | present | present | present | present | present | present | present | present | present | present* | present | present* | present* | present | present | present | present* | present* | present | present | present | present | present* | present | present | present | present | present | present | present* | present | present |
| *nagB* | present | present | present | present | present | present | present | present | present | present | present | present | present | present | present | present | present | present | present | present | present | present | present | present | present | present | present | present | present | present | present | present | present | present | present | present | present | present | present |
| *nagK* | present | present | present | present | present | present | present | present | present | present | present | present | present | present | present | present | present | present | present | present | present | present | present | present | present | present | present | present | present | present | present | present | present | present | present | present | present | present | present |
| *nanK* | present | present | present | present | present | present | present | present | present | present | present | present | present | present | present | present | present | present | present | present | present | present | present | present | present | present | present | present | present | present | present | present | present | present | present | present | present | present | present |
| *neuA* | present | present | present | present | present | present | present | present | present | present | present | present | present | present | present | present | present | present | present | present | present | present | present | present | present | present | present | present | present | present | present | present | present | present | present | present | present | present | present |
| *nhaA* | present | present | present | present | present | present | present | present | present | present | present | present | present | present | present | present | present | present | present | present | present | present | present | present | present | present | present | present | present | present | present | present | present | present | present | present | present | present | present |
| *norM* | present | present | present | absent | present | present | present | present | present | present | present | present | present | present | present | present | present | present | present | present | present | present | present | present | present | present | present | present | absent | present | present | present | present | present | present | present | present | present | present |
| *ompP1* | present | present | present | present | present | present | present | present | present | present | present | present | present | absent | present | present | present | present | present | present | present | present | present | present | present | present | present | present | present | present | present | present | present | present | present | present | present | present | present |
| *opgE* | present | present | present | present | present | present | present | present | present | present | present | present | present | present | present | present | present | present | present | present | present | present | present | present | present | present | present | present | present | present | present | present | present | present | present | present | present | present | present |
| *oppA* | present | present | present | present | present | present | present | present | present | present | present | present | present | present | present | present | present | present | present | present | present | present | present | present | present | present | present | present | present | present | present | present | present | present | present | present | present | present | present |
| *patB* | present | present | present | absent | present | present | present | present | present | present | present | present | present | present | present | present | present | present | present | present | present | present | present | present | present | present | present | absent | present | present | present | present | present | present | present | absent | present | present | present |
| *pcp* | present | present | present | present | present | present | present | present | present | present | present | present | present | present | present | present | present | present | present | present | present | present | present | present | present | present | present | present | present | present | present | present | present | present | present | present | present | present | present |
| *pe* | present | present | present | present | present | present | present | present | present | present | present | present | present | present | present | present | present | absent | present | present | present | present | present | present | present | present | present | present | present | present | present | present | absent | present | present | present | present | present | present |
| *pepB* | present | present | present | present | present | present | present | present | present | present | present | present | present | present | present | present | present | present | present | present | present | present | present | present | present | present | present | present | present | present | present | present | present | present | present | present | present | present | present |
| *pepT* | present | present | present | present | present | present | present | present | present | present | absent | present | present | present | present | present | present | present | present | present | present | present | present | present | present | present | present | present | present | present | present | present | present | present | present | present | present | present | present |
| *phoR* | present | present | present | present | present | present | present | present | present | present | present | present | present | present | present | present | present | present | present | present | present | present | present | present | present | present | present | absent | present | present | present | present | present | present | present | present | present | present | present |
| *pldB* | present | present | present | present | present | present | present | present | present | present | present | present | present | present | present | present | present | present | present | present | present | present | present | present | present | present | present | present | present | present | present | present | present | present | present | present | present | present | present |
| *plsY* | present | present | present | present | present | present | present | present | present | present | present | present | present | present | present | present | present | present | present | present | present | present | present | present | present | present | present | present | present | present | present | present | present | present | present | present | present | present | present |
| *potB* | present | present | present | present | present | present | present | present | present | present | present | present | present | present | present | present | present | present | present | present | present | present | present | present | present | present | present | present | present | present | present | present | present | present | present | present | present | present | present |
| *purR* | present | present | present | present | present | present | present | present | present | present | present | present | present | present | present | present | present | present | present | present | present | present | present | present | present | present | present | present | present | present | present | present | present | present | present | present | present | present | present |
| *sixA* | present | present | present | present | present | absent | present | present | present | present | present | present | present | present | present | present | present | present | present | present | present | present | present | present | present | present | present | present | present | present | present | present | present | present | present | present | present | present | present |
| *slp* | present | present | present | present | present | present | present | present | present | present | absent | present | present | present | present | present | present | present | present | present | present | present | present | present | present | present | present | present | present | present | present | present | present | present | present | present | present | present | present |
| *sotB* | present | present | present | present | present | present | present | present | present | present | present | present | present | present | present | present | present | present | present | present | present | present | present | present | present | present | present | present | present | present | present | present | present | present | present | present | present | present | present |
| *ssb_1* | present | present* | present* | present | present | present* | present* | present* | present | present | present* | present | present | present* | present* | present | present | present | present* | present | present* | present | present* | present | present | present* | present* | present* | present* | present* | present* | present* | present | present | present* | present | present* | present | present* |
| *tamB* | present | present | present | present | present | present | present | present | present | present | absent | present | present | present | present | present | present | present | present | present | present | present | present | present | present | present | present | present | present | present | present | present | present | present | present | absent | present | present | present |
| *tbpB* | present | present | present | present | present | present | present | present | present | present | present | present | present | present | present | present | present | present | present | present | present | present | present | present | present | present | present | absent | present | present | present | present | present | present | present | present | present | present | present |
| *tdk* | present | present | present | present | present | present | present | present | present | present | present | present | present | present | present | present | present | present | present | present | present | present | present | present | present | present | present | present | present | present | present | present | present | present | present | present | present | present | present |
| *trpC* | present | present | present | present | present | present | present | present | present | present | present | present | present | present | present | present | present | present | present | present | present | present | present | present | present | present | present | present | present | present | present | present | present | present | present | present | present | present | present |
| *trpE* | present | present | present | present | present | present | present | present | present | present | present | present | present | present | present | present | absent | present | present | present | present | present | present | present | present | present | present | present | present | present | present | present | present | present | present | present | present | present | present |
| *tuaG* | absent | present | present | absent | absent | absent | absent | absent | absent | present | absent | absent | absent | absent | absent | absent | present | present | present | present | present | present | present | present | present | present | present | absent | present | present | present | absent | present | present | present | present | present | present | present |
| *tusB* | present | present | present | present | present | present | present | present | present | present | present | present | present | present | present | present | present | present | present | present | present | present | present | present | present | present | present | present | present | present | present | present | present | present | present | present | present | present | present |
| *tusC* | present | present | present | present | present | present | present | present | present | present | present | present | present | present | present | present | present | present | present | present | present | present | present | present | present | present | present | present | present | present | present | present | present | present | present | present | present | present | present |
| *tusE* | present | present | present | present | present | present | present | present | present | present | present | present | present | present | present | present | present | present | present | present | present | present | present | present | present | present | present | present | present | present | present | present | present | present | present | present | present | present | present |
| *tyrP_1* | present | present | present | present | present | present | present | present | present | present | present | present | present | present | present | present | present | present | present | present | present | present | present | present | present | present | present | present | present | present | present | present | present | present | present | present | present | present | present |
| *ulaE* | present | present | present | absent | present | absent | absent | absent | present | absent | present | present | absent | present | present | absent | present | absent | absent | absent | present | absent | present | present | present | absent | absent | absent | absent | present | present | absent | absent | present | absent | present | absent | present | present |
| *ung* | present | present | present | present | present | present | present | present | present | present | present | present | present | present | present | present | present | present | present | present | present | present | present | present | present | present | present | present | present | present | present | present | present | present | present | present | present | present | present |
| *yejM* | present | present | present | present | present | present | present | present | present | present | present | present | present | present | present | present | present | present | present | present | present | present | present | present | present | present | present | present | present | present | present | present | present | present | present | present | present | present | present |
| *ytfE* | present | present | present | present | present | present | present | present | present | present | present | present | present | present | present | present | present | present | present | present | present | present | present | present | present | present | present | present | present | present | present | present | present | present | present | present | present | present | present |
| *znuA* | present | present | present | present | present | present | present | present | present | present | present | present | present | present | present | present | present | present | present | present | present | present | present | absent | present | present | present | present | present | present | present | present | present | present | present | present | present | present | present |

*multiple copies

**different gene (different sequence) with same annotation

**TableS6.Codon position of all SNPs identified in the 805 polymorphic genes.**

| Gene | Type | Polymorphism | Substitution | Position in codon |
|---|---|---|---|---|
| *adk* | Nonsynonymous | C640G | G214R | 1 |
| *adk* | Nonsynonymous | C641A | G214V | 2 |
| *adk* | Synonymous | G642T | G214G | NA |
| *adk* | Nonsynonymous | T643C | *215Q | 1 |
| *adk* | Nonsynonymous | T644A | *215L | 2 |
| *argG* | Nonsynonymous | T1321A | L441I | 1 |
| *aroA* | Nonsynonymous | T1294C | N432D | 1 |
| *aroA* | Nonsynonymous | T1295C | N432S | 2 |
| *aroA* | Synonymous | A1296G | N432N | NA |
| *aroA* | Nonsynonymous | A1297G | *433Q | 1 |
| *aroA* | Nonsynonymous | A1298C | *433S | 2 |
| *artP* | Nonsynonymous | T109A | S37C | 1 |
| *artP* | Nonsynonymous | C110G | S37T | 2 |
| *artP* | Nonsynonymous | T139A | T47S | 1 |
| *artP* | Nonsynonymous | G140C | T47R | 2 |
| *aspA* | Nonsynonymous | A1346C | E449A | 2 |
| *atsB* | Nonsynonymous | T218A | Y73F | 2 |
| *atsB* | Nonsynonymous | T225A | R75S | 3 |
| *atsB* | Nonsynonymous | T240G | *80Y | 3 |
| *bacA* | Nonsynonymous | C36G | N12K | 3 |
| *bacA* | Nonsynonymous | A40T | I14F | 1 |
| *bacA* | Nonsynonymous | T40A | I14F | 1 |
| *bacA* | Nonsynonymous | T56A | F19Y | 2 |
| *bacA* | Nonsynonymous | T57A | F19L | 3 |
| *bacA* | Nonsynonymous | T58G | W20G | 1 |
| *bacA* | Nonsynonymous | G60T | W20C | 3 |
| *bacA* | Nonsynonymous | G61A | V21I | 1 |
| *bacA* | Nonsynonymous | C518G | T173S | 2 |
| *bacA* | Nonsynonymous | G556A | V186I | 1 |
| *bamA* | Nonsynonymous | C1982G | G661A | 2 |
| *bamA* | Nonsynonymous | T2322A | K774N | 3 |
| *bamA* | Nonsynonymous | A2323T | Y775N | 1 |
| *bcp* | Nonsynonymous | T469C | Y157H | 1 |
| *bioD1* | Nonsynonymous | T706G | M236L | 1 |
| *bioD1* | Nonsynonymous | A707T | M236K | 2 |
| *bioD1* | Nonsynonymous | C708T | M236I | 3 |
| *bioD1* | Nonsynonymous | T710G | Q237P | 2 |
| *bioD1* | Nonsynonymous | A715G | *239Q | 1 |
| *bioD1* | Nonsynonymous | T724C | K242E | 1 |
| *bioD1* | Nonsynonymous | A727C | *243E | 1 |
| *bioD1* | Nonsynonymous | T727G | *243E | 1 |
| *bolA* | Nonsynonymous | A251T | E84V | 2 |

| | | | | |
|---|---|---|---|---|
| *bolA* | Nonsynonymous | A269T | E90V | 2 |
| *bolA* | Synonymous | A270G | E90E | NA |
| *bolA* | Nonsynonymous | C272T | T91I | 2 |
| *bspRIM* | Nonsynonymous | T660A | K220N | 3 |
| *bspRIM* | Nonsynonymous | C661G | A221P | 1 |
| *bspRIM* | Nonsynonymous | G662C | A221G | 2 |
| *bspRIM* | Nonsynonymous | A664C | L222V | 1 |
| *bspRIM* | Nonsynonymous | A665G | L222S | 2 |
| *bspRIM* | Nonsynonymous | A687T | N229K | 3 |
| *bspRIM* | Nonsynonymous | T690G | I230M | 3 |
| *bspRIM* | Synonymous | T693A | A231A | NA |
| *bspRIM* | Nonsense | G694T | E232* | 1 |
| *bspRIM* | Nonsynonymous | A700C | L234V | 1 |
| *bspRIM* | Nonsynonymous | A701G | L234S | 2 |
| *bspRIM* | Nonsynonymous | A723T | N241K | 3 |
| *bspRIM* | Nonsynonymous | A726C | I242M | 3 |
| *can* | Synonymous | A684G | N228N | NA |
| *can* | Nonsynonymous | T685G | T229P | 1 |
| *can* | Nonsynonymous | G686A | T229I | 2 |
| *can* | Nonsynonymous | A688G | *230Q | 1 |
| *can* | Nonsynonymous | A688T | *230K | 1 |
| *can* | Nonsynonymous | T690A | *230Y | 3 |
| *can* | Nonsense | G704T | S235* | 2 |
| *can* | Nonsynonymous | T707C | D236G | 2 |
| *cbiM* | Nonsynonymous | G130A | A44T | 1 |
| *ccmC* | Nonsynonymous | T361A | T121S | 1 |
| *ccmC* | Nonsynonymous | G731A | T244I | 2 |
| *ccmC* | Synonymous | T732A | T244T | NA |
| *ccmC* | Nonsynonymous | T734A | L245Q | 2 |
| *ccmC* | Synonymous | G735C | L245L | NA |
| *ccmC* | Nonsense | A736T | K246* | 1 |
| *chuR_1* | Nonsynonymous | A225T | R75S | 3 |
| *chuR_1* | Nonsynonymous | A240C | *80Y | 3 |
| *chuR_2* | Nonsynonymous | T218A | Y73F | 2 |
| *chuR_2* | Nonsynonymous | T225A | R75S | 3 |
| *citC* | Nonsynonymous | C1001A | R334L | 2 |
| *citG* | Nonsynonymous | G3T | L1F | 3 |
| *cntO* | Nonsynonymous | A142C | L48V | 1 |
| *cntO* | Nonsynonymous | T144A | L48F | 3 |
| *cntO* | Nonsynonymous | G148C | L50V | 1 |
| *cntO* | Nonsynonymous | T160C | N54D | 1 |
| *cntO* | Nonsynonymous | T166C | T56A | 1 |
| *coaBC* | Nonsynonymous | T785C | L262P | 2 |
| *coaBC* | Nonsynonymous | G787C | E263Q | 1 |
| *copA* | Nonsynonymous | T5C | L2P | 2 |

| | | | | |
|---|---|---|---|---|
| *copA* | Synonymous | C36T | S12S | NA |
| *copA* | Nonsynonymous | T55C | T19A | 1 |
| *copA* | Nonsynonymous | A2165C | F722C | 2 |
| *copA* | Nonsynonymous | G2166T | F722L | 3 |
| *cpxR* | Nonsynonymous | A475T | F159I | 1 |
| *cpxR* | Nonsynonymous | A476T | F159Y | 2 |
| *csdA* | Nonsynonymous | C417A | E139D | 3 |
| *csdE* | Nonsynonymous | G281A | T94I | 2 |
| *csdE* | Nonsynonymous | A284G | V95A | 2 |
| *csdE* | Nonsynonymous | T314C | H105R | 2 |
| *cydD* | Nonsynonymous | T115G | F39V | 1 |
| *dacA* | Nonsynonymous | T1177C | S393P | 1 |
| *dacA* | Nonsense | C1178A | S393* | 2 |
| *dacA* | Nonsynonymous | T1180A | *394K | 1 |
| *dam* | Nonsynonymous | T854C | H285R | 2 |
| *dapD* | Nonsynonymous | T829A | *277K | 1 |
| *dcuD* | Nonsynonymous | A1292G | L431S | 2 |
| *dcuD* | Nonsynonymous | A1294T | *432K | 1 |
| *dmsD* | Nonsynonymous | A157T | Y53N | 1 |
| *dmsD* | Nonsynonymous | C181T | E61K | 1 |
| *dtd* | Nonsynonymous | A327T | F109L | 3 |
| *dtd* | Nonsynonymous | T348A | K116N | 3 |
| *dtd* | Nonsynonymous | T351A | L117F | 3 |
| *dtd* | Nonsynonymous | G352C | P118A | 1 |
| *dtd* | Nonsynonymous | G353A | P118L | 2 |
| *envC* | Nonsynonymous | A28T | T10S | 1 |
| *era* | Synonymous | G54A | R18R | NA |
| *era* | Nonsynonymous | T896G | Y299S | 2 |
| *era* | Synonymous | A897G | Y299Y | NA |
| *era* | Synonymous | T897C | Y299Y | NA |
| *era* | Nonsynonymous | A899T | M300K | 2 |
| *era* | Nonsynonymous | G901A | D301N | 1 |
| *esiB* | Synonymous | G9C | L3L | NA |
| *esiB* | Synonymous | G9C | L3L | NA |
| *esiB* | Synonymous | G9C | L3L | NA |
| *esiB* | Nonsynonymous | T14C | K5R | 2 |
| *esiB* | Nonsynonymous | T24A | L8F | 3 |
| *esiB* | Nonsynonymous | A24T | L8F | 3 |
| *esiB* | Nonsynonymous | T39G | L13F | 3 |
| *esiB* | Nonsynonymous | C180A | E60D | 3 |
| *esiB* | Nonsynonymous | A193C | F65V | 1 |
| *esiB* | Nonsynonymous | C520A | D174Y | 1 |
| *esiB* | Synonymous | T561C | Q187Q | NA |
| *esiB* | Nonsynonymous | G565A | H189Y | 1 |
| *esiB* | Synonymous | T570C | A190A | NA |

| | | | | |
|---|---|---|---|---|
| *esiB* | Nonsynonymous | T571C | K191E | 1 |
| *esiB* | Nonsynonymous | C573A | K191N | 3 |
| *esiB* | Nonsynonymous | T581A | Y194F | 2 |
| *esiB* | Synonymous | A591G | G197G | NA |
| *esiB* | Nonsynonymous | C601A | A201S | 1 |
| *esiB* | Nonsynonymous | G602T | A201D | 2 |
| *esiB* | Synonymous | A606G | N202N | NA |
| *esiB* | Synonymous | C606T | Q202Q | NA |
| *esiB* | Synonymous | T609C | G203G | NA |
| *esiB* | Synonymous | C609T | G203G | NA |
| *esiB* | Nonsynonymous | T610G | N204H | 1 |
| *esiB* | Nonsense | T610A | K204* | 1 |
| *esiB* | Nonsynonymous | C611T | R204H | 2 |
| *esiB* | Nonsynonymous | C612A | K204N | 3 |
| *esiB* | Synonymous | G615T | G205G | NA |
| *esiB* | Nonsynonymous | T616C | K206E | 1 |
| *esiB* | Nonsynonymous | C618A | K206N | 3 |
| *esiB* | Nonsynonymous | A618T | N206K | 3 |
| *esiB* | Nonsynonymous | T626C | D209G | 2 |
| *esiB* | Nonsynonymous | A628C | Y210D | 1 |
| *esiB* | Nonsynonymous | T628G | S210R | 1 |
| *esiB* | Nonsynonymous | C629T | S210N | 2 |
| *esiB* | Nonsynonymous | T637C | I213V | 1 |
| *esiB* | Nonsynonymous | T649C | S217G | 1 |
| *esiB* | Synonymous | C654T | G218G | NA |
| *esiB* | Synonymous | G879T | V293V | NA |
| *esiB* | Nonsynonymous | C892A | V298F | 1 |
| *esiB* | Synonymous | A900T | A300A | NA |
| *esiB* | Nonsynonymous | G906T | N302K | 3 |
| *esiB* | Nonsynonymous | C917T | R306K | 2 |
| *esiB* | Synonymous | T921C | A307A | NA |
| *fbp* | Nonsynonymous | G3A | M1I | 3 |
| *fbp* | Nonsynonymous | G514T | H172N | 1 |
| *fbp* | Nonsynonymous | T691G | N231H | 1 |
| *fbp* | Nonsynonymous | G704A | A235V | 2 |
| *fbp* | Nonsynonymous | C898G | E300Q | 1 |
| *fbp* | Nonsynonymous | G908T | A303E | 2 |
| *fbp* | Nonsynonymous | C916G | E306Q | 1 |
| *fbp* | Nonsynonymous | G923A | A308V | 2 |
| *fbpC2* | Synonymous | G39C | T13T | NA |
| *fbpC2* | Synonymous | A42T | V14V | NA |
| *fbpC2* | Synonymous | T42A | V14V | NA |
| *fbpC2* | Nonsynonymous | T63A | F21L | 3 |
| *fbpC2* | Nonsynonymous | A64C | N22H | 1 |
| *fbpC2* | Nonsynonymous | A65C | N22T | 2 |

| | | | | |
|---|---|---|---|---|
| *fbpC2* | Nonsynonymous | T66G | N22K | 3 |
| *fbpC2* | Nonsense | C887A | S296* | 2 |
| *fbpC2* | Nonsynonymous | C895T | P299S | 1 |
| *fbpC2* | Nonsynonymous | T911G | N304T | 2 |
| *fhuD* | Nonsynonymous | T8A | K3I | 2 |
| *fnr* | Nonsynonymous | T746G | N249T | 2 |
| *fnr* | Nonsynonymous | T756G | K252N | 3 |
| *fnr* | Nonsynonymous | T767A | V256D | 2 |
| *ftsH_1* | Nonsynonymous | G1870A | E624K | 1 |
| *ftsH_1* | Nonsynonymous | A1892C | D631A | 2 |
| *ftsH_1* | Nonsynonymous | A1894G | N632D | 1 |
| *ftsH_1* | Nonsynonymous | A1895T | N632I | 2 |
| *ftsH_2* | Synonymous | C1020T | A340A | NA |
| *ftsH_2* | Synonymous | A1245T | A415A | NA |
| *ftsH_2* | Synonymous | A1251G | H417H | NA |
| *ftsH_2* | Synonymous | C1254G | A418A | NA |
| *ftsH_2* | Synonymous | C1263A | G421G | NA |
| *ftsH_2* | Synonymous | T1314A | R438R | NA |
| *ftsH_2* | Synonymous | T1317A | G439G | NA |
| *ftsH_2* | Synonymous | C1323G | A441A | NA |
| *ftsH_2* | Synonymous | T1458C | T486T | NA |
| *ftsH_2* | Synonymous | A1620G | T540T | NA |
| *ftsH_2* | Synonymous | T1623A | A541A | NA |
| *ftsH_2* | Nonsynonymous | A1627C | S543A | 1 |
| *ftsH_2* | Synonymous | C1629T | S543S | NA |
| *ftsH_2* | Synonymous | A1662G | R554R | NA |
| *ftsH_2* | Nonsynonymous | G1870A | E624K | 1 |
| *ftsH_2* | Nonsynonymous | A1892C | D631A | 2 |
| *ftsI* | Nonsynonymous | G166T | G56C | 1 |
| *ftsI* | Nonsynonymous | G224A | S75N | 2 |
| *ftsI* | Nonsynonymous | C364A | H122N | 1 |
| *ftsI* | Nonsynonymous | T374C | V125A | 2 |
| *ftsI* | Nonsynonymous | T391G | S131A | 1 |
| *ftsI* | Nonsynonymous | C403G | Q135E | 1 |
| *ftsI* | Nonsynonymous | G454T | A152S | 1 |
| *ftsI* | Nonsynonymous | C494T | S165L | 2 |
| *ftsI* | Nonsynonymous | A497G | N166S | 2 |
| *ftsI* | Nonsynonymous | G1578T | K526N | 3 |
| *ftsI* | Nonsynonymous | G1652A | G551D | 2 |
| *ftsN* | Nonsynonymous | A41C | N14T | 2 |
| *glmM* | Nonsynonymous | A1T | M1L | 1 |
| *glmM* | Nonsynonymous | T2C | M1T | 2 |
| *glnE* | Synonymous | G1362A | D454D | NA |
| *glnE* | Nonsynonymous | C1363T | D455N | 1 |
| *glnE* | Nonsynonymous | G1380T | L460F | 3 |

| | | | | |
|---|---|---|---|---|
| *glpQ* | Nonsynonymous | T238G | C80G | 1 |
| *glpQ* | Nonsynonymous | G1066T | G356C | 1 |
| *glpQ* | Nonsynonymous | T1094G | *365S | 2 |
| *gltC* | Nonsynonymous | C4T | E2K | 1 |
| *gmhA_2* | Nonsynonymous | C471A | K157N | 3 |
| *gor* | Nonsynonymous | G310T | A104S | 1 |
| *gor* | Nonsynonymous | T321A | N107K | 3 |
| *gpt_1* | Nonsynonymous | T53C | K18R | 2 |
| *gpt_2* | Synonymous | G21A | V7V | NA |
| *gpt_2* | Synonymous | G51T | R17R | NA |
| *gpt_2* | Nonsynonymous | C53T | R18K | 2 |
| *gpt_2* | Synonymous | T135C | V45V | NA |
| *gpt_2* | Synonymous | A186G | S62S | NA |
| *gpt_2* | Synonymous | T207C | Q69Q | NA |
| *gpt_2* | Synonymous | A210T | G70G | NA |
| *gpt_2* | Synonymous | A214G | L72L | NA |
| *gpt_2* | Synonymous | A225C | L75L | NA |
| *gpt_2* | Synonymous | C447A | V149V | NA |
| *gpt_2* | Nonsynonymous | A468T | *156Y | 3 |
| *gpt_2* | Nonsynonymous | T469C | F157L | 1 |
| *greA* | Synonymous | C465T | V155V | NA |
| *greA* | Nonsynonymous | T467C | N156S | 2 |
| *greA* | Nonsynonymous | A469T | Y157N | 1 |
| *greA* | Nonsynonymous | T470G | Y157S | 2 |
| *greA* | Synonymous | A471G | Y157Y | NA |
| *greA* | Nonsense | A471T | Y157* | 3 |
| *greA* | Nonsynonymous | A473G | I158T | 2 |
| *greA* | Nonsynonymous | T473A | I158N | 2 |
| *greA* | Synonymous | T474A | I158I | NA |
| *greA* | Nonsynonymous | A475C | *159E | 1 |
| *greA* | Nonsynonymous | T477A | *159Y | 3 |
| *greA* | Nonsense | C493A | G165* | 1 |
| *hap* | Nonsynonymous | C3203T | P1068L | 2 |
| *hindIIIM* | Nonsynonymous | A53T | D18V | 2 |
| *hindIIIR* | Nonsynonymous | T632A | N211I | 2 |
| *hindIIIR* | Nonsynonymous | T710G | K237T | 2 |
| *hindIIIR* | Nonsynonymous | A733G | R245G | 1 |
| *hindIIIR* | Nonsynonymous | C734A | R245I | 2 |
| *hsdR* | Nonsynonymous | T566G | F189C | 2 |
| *hsdR* | Nonsynonymous | A568T | N190Y | 1 |
| *hsdR* | Nonsynonymous | A590G | K197R | 2 |
| *hsdR* | Nonsynonymous | A590T | K197I | 2 |
| *hsdR* | Nonsynonymous | T590A | K197I | 2 |
| *hsdR* | Synonymous | T591C | K197K | NA |
| *hsdR* | Nonsynonymous | A592T | Y198N | 1 |

| | | | | |
|---|---|---|---|---|
| *hxuC* | Nonsynonymous | A1673G | N558S | 2 |
| *hxuC* | Nonsynonymous | A1678G | S560G | 1 |
| *hxuC* | Nonsynonymous | A1684G | S562G | 1 |
| *iaaH* | Nonsynonymous | T5C | N2S | 2 |
| *iaaH* | Nonsynonymous | T7A | L3I | 1 |
| *iaaH* | Nonsense | T8G | L3* | 2 |
| *iga* | Nonsynonymous | A1012T | S338T | 1 |
| *iga* | Nonsynonymous | G1013T | S338Y | 2 |
| *iga* | Nonsynonymous | G1015C | Q339E | 1 |
| *ihfB* | Nonsynonymous | G244A | V82M | 1 |
| *kdgK* | Synonymous | T423C | L141L | NA |
| *kdgK* | Synonymous | T423C | L141L | NA |
| *kdgK* | Nonsynonymous | T423G | L141F | 3 |
| *kdgK* | Nonsynonymous | T427C | K143E | 1 |
| *kdgK* | Nonsynonymous | T430A | N144Y | 1 |
| *kdgK* | Nonsynonymous | T431A | N144I | 2 |
| *kdgK* | Nonsynonymous | T431G | N144T | 2 |
| *kdgK* | Nonsense | C451A | E151* | 1 |
| *kdgK* | Nonsynonymous | T455G | Q152P | 2 |
| *kdgK* | Nonsynonymous | T456A | Q152H | 3 |
| *kdgK* | Synonymous | A457G | L153L | NA |
| *kdgK* | Nonsynonymous | T459A | L153F | 3 |
| *lapA* | Nonsynonymous | T50G | V17G | 2 |
| *lapA* | Nonsynonymous | G59A | T20I | 2 |
| *lapA* | Nonsynonymous | T61C | I21V | 1 |
| *lapA* | Nonsynonymous | A62C | I21S | 2 |
| *lapA* | Synonymous | A63G | I21I | NA |
| *lepB* | Nonsense | T98G | L33* | 2 |
| *lepB* | Synonymous | A99G | L33L | NA |
| *lepB* | Nonsynonymous | G119A | S40L | 2 |
| *lepB* | Synonymous | G123A | G41G | NA |
| *lepB* | Synonymous | G123A | G41G | NA |
| *lepB* | Nonsynonymous | C124A | V42L | 1 |
| *lldP* | Synonymous | G102A | A34A | NA |
| *lldP* | Synonymous | T120C | V40V | NA |
| *lldP* | Synonymous | G186A | A62A | NA |
| *lldP* | Synonymous | T252C | G84G | NA |
| *lldP* | Synonymous | C277T | L93L | NA |
| *lldP* | Synonymous | T336C | A112A | NA |
| *lldP* | Synonymous | A351T | G117G | NA |
| *lldP* | Synonymous | G375A | A125A | NA |
| *lldP* | Synonymous | T441G | L147L | NA |
| *lldP* | Synonymous | C648T | I216I | NA |
| *lldP* | Synonymous | A663G | L221L | NA |
| *lldP* | Synonymous | T717A | S239S | NA |

| | | | | |
|---|---|---|---|---|
| *lldP* | Synonymous | T750C | S250S | NA |
| *lldP* | Synonymous | T762G | A254A | NA |
| *lldP* | Synonymous | A768C | R256R | NA |
| *lldP* | Synonymous | T975C | S325S | NA |
| *lldP* | Synonymous | A1047G | V349V | NA |
| *lldP* | Synonymous | A1077G | V359V | NA |
| *lldP* | Synonymous | G1086A | A362A | NA |
| *lldP* | Synonymous | C1107A | S369S | NA |
| *lldP* | Synonymous | T1239A | I413I | NA |
| *lldP* | Synonymous | C1243A | R415R | NA |
| *lldP* | Synonymous | T1299C | G433G | NA |
| *lldP* | Synonymous | T1389A | I463I | NA |
| *lldP* | Synonymous | T1404C | V468V | NA |
| *lldP* | Synonymous | A1413G | L471L | NA |
| *lldP* | Synonymous | C1422T | V474V | NA |
| *lldP* | Synonymous | A1428G | G476G | NA |
| *lldP* | Synonymous | A1476T | S492S | NA |
| *lldP* | Synonymous | T1530G | P510P | NA |
| *lldP* | Synonymous | C1548T | I516I | NA |
| *lldP* | Nonsynonymous | A1550G | I517T | 2 |
| *lldP* | Synonymous | T1567C | L523L | NA |
| *lldP* | Nonsynonymous | G1594T | L532I | 1 |
| *lolB* | Nonsynonymous | G28A | L10F | 1 |
| *lolB* | Nonsynonymous | A37G | T13A | 1 |
| *lolB* | Nonsynonymous | G46A | A16T | 1 |
| *lolB* | Nonsynonymous | T46A | T16S | 1 |
| *lolB* | Synonymous | A48C | T16T | NA |
| *lolB* | Synonymous | A51T | A17A | NA |
| *lolB* | Synonymous | T84A | R28R | NA |
| *lolB* | Synonymous | T90A | T30T | NA |
| *lolB* | Synonymous | C96T | V32V | NA |
| *lolB* | Synonymous | A102G | Y34Y | NA |
| *lolB* | Synonymous | T114C | T38T | NA |
| *lolB* | Synonymous | C129T | Q43Q | NA |
| *lolB* | Synonymous | C132T | Q44Q | NA |
| *lolB* | Synonymous | A171G | A57A | NA |
| *lolB* | Synonymous | C255T | T85T | NA |
| *lolB* | Synonymous | A291C | L97L | NA |
| *lolB* | Synonymous | G306A | H102H | NA |
| *lolB* | Synonymous | G336A | N112N | NA |
| *lolB* | Synonymous | A370G | L124L | NA |
| *lolB* | Synonymous | T390G | G130G | NA |
| *lolB* | Synonymous | G424A | L142L | NA |
| *lolB* | Synonymous | A447G | N149N | NA |
| *lolB* | Synonymous | T450C | A150A | NA |

| | | | | |
|---|---|---|---|---|
| *lolB* | Synonymous | G564A | N188N | NA |
| *lolB* | Synonymous | A598G | L200L | NA |
| *lolB* | Synonymous | A615G | D205D | NA |
| *lst* | Nonsynonymous | T410A | I137N | 2 |
| *lst* | Nonsynonymous | T456A | K152N | 3 |
| *lst* | Nonsynonymous | A460T | I154L | 1 |
| *lst* | Nonsynonymous | A468C | K156N | 3 |
| *lst* | Nonsynonymous | C469A | H157N | 1 |
| *lst* | Nonsynonymous | A475C | T159P | 1 |
| *lst* | Nonsynonymous | C476G | T159R | 2 |
| *lst* | Synonymous | T486G | P162P | NA |
| *lst* | Nonsynonymous | A491G | Y164C | 2 |
| *lst* | Nonsynonymous | A491T | Y164F | 2 |
| *lst* | Nonsynonymous | T494G | K165T | 2 |
| *lst* | Nonsynonymous | A499T | I167F | 1 |
| *lst* | Synonymous | C564T | E188E | NA |
| *lst* | Synonymous | G567A | V189V | NA |
| *lst* | Synonymous | A579T | I193I | NA |
| *lst* | Nonsynonymous | T594A | L198F | 3 |
| *lst* | Nonsynonymous | T613C | I205V | 1 |
| *lst* | Synonymous | G615T | I205I | NA |
| *lst* | Synonymous | T624A | I208I | NA |
| *lst* | Nonsynonymous | A646G | S216P | 1 |
| *lst* | Synonymous | G690A | V230V | NA |
| *lst* | Nonsynonymous | T898A | I300L | 1 |
| *lst* | Nonsynonymous | A899G | I300T | 2 |
| *lst* | Nonsynonymous | A901T | Y301N | 1 |
| *lst* | Nonsense | A903T | Y301* | 3 |
| *mdh* | Nonsynonymous | T918G | E306D | 3 |
| *mdh* | Nonsense | T919A | K307* | 1 |
| *menE* | Nonsynonymous | G1196A | S399N | 2 |
| *menE* | Nonsynonymous | T1203A | S401R | 3 |
| *menE* | Nonsynonymous | G1221T | M407I | 3 |
| *menE* | Nonsynonymous | T1222G | F408V | 1 |
| *mepA* | Nonsynonymous | T25G | T9P | 1 |
| *mepA* | Nonsynonymous | C26G | T9S | 2 |
| *mepA* | Synonymous | C48T | F16F | NA |
| *mepA* | Nonsynonymous | C50A | S17Y | 2 |
| *mepA* | Nonsynonymous | G52T | L18M | 1 |
| *mepA* | Synonymous | C54A | L18L | NA |
| *mglA* | Synonymous | T636C | I212I | NA |
| *mglA* | Nonsynonymous | T654A | D218E | 3 |
| *mglA* | Nonsynonymous | G655A | V219M | 1 |
| *mglA* | Synonymous | G657T | V219V | NA |
| *mglA* | Synonymous | T660G | T220T | NA |

| | | | | |
|---|---|---|---|---|
| *mglA* | Nonsynonymous | A661G | I221V | 1 |
| *mlaA* | Nonsynonymous | C14A | A5E | 2 |
| *mlaA* | Synonymous | T36G | G12G | NA |
| *mlaA* | Synonymous | A36C | G12G | NA |
| *mlaA* | Nonsynonymous | G37T | A13S | 1 |
| *mlaA* | Synonymous | C39A | A13A | NA |
| *mlaA* | Nonsynonymous | A752T | *251L | 2 |
| *mlaA* | Synonymous | A753G | *251* | NA |
| *mlaA* | Nonsynonymous | T754A | *252K | 1 |
| *mlaC* | Nonsynonymous | G169C | D57H | 1 |
| *mntH* | Nonsynonymous | A1189T | S397C | 1 |
| *moaC* | Nonsynonymous | A1T | M1L | 1 |
| *moaC* | Nonsynonymous | G448A | G150R | 1 |
| *modA* | Nonsynonymous | A13C | T5P | 1 |
| *modA* | Nonsynonymous | C14T | T5I | 2 |
| *modA* | Nonsynonymous | T19A | I7F | 1 |
| *modA* | Nonsynonymous | G23A | S8L | 2 |
| *modA* | Synonymous | A24T | S8S | NA |
| *modA* | Nonsynonymous | T35C | L12S | 2 |
| *modA* | Nonsynonymous | T36A | L12F | 3 |
| *modA* | Nonsynonymous | A38C | I13S | 2 |
| *modA* | Synonymous | A39G | I13I | NA |
| *modA* | Nonsynonymous | G39C | I13M | 3 |
| *modA* | Nonsynonymous | C40T | A14T | 1 |
| *modA* | Nonsynonymous | G41T | A14E | 2 |
| *modA* | Synonymous | G42A | A14A | NA |
| *modA* | Nonsynonymous | G43A | G15R | 1 |
| *mog* | Nonsynonymous | T1A | M1L | 1 |
| *mog* | Nonsynonymous | A19T | I7F | 1 |
| *mog* | Nonsynonymous | T20A | I7N | 2 |
| *mog* | Synonymous | T21A | I7I | NA |
| *mpl* | Nonsense | T1357A | K453* | 1 |
| *mpl* | Nonsynonymous | T1358G | K453T | 2 |
| *mpl* | Nonsynonymous | T1359G | K453N | 3 |
| *mpl* | Nonsynonymous | A1360T | *454K | 1 |
| *mpl* | Synonymous | T1361C | *454* | NA |
| *mpl* | Nonsynonymous | A1361C | *454S | 2 |
| *msbA_1* | Nonsynonymous | G35A | A12V | 2 |
| *msbA_1* | Synonymous | A57G | G19G | NA |
| *msbA_1* | Nonsynonymous | A115C | F39V | 1 |
| *msbA_1* | Nonsynonymous | G119C | S40C | 2 |
| *msbA_1* | Synonymous | C867G | P289P | NA |
| *mtr* | Nonsynonymous | C914T | T305I | 2 |
| *murE* | Nonsynonymous | T2C | M1T | 2 |
| *murE* | Synonymous | C12A | L4L | NA |

| | | | | |
|---|---|---|---|---|
| *murE* | Synonymous | C78T | S26S | NA |
| *murE* | Synonymous | G84A | K28K | NA |
| *murE* | Nonsynonymous | A221C | E74A | 2 |
| *murE* | Synonymous | T228C | I76I | NA |
| *murE* | Nonsynonymous | C253T | H85Y | 1 |
| *murE* | Synonymous | G276A | S92S | NA |
| *murE* | Synonymous | C279T | S93S | NA |
| *murE* | Synonymous | A282G | L94L | NA |
| *murE* | Synonymous | T285G | A95A | NA |
| *murE* | Synonymous | T330C | G110G | NA |
| *murE* | Synonymous | G342A | T114T | NA |
| *murE* | Synonymous | T375G | A125A | NA |
| *murE* | Nonsynonymous | T383C | V128A | 2 |
| *murE* | Synonymous | G432A | G144G | NA |
| *murE* | Synonymous | A438G | L146L | NA |
| *murE* | Synonymous | A441G | G147G | NA |
| *murE* | Synonymous | G468A | T156T | NA |
| *murE* | Synonymous | G474A | S158S | NA |
| *murE* | Synonymous | A489G | Q163Q | NA |
| *murE* | Synonymous | G492A | S164S | NA |
| *murE* | Synonymous | C495T | S165S | NA |
| *murE* | Nonsynonymous | A503C | N168T | 2 |
| *murE* | Synonymous | G510A | K170K | NA |
| *murE* | Nonsynonymous | A513C | Q171H | 3 |
| *murE* | Synonymous | C519T | G173G | NA |
| *murE* | Nonsynonymous | G529A | A177T | 1 |
| *murE* | Nonsynonymous | T565C | Y189H | 1 |
| *murE* | Nonsynonymous | C572T | A191V | 2 |
| *murE* | Nonsynonymous | C593T | A198V | 2 |
| *murF* | Nonsynonymous | A1195G | I399V | 1 |
| *mutM* | Nonsynonymous | A277C | I93L | 1 |
| *mutM* | Nonsynonymous | T289C | M97V | 1 |
| *mutM* | Synonymous | T435C | K145K | NA |
| *mutM* | Nonsynonymous | T438A | K146N | 3 |
| *mutM* | Nonsynonymous | C443G | R148P | 2 |
| *nagB* | Nonsynonymous | A13G | S5P | 1 |
| *nagB* | Nonsynonymous | A35G | V12A | 2 |
| *nagB* | Nonsynonymous | T67C | T23A | 1 |
| *nagB* | Nonsynonymous | T274A | N92Y | 1 |
| *nagB* | Synonymous | G739A | L247L | NA |
| *nagB* | Nonsynonymous | G788A | A263V | 2 |
| *nagK* | Nonsynonymous | A53G | N18S | 2 |
| *nanK* | Nonsynonymous | T627G | K209N | 3 |
| *neuA* | Nonsense | C674T | W225* | 2 |
| *nhaA* | Nonsynonymous | T1202A | *401L | 2 |

| | | | | |
|---|---|---|---|---|
| *norM* | Nonsynonymous | T16G | L6V | 1 |
| *norM* | Nonsynonymous | T1163G | D388A | 2 |
| *ompP1* | Nonsynonymous | A436G | N146D | 1 |
| *opgE* | Nonsynonymous | T354A | S118R | 3 |
| *opgE* | Nonsynonymous | T1124C | F375S | 2 |
| *oppA* | Nonsynonymous | G29A | R10Q | 2 |
| *patB* | Nonsynonymous | T4A | N2Y | 1 |
| *patB* | Nonsynonymous | T23G | D8A | 2 |
| *patB* | Nonsynonymous | G119C | P40R | 2 |
| *patB* | Nonsynonymous | A148C | F50V | 1 |
| *patB* | Nonsynonymous | A457C | C153G | 1 |
| *patB* | Nonsynonymous | A560G | K187R | 2 |
| *patB* | Nonsynonymous | T773A | M258K | 2 |
| *patB* | Nonsense | A831T | Y277* | 3 |
| *patB* | Nonsynonymous | C991G | E331Q | 1 |
| *pcp* | Nonsynonymous | G367C | E123Q | 1 |
| *pe* | Nonsynonymous | A479C | K160T | 2 |
| *pe* | Nonsynonymous | T479A | K160I | 2 |
| *pe* | Nonsynonymous | T482A | *161L | 2 |
| *pepB* | Nonsynonymous | A89C | H30P | 2 |
| *pepB* | Synonymous | T90C | H30H | NA |
| *pepB* | Synonymous | T90C | H30H | NA |
| *pepB* | Nonsynonymous | T92C | L31S | 2 |
| *pepB* | Nonsynonymous | A93T | L31F | 3 |
| *pepB* | Synonymous | A99G | N33N | NA |
| *pepB* | Nonsynonymous | T104A | E35V | 2 |
| *pepB* | Nonsynonymous | C110T | T37I | 2 |
| *pepB* | Synonymous | T111C | T37T | NA |
| *pepB* | Nonsynonymous | A113C | D38A | 2 |
| *pepB* | Nonsynonymous | G124C | V42L | 1 |
| *pepB* | Nonsynonymous | C125T | T42I | 2 |
| *pepB* | Nonsynonymous | T125G | V42G | 2 |
| *pepB* | Nonsynonymous | A420T | N140K | 3 |
| *pepT* | Nonsynonymous | G19A | K7E | 1 |
| *phoR* | Nonsynonymous | T20A | F7Y | 2 |
| *phoR* | Nonsynonymous | A37C | L13V | 1 |
| *phoR* | Nonsynonymous | C39G | L13F | 3 |
| *phoR* | Nonsynonymous | A279T | K93N | 3 |
| *phoR* | Nonsynonymous | A389C | V130G | 2 |
| *phoR* | Nonsynonymous | C393G | Q131H | 3 |
| *phoR* | Nonsynonymous | A408C | D136E | 3 |
| *phoR* | Nonsynonymous | G421C | E141Q | 1 |
| *phoR* | Nonsynonymous | A422C | E141A | 2 |
| *phoR* | Nonsynonymous | T429A | F143L | 3 |
| *phoR* | Nonsynonymous | A437T | Y146F | 2 |

| | | | | |
|---|---|---|---|---|
| *phoR* | Synonymous | T438C | Y146Y | NA |
| *phoR* | Nonsynonymous | T439G | F147V | 1 |
| *phoR* | Nonsynonymous | C444A | F148L | 3 |
| *phoR* | Nonsynonymous | A445C | S149R | 1 |
| *phoR* | Nonsynonymous | T445A | S149C | 1 |
| *phoR* | Nonsynonymous | C448A | Q150K | 1 |
| *phoR* | Synonymous | A450G | Q150Q | NA |
| *phoR* | Synonymous | A457C | R153R | NA |
| *phoR* | Synonymous | A457C | R153R | NA |
| *phoR* | Nonsynonymous | G458C | R153T | 2 |
| *phoR* | Nonsynonymous | T854A | N285I | 2 |
| *pldB* | Nonsynonymous | A793C | N265H | 1 |
| *pldB* | Nonsense | G793A | Q265* | 1 |
| *pldB* | Nonsynonymous | C795A | N265K | 3 |
| *pldB* | Nonsynonymous | T796G | K266Q | 1 |
| *pldB* | Nonsynonymous | T801A | N267K | 3 |
| *plsY* | Nonsense | A589T | K197* | 1 |
| *plsY* | Nonsynonymous | G594T | K198N | 3 |
| *plsY* | Nonsynonymous | A595C | K199Q | 1 |
| *plsY* | Nonsense | A595T | K199* | 1 |
| *plsY* | Nonsynonymous | A597C | K199N | 3 |
| *potB* | Nonsynonymous | C301A | L101I | 1 |
| *purR* | Nonsynonymous | A1009T | *337K | 1 |
| *sixA* | Nonsynonymous | G46A | D16N | 1 |
| *sixA* | Nonsynonymous | G473T | S158Y | 2 |
| *slp* | Nonsynonymous | C21A | F7L | 3 |
| *sotB* | Synonymous | A1179C | L393L | NA |
| *sotB* | Nonsynonymous | C1183A | Q395K | 1 |
| *sotB* | Nonsynonymous | T1187G | K396T | 2 |
| *sotB* | Nonsynonymous | A1190C | K397T | 2 |
| *ssb_1* | Nonsynonymous | C128T | R43K | 2 |
| *ssb_1* | Nonsynonymous | T263A | Q88L | 2 |
| *ssb_1* | Nonsynonymous | T338C | N113S | 2 |
| *ssb_1* | Nonsynonymous | G359C | T120S | 2 |
| *ssb_1* | Nonsynonymous | T389C | N130S | 2 |
| *ssb_1* | Nonsynonymous | C424T | V142I | 1 |
| *tamB* | Nonsynonymous | A527T | N176I | 2 |
| *tamB* | Synonymous | G543A | V181V | NA |
| *tamB* | Nonsynonymous | T549A | F183L | 3 |
| *tamB* | Nonsynonymous | T549A | L183F | 3 |
| *tamB* | Nonsynonymous | T549A | F183L | 3 |
| *tamB* | Nonsynonymous | T551C | N184S | 2 |
| *tamB* | Nonsynonymous | A552C | N184K | 3 |
| *tamB* | Nonsynonymous | T554C | N185S | 2 |
| *tbpB* | Nonsynonymous | A315C | N105K | 3 |

| | | | | |
|---|---|---|---|---|
| *tbpB* | Nonsynonymous | T323C | K108R | 2 |
| *tbpB* | Synonymous | C327T | T109T | NA |
| *tbpB* | Nonsynonymous | C328T | G110R | 1 |
| *tbpB* | Nonsynonymous | C329T | G110E | 2 |
| *tbpB* | Synonymous | C330A | G110G | NA |
| *tbpB* | Nonsynonymous | G338T | A113D | 2 |
| *tbpB* | Nonsynonymous | G340T | L114I | 1 |
| *tbpB* | Synonymous | G348C | G116G | NA |
| *tbpB* | Nonsynonymous | T427A | I143F | 1 |
| *tbpB* | Nonsynonymous | A1702T | T568S | 1 |
| *tbpB* | Nonsynonymous | C1703T | T568I | 2 |
| *tbpB* | Synonymous | C1704T | T568T | NA |
| *tbpB* | Nonsynonymous | C1705G | Q569E | 1 |
| *tdk* | Nonsynonymous | G574C | Q192E | 1 |
| *tdk* | Nonsynonymous | T577A | I193F | 1 |
| *tdk* | Nonsynonymous | T578A | I193N | 2 |
| *tdk* | Nonsynonymous | A581C | *194S | 2 |
| *tdk* | Nonsynonymous | A583C | I195L | 1 |
| *trpC* | Synonymous | G66A | D22D | NA |
| *trpC* | Synonymous | A171G | G57G | NA |
| *trpC* | Synonymous | A189G | A63A | NA |
| *trpC* | Synonymous | G192A | Y64Y | NA |
| *trpC* | Synonymous | C252T | L84L | NA |
| *trpC* | Synonymous | A261G | I87I | NA |
| *trpC* | Synonymous | C270G | V90V | NA |
| *trpC* | Synonymous | G285C | A95A | NA |
| *trpC* | Synonymous | A288G | S96S | NA |
| *trpC* | Synonymous | C327A | G109G | NA |
| *trpC* | Synonymous | T342C | L114L | NA |
| *trpC* | Synonymous | G396A | I132I | NA |
| *trpC* | Synonymous | T660G | S220S | NA |
| *trpC* | Synonymous | A663G | D221D | NA |
| *trpC* | Synonymous | A666T | R222R | NA |
| *trpC* | Synonymous | G669T | I223I | NA |
| *trpC* | Synonymous | T744C | A248A | NA |
| *trpC* | Synonymous | C801A | V267V | NA |
| *trpC* | Synonymous | C882T | A294A | NA |
| *trpC* | Synonymous | A885G | L295L | NA |
| *trpC* | Nonsynonymous | A1391G | D464G | 2 |
| *trpC* | Nonsense | C1393T | Q465* | 1 |
| *trpC* | Nonsense | G1396T | E466* | 1 |
| *trpC* | Nonsynonymous | A1411C | F471V | 1 |
| *trpC* | Nonsynonymous | C1416A | L472F | 3 |
| *trpC* | Nonsynonymous | T1416G | F472L | 3 |
| *trpC* | Nonsynonymous | A1420T | N474Y | 1 |

| | | | | |
|---|---|---|---|---|
| *trpE* | Nonsynonymous | T257C | Y86C | 2 |
| *trpE* | Nonsense | T280A | K94* | 1 |
| *tuaG* | Nonsense | A454T | K152* | 1 |
| *tuaG* | Nonsynonymous | T461A | I154K | 2 |
| *tusB* | Synonymous | G66A | I22I | NA |
| *tusB* | Synonymous | T87A | V29V | NA |
| *tusB* | Synonymous | G226A | L76L | NA |
| *tusB* | Synonymous | A285T | L95L | NA |
| *tusB* | Synonymous | A285T | L95L | NA |
| *tusB* | Nonsynonymous | T286C | *96Q | 1 |
| *tusB* | Nonsynonymous | A287C | *96S | 2 |
| *tusC* | Nonsynonymous | G293C | C98S | 2 |
| *tusC* | Nonsynonymous | T310A | S104T | 1 |
| *tusC* | Synonymous | A321C | L107L | NA |
| *tusC* | Nonsynonymous | G322A | E108K | 1 |
| *tusC* | Nonsense | A325T | K109* | 1 |
| *tusE* | Nonsynonymous | C325G | L109V | 1 |
| *tusE* | Nonsynonymous | T326A | L109Q | 2 |
| *tusE* | Synonymous | A327T | L109L | NA |
| *tusE* | Nonsynonymous | T328A | *110K | 1 |
| *tyrP_1* | Nonsynonymous | T1201A | *401K | 1 |
| *tyrP_1* | Nonsynonymous | T1201G | *401E | 1 |
| *tyrP_1* | Synonymous | A1202G | *401* | NA |
| *tyrP_1* | Nonsynonymous | A1202T | *401L | 2 |
| *ulaE* | Nonsynonymous | C10T | H4Y | 1 |
| *ulaE* | Nonsynonymous | A11T | H4L | 2 |
| *ulaE* | Nonsynonymous | A13G | K5E | 1 |
| *ulaE* | Nonsynonymous | A14C | K5T | 2 |
| *ung* | Nonsynonymous | T8G | N3T | 2 |
| *yejM* | Nonsynonymous | A1135G | N379D | 1 |
| *yejM* | Nonsynonymous | A1138G | N380D | 1 |
| *yejM* | Nonsynonymous | T1160A | L387H | 2 |
| *yejM* | Nonsynonymous | T1160G | L387R | 2 |
| *yejM* | Nonsynonymous | A1162C | N388H | 1 |
| *yejM* | Nonsynonymous | A1163T | N388I | 2 |
| *yejM* | Nonsynonymous | A1166T | D389V | 2 |
| *yejM* | Nonsynonymous | T1170A | F390L | 3 |
| *ytfE* | Nonsynonymous | T673A | F225I | 1 |
| *znuA* | Nonsynonymous | G47C | S16T | 2 |
